# Programmed cell death-1 receptor mediated regulation of Tbet^+^ NK1.1^−^ Innate Lymphoid Cells within the Tumor Microenvironment

**DOI:** 10.1101/2022.09.21.507469

**Authors:** Jing Xuan Lim, Chester Y. Lai, Grace E. Mallett, David McDonald, Gillian Hulme, Stephanie Laba, Andrew Shapanis, Megan Payne, Warren Patterson, Michael Alexander, Jonathan Coxhead, Andrew Filby, Ruth Plummer, Penny E. Lovat, Giuseppe Sciume, Eugene Healy, Shoba Amarnath

## Abstract

Innate Lymphoid Cells (ILCs) play a key role in tissue mediated immunity and can be controlled by co-receptor signaling. Here we define a subset of ILCs that are Tbet^+^NK1.1^−^ and are present within the tumor microenvironment (TME). We show programmed death-1 receptor (PD-1) expression on ILCs within TME is found in Tbet^+^NK1.1^−^ILCs. PD-1 significantly controlled the proliferation and function of Tbet^+^NK1.1^−^ILCs in multiple murine and human tumors. We found tumor derived lactate enhanced PD-1 expression on Tbet^+^NK1.1^−^ILCs within the TME, which resulted in dampened mTOR signaling along with increased fatty acid uptake. In line with these metabolic changes, PD-1 deficient Tbet^+^NK1.1^−^ILCs expressed significantly increased IFNγ, granzyme B and K. Furthermore, PD1 deficient Tbet^+^NK1.1^−^ ILCs contributed towards diminished tumor growth in an experimental murine model of melanoma. These data demonstrate that PD-1 can regulate anti-tumor responses of Tbet^+^NK1.1^−^ILCs within the tumor microenvironment.

**Highlights:**

1. Tbet^+^NK1.1^−^ ILCs are found in WT and PD1 ko mice
2. PD-1 is expressed on Tbet^+^NK1.1^−^ ILC1s within multiple TME
3. PD-1 controls the proliferation and function of Tbet^+^NK1.1^−^ ILCs within the tumor microenvironment by modulating fatty acid metabolism.
4. PD-1 regulates the proliferation of human Tbet^+^ ILC1s in human cutaneous squamous cell carcinoma (cSCC) and melanoma tumor microenvironment.

## Introduction

Innate lymphoid cells (ILCs) are counterparts of CD4^+^ helper T (Th) cells with diverse phenotypes and functions that mirror their respective Th effector subsets ^1^. ILCs are classified into three groups, with natural killer (NK) cells and lymphoid tissue inducing (LTi) cells being the first to be identified. In recent years, this family of ILCs have expanded and recent nomenclature has classified them into three groups that mirror respective T effector cell lineages ^1^: Group 1 ILCs include NK cells and non-NK cells, constitutively expressing the transcription factor Tbet and respond to type 1 cytokines, namely IL-12, IL-15 and IL-18 by producing IFNγ and TNFα ^2–4^. The expression of Eomes within group 1 ILCs further enable the differentiation between NK cells and helper-like ILC1s ^5, 6^. Mostly, ILC1s are defined as Lin^−^NKp46^+^NK1.1^+^CD49a^+^CD49b^−^ cells while NK cells are identified as subsets that are CD49b^+^. However, significant redundancies occur within ILC1s which renders their classification difficult within biological systems in mice and humans. Unlike Group 1 ILC, Group 2 ILCs are marked by the transcription factor GATA3 and express type 2 cytokines IL-5, IL-9, IL-13 and amphiregulin in response to the alarmin cytokines IL-33, IL-25 and TSLP ^7–10^. Group 3 ILCs express the transcription factor RORγt and produce cytokines IL-17 and/or IL-22 in response to IL-1β, IL-6 and IL-23, and can be further divided on the basis of NKp46 (NCR) expression ^11–18^.

The role of NK cells in eradicating tumors has been well established, but the function of helper like ILC1s within the tumor microenvironment (TME) is yet to be fully understood. Gao *et al* ^19^ showed that in the presence of TGFβ, NK cells (CD49b^+^) can be converted to a helper ILC1 like phenotype (CD49a^+^), which in turn promote tumor progression through a TNFα dependent mechanism. The function of group 2 ILCs in pro and anti-tumorigenic responses has also been explored by a number of reports whereby ILC2 derived IL-13 drove tumor growth in two different tumor models ^20, 21^ whereas IL-5 production by ILC2s inhibited tumor progression in lung metastasis ^22^. Further reports on ILC2s have demonstrated that IL-33 drives anti-tumor activity by promoting adaptive immune responses ^23–25^. Similarly, IL-22 expressing ILC3s have been implicated in pro-tumor responses in bacteria-induced colon cancer ^26, 27^. Conversely, anti-tumor effects of ILC3s have been demonstrated in a model of malignant melanoma ^28^. This anti-tumor function in melanoma is driven by IL-12, which in turn induces Tbet expression in ILC3s, resulting in the development of Tbet^+^ “ex-ILC3s” that resemble ILC1s in phenotype. Despite inconsistencies, emerging literature suggests that the function of ILCs within the TME is dependent on the tissue microenvironment and cytokine availability. Indeed, it has been elegantly demonstrated by Nussbaum *et al* ^29^ that lymphoid tissue resident ILCs were specifically capable of anti-tumor responses while tumor tissue resident ILCs failed to inhibit tumor progression.

While cytokine mediated regulation of ILCs has been widely investigated, the role of co-receptors in regulating ILC function has been limited. The expression of several coreceptors on ILCs have been reported recently thereby providing an avenue for boosting ILC responses in cancer ^30^. We found that programmed cell death-1 receptor (PD-1) controlled group 2 ILC function during parasitic helminth responses ^31^ and emerging literature has identified a significant role for PD-1 in ILC-2 regulation in the presence of IL-33 within the tumor microenvironment (TME) ^25^; ^32^. Tumor derived ILCs are known to express PD-1 and blocking PD-1 has been demonstrated to enhance NK cell function in driving anti-tumor responses ^30 33^.

In this study, we sought to investigate whether PD-1 regulated ILC subsets in the TME, in the absence of IL-33. Using an unbiased scRNA-seq approach, we found that there was an increase in NK1.1^−^Tbet^+^ cells within the TME that was controlled by PD-1. This population although possessed type 1 signatures of ILCs, did not express NKp46 or NK1.1 protein as measured by AbSeq antibodies. We found that the NK1.1^−^Tbet^+^ population could be identified in both WT and PD1 deficient mice in steady state. In tumor, PD1 deficiency resulted in a significant increase in the frequency of NK1.1^−^Tbet^+^ cells. We found this subset upregulated PD-1 in the presence of tumour derived lactate and PD-1 upregulation significantly downregulated mTOR mediated proliferation of this novel subset within the TME. Taken together this work is the first to define a subset of NK1.1^−^Tbet^+^ ILCs within the TME that is regulated by PD-1.

## Results

### PD-1 expression is noted within Tbet^+^ILCs subpopulations in tumor

The role of PD-1 in IL-33 driven ILC-2 anti-tumor responses has been reported in models of pancreatic cancer ^25^ and in metastatic melanoma ^32^, but whether PD-1 function is restricted to IL-33 microenvironment is unknown. Here we sought to determine ILC subsets within TME that is controlled by PD-1. WT or *pdcd1*^*−/−*^ mice were engrafted with B16F10 melanoma cells (B16) and the tumor growth was monitored until day 12 (**Fig. S1a**). Tumors were harvested at day 12 post inoculation and tumor derived lymphocytes were isolated. Using an unbiased approach, we investigated the distribution of Lineage^−^Thy1^+^ subsets and their relative PD-1 expression within the TME of B16-BL6 murine melanomas. For this purpose, we combined PD-1 protein expression with transcriptomic analysis using the BD Rhapsody platform. We subjected enriched Lineage^−^ Thy1^+^ ILCs from the TME (n=5 WT and n=5KO mice) to single cell analysis. Initial clustering analysis identified 13 clusters within the enriched TILs (**Fig.1a**). First, we investigated whether the TILs clustered in a similar fashion in the WT and PD1 knock out mice. On comparing WT and PD1^−/−^ TIL populations, both WT and PD1^−/−^ TIL populations clustered in a similar fashion but within the KO, an increase in cell numbers were noted within certain clusters (**Fig.1b**). The differential gene expression patterns and pathway analysis of the 13 clusters was done in order to identify immune populations that occupy the TME (Supplementary **Table 1&2; S2, S3**). Differential gene expression analysis identified cluster 5 and cluster 1 as predominantly expressing CD3 and NK1.1 protein (Abseq). Of note cluster 10, possessed a type 1 phenotype but did not express NK1.1 (AbSeq) or NKp46 (AbSeq) protein on the surface (**Fig.1c)**. We next investigated the transcriptomic profile of cluster 10 and found that cluster 10 was enriched in type 1 genes such as *Tbet (Tbx21), Ifng, Stat4* as compared to cluster 1 and 5. Of note, cluster 10 also showed cytotoxic potential identified through *Gzmb* and *Gzmk* gene transcripts **(Fig.1d**). Finally, we performed quantitative analysis of proteins and transcript expression between clusters 1, 5 and 10 which identified a significant increase in the expression of CD25 (Abseq), *Gzmk, Tbx21, Stat4, Ifng, Gzmb*, in cluster 10 as compared to cluster 1 (NK1.1^+^ cluster; Fig.**1e**) and/or cluster 5 (CD3^+^ cluster). Cluster 10 expressed significantly low CD335 (Nkp46), *Gzma* and NK1.1 (**Fig. 1f**). No difference in *Eomes* transcript expression was noted between cluster 1 and 10 (**Fig.1g**). Of note, PD-1 protein expression was noted in cluster 10 suggesting that PD1 may control this population of ILCs within the TME (**Fig.1h)**. Finally, we investigated whether the NK1.1^−^Tbet^+^ population of ILCs existed under normal homeostatic conditions or whether NK1.1 was downregulated within TILs. Indeed, our single cell Abseq demonstrated that NK1.1 was expressed on TILs and was not downregulated. However, in order to confirm this finding, we determined the existance of this population in non-tumor mice. We found that WT mice possessed a similar immune cell subset within the bone marrow, liver and spleen. These cells did not express NK1.1, and were Tbet^+^RORγt^−^NKp46^−^ (**S1b-k**). On identifying this population in WT mice, we next investigated whether PD-1 deficiency regulated the frequency of these subsets under homeostatic conditions. Our data demonstrate that this population occurs both in WT and PD1 knock out mice under steady state (**Fig. S1b-k**).

**Figure 1:**
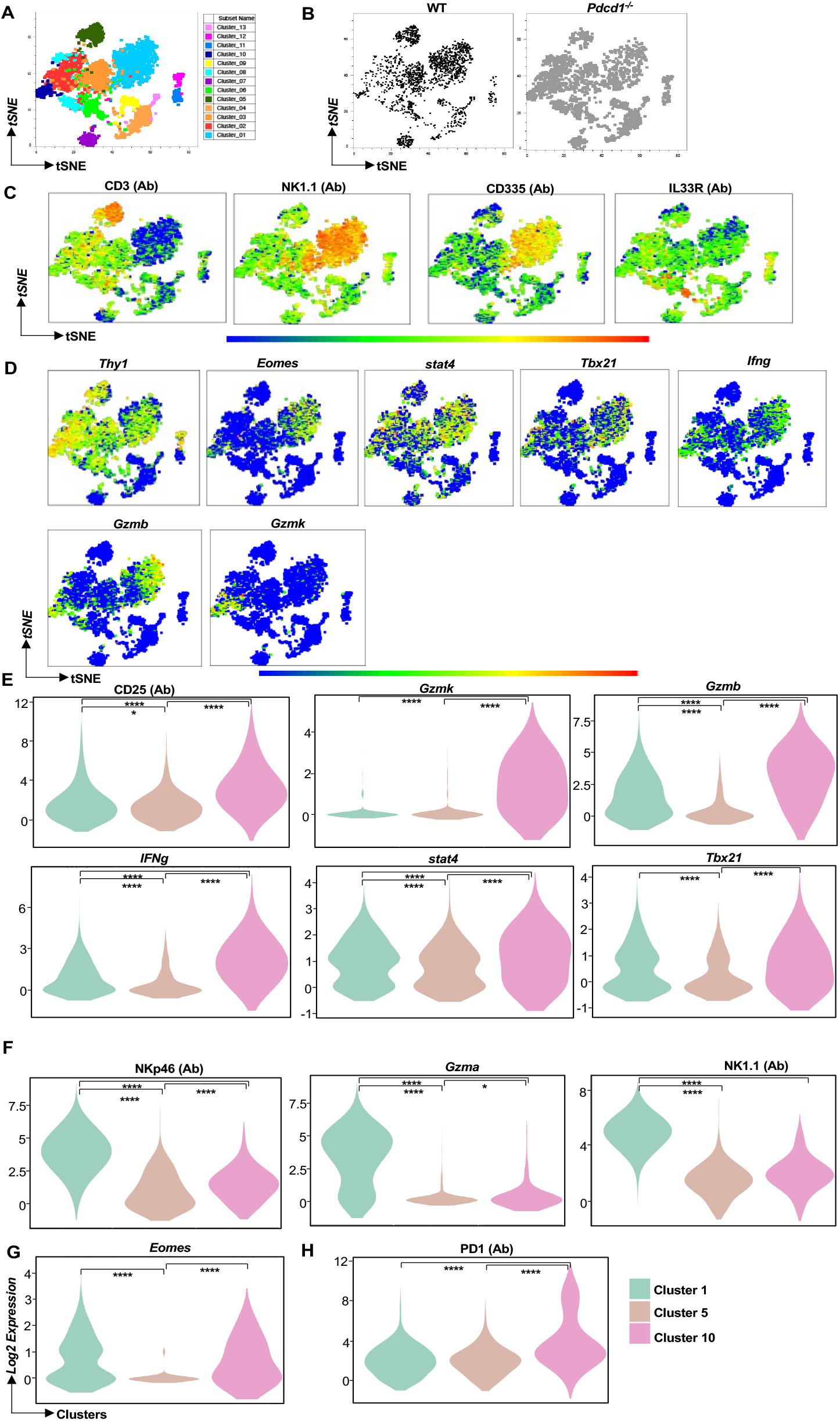
Single Cell Sequencing analysis reveals PD-1 expression in Tbet^+^ ILCs within the tumor microenvironment. C57BL6 WT and *pdcd1*^*−/−*^ mice were reconstituted with B16F10 melanoma cells via subcutaneous injection. At day 12, tumors were resected and tumor infiltrating lymphocytes were isolated and then subjected to single cell analysis. Analysis was performed with SeqGeq software with specific plugins used for gene expression analysis. Cluster analysis was performed using the phenograph plugin and the various populations are highlighted **A**, both WT and PD1 ko samples were then decoupled to show similar clustering between the samples **B**. tSNE plots of CD3, NK1.1, Nkp46 and IL33R is shown **C**. tSNE analysis of type 1 gene transcripts within the various clusters is shown in **D**. Violin plots were created using the violin plot plugin between T cell, NK cell and cluster 10 cells and significant protein and gene transcript changes is shown in **E-H**. Data shown are form n=5 WT and n=5 KO mice. Statistical analysis used was Kruskal wallis test.

### PD-1 deficiency specifically increases Tbet^+^ NK1.1^−^ ILCs within the TME but not in other target tissues

In order to further characterize the Tbet^+^ NK1.1^−^ ILCs subset, we generated PD1^−/−^*Tbet*^*ZsGreen*^ mice. WT *Tbet*^*ZsGreen*^ and PD1^−/−^*Tbet*^*ZsGreen*^ mice were reconstituted with B16 melanoma and tumour growth was monitored (**Fig. S4a-c**) which showed significance decrease in tumor growth in the PD1^−/−^*Tbet*^*ZsGreen*^ mice. In subsequent experiments, tumors were resected at day14 and then ILC subsets were determined within the TME. The expression of PD-1 in Tbet^+^NK1.1^−^ ILCs and ILC2s were first measured by flow cytometry within TILs (**S4d)**. In line with previous observation, PD-1 expression was noted in ILC2s. Of significance was the expression of PD-1 on Tbet^+^ NK1.1^−^ ILCs. Therefore, we investigated which NK1.1^−^Tbet^+^ cells (Eomes^−/−^) were controlled by PD-1 within the TME. In the absence of PD-1, a substantial increase in Tbet expression was noted in the NK1.1^−^RORγt^−^ ILC subset within the TME (**Fig. 2a**). On cumulative analysis, we found that PD-1 specifically regulated the Tbet^+^Eomes^−^ RORγt^−^ (i.e. NK1.1^−^Tbet^+^ ILC) subset within the TME (**Fig. 2b-d**) but such regulation was not apparent in the small intestine and skin but a significant difference was noted in the lungs(**Fig. S4 f-h**). Within the secondary lymphoid organs, again PD-1 deficiency was associated with no significant increase in the Tbet^+^ subset in the spleen but a significant increase was observed within the tumor draining lymph nodes (**Fig.S4i-j**). These data suggest that PD-1 specifically regulates Tbet^+^ ILCs within the melanoma TME and the associated draining lymph nodes.

**Figure 2:**
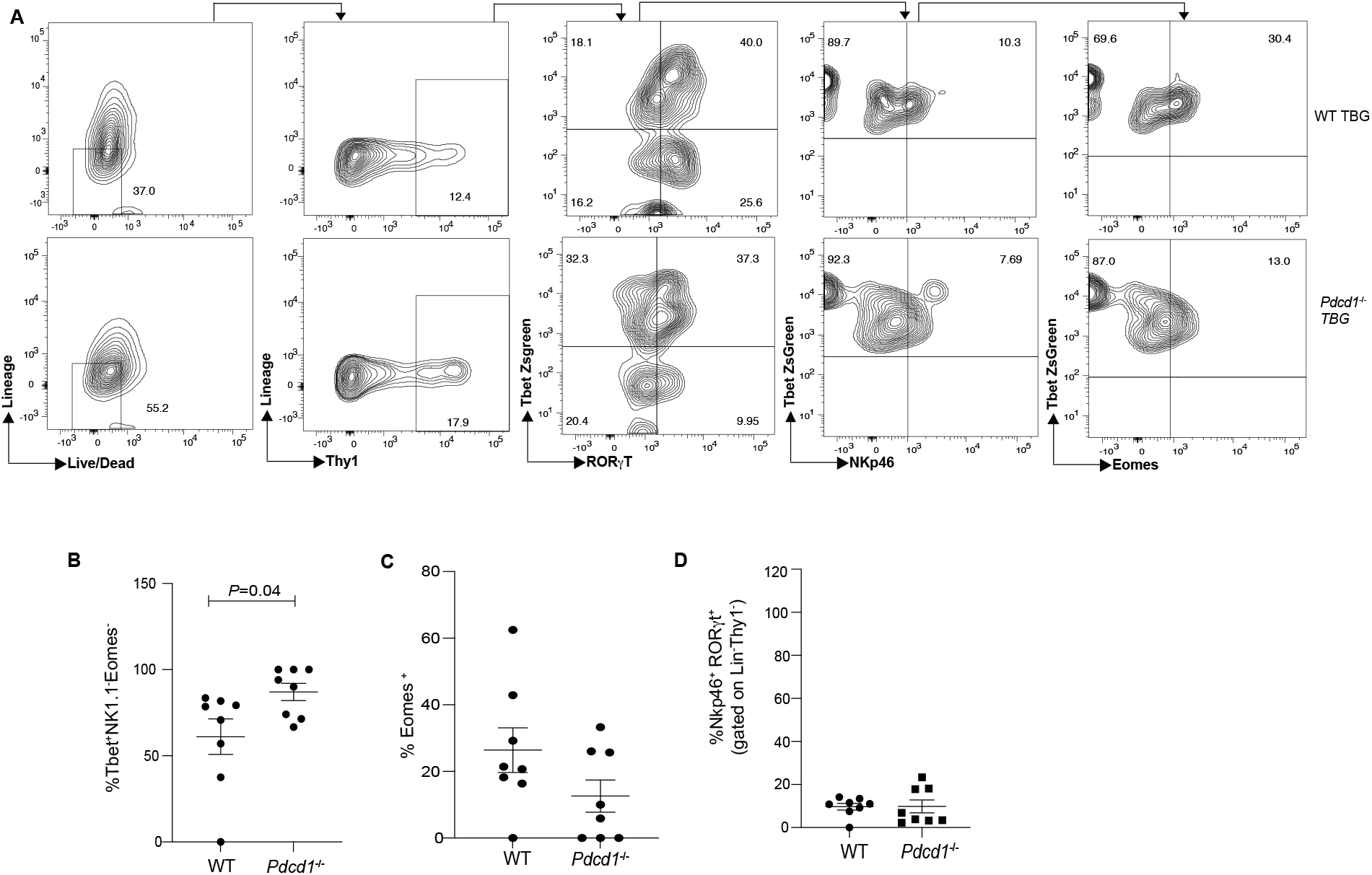
PD-1 deficiency significantly increases Tbet^+^RORγt^−^Eomes^−^NK1.1^−^ ILCs within the B16 TME. C57BL6 WT *TbetZsGreen* mice (WT TBG) or *Pdcd1*^*−/−*^*TbetZsGreen* mice (PD1^−/−^ TBG) were reconstituted with B16F10 melanoma cells via subcutaneous injection. At day 12, tumors were resected and tumor infiltrating lymphocytes were isolated and characterized by flow cytometry. ILCs were characterized as Lineage^−^Thy1^+^, lineage gate included antibodies to CD3, CD4, NK1.1, CD49b, CD5, CD8, CD11b, CD11c, CD19, Ter119, F4/80, B220, and Gr1. Representative flow cytometry plot showing helper Tbet^+^ NK1.1^−^ ILC characterization in various cohorts within the TME. Lin^−^Thy1^+^ ILCs were further characterized as RORγt^−/−^Tbet^−/−^. The ILCs within the Tbet^+^RORγt^−^ gate was further analyzed for NKp46 and Eomes expression **A**, Summary of the frequency of NK1.1^−^Tbet^+^Eomes^−^ILCs in WT and PD1^−/−^ mice within the tumor infiltrating lymphocytes is shown **B**, frequency of Eomes^+^ ILCs **C**, and NCR^+^ ILCs **D** is shown. Data shown are Mean+SEM, each data point refers to the number of mice per cohort used per experiment, statistical significance was performed using a student t test.

### PD-1 controls IFNγ production in Tbet^+^ NK1.1^−^ ILCs within the TME

Previously we found that PD-1 controlled ILC-2 cytokine secretion by regulating STAT5 ^31^. We sought to determine whether PD-1 had a similar function in tumor derived Tbet^+^ ILCs. For these experiments, since an increase in helper Tbet^+^ ILCs was noted, the cytokine potential of these cells within the TME was tested. Flow cytometry data suggested that in the absence of PD-1, a significant increase in IFNγ production was noted within the Tbet^+^ ILCs (**Fig.3a-d**) but these cells did not show increased production of IL-17 or IL-22 (**Fig.3a-d**). In addition to enhanced IFNγ, Tbet^+^ ILCs in the TME were also enriched for IFNγ+TNFα+ cells (**Fig. 3e&g**). Next, it was determined if the changes seen in ILC driven IFNγ production were also reproduced within the Lineage^+^NKp46^+^ population within the TME. Since the lineage panel included NK1.1 and CD49b, any NKp46^+^ population may have reflected the regulation of NK cells within the lineage gate. No difference was noted in IFNγ production within the NKp46^+^ Lineage^+^ gate (**Fig. 3h**). However, significant IFNγ production was noted within the Lineage^+^ gate originating from cells that are NKp46^−^ which may include T cells within the TME (**Fig. 3i**). Taken together, the data suggests that within the ILCs in the TME, PD-1 regulates cytokine production by Tbet^+^ ILC populations that possess a helper-1 like phenotype.

**Figure 3:**
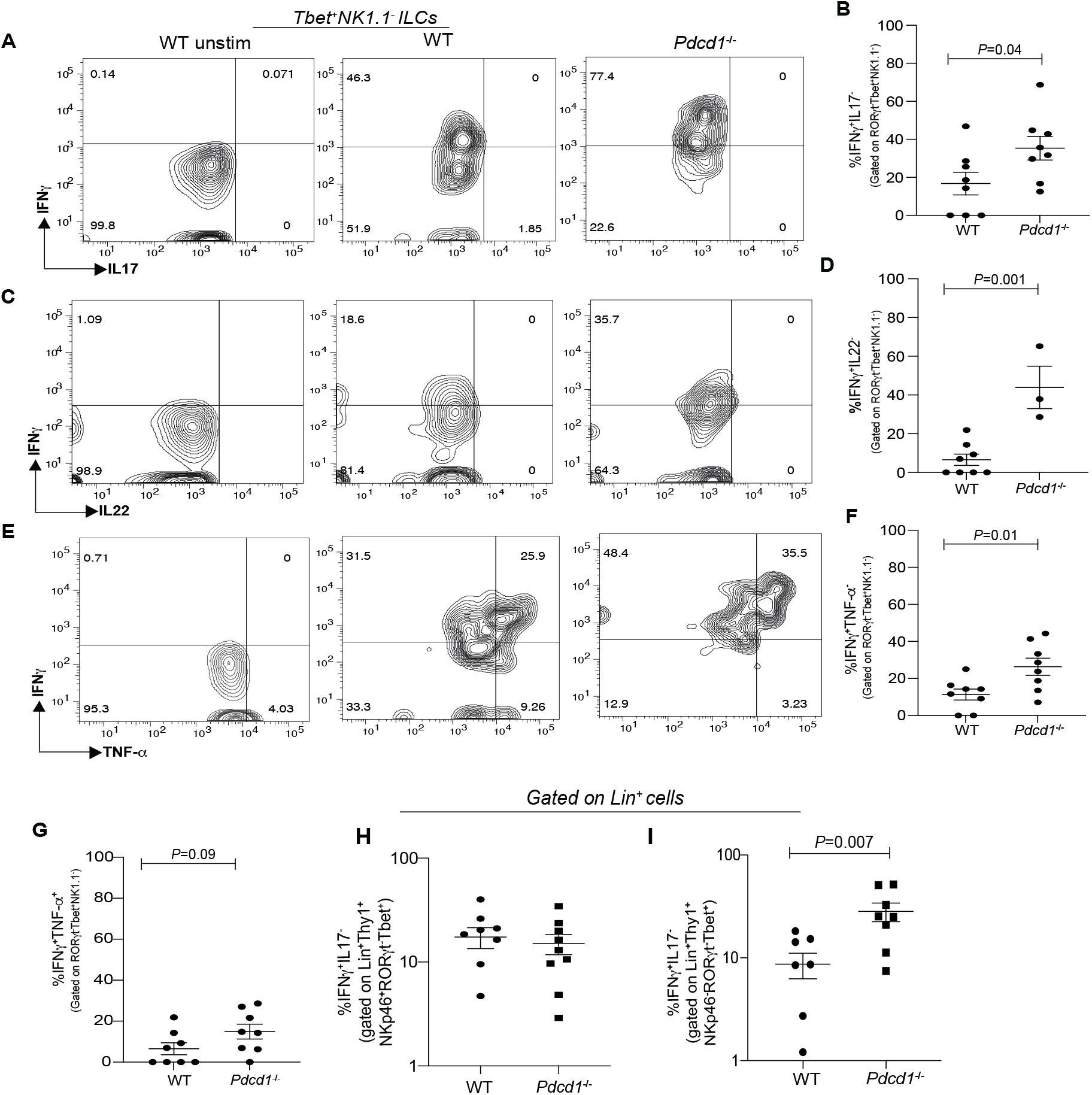
PD-1 deficiency significantly increases IFNγ+Tbet^+^ILCs within the B16 melanoma TME. C57BL6 WT *TbetZsGreen* mice (WT TBG) or *Pdcd1*^*−/−*^*TbetZsGreen* mice (PD1^−/−^ TBG) were reconstituted with B16F10 melanoma cells via subcutaneous injection. At day 12, tumors were resected and tumor infiltrating lymphocytes were isolated, stimulated for 4hrs in the presence of cytokine stimulation cocktail and then characterized by flow cytometry. ILCs were characterized as Lineage^−^Thy1^+^, lineage gate included antibodies to CD3, CD4, NK1.1, CD49b, CD5, CD8, CD11b, CD11c, CD19, Ter119, F4/80, B220, and Gr1. Representative flow plots and summary data for IFNγ and IL17 expression in WT TBG and PD1^−/−^ TBG **A-B**, Representative flow plots and summary data for IFNγ and IL22 expression in WT TBG and PD1^−/−^ TBG **C-D**, Representative flow plots and summary data for IFNγ and TNFα expression in WT TBG and PD1^−/−^ TBG **E-G**. The expression of IFNγ in Lineage^+^NKp46^+^ TILs **H** and Lineage^+^NKp46^−^ TILs are shown **I**. Data shown are Mean+SEM, each data point refers to the number of mice per cohort used per experiment, statistical significance was performed using a student t test.

### PD-1 controls proliferation of Tbet^+^ NK1.1^−^ ILCs within the TME

The molecular mechanism by which PD-1 regulated the frequency of Tbet^*+*^ *NK1*.*1*^*−*^ ILCs within the TME was next determined. First, PD-1 expression pattern in Tbet^+^ ILCs within the TME and normal surrounding skin was evaluated. Tbet^+^ILCs within the TME significantly upregulated PD-1 expression when compared to their counterparts within normal skin (**Fig.4a; S5a**). This suggested that PD-1 was selectively enhanced within the TME *Tbet*^*+*^ *NK1*.*1*^*−*^ ILCs. Next, the ability of tumor cells to induce PD1 on Tbet^+^ ILCs was evaluated. In transwell experiments, B16 melanoma tumor cells co-cultured in transwell plates with ILCs induced PD-1 with an increase in frequency in PD1^+^ ILCs observed at the 1:1 ratio at 6hrs, with a trend noted at 1:10 ratio which was completely abrogated at a 1:100 ratio (**Fig. 4b, S5b)**. Since tumor derived lactate can suppress proliferation of ILC2s ^24^, it was next investigated whether lactic acid was produced by B16 tumors and if lactic acid induced PD-1 on Tbet^+^ILCs. In *in-vitro* experiments, tumor supernatants contained significant amounts of lactic acid (**Fig. S5c**) and lactic acid addition significantly increased PD-1 expression on Tbet^+^ILCs (**Fig. 4c, S5d**). We next investigated if tumor cell supernatant affected the proliferative capacity of ILCs through PD-1. We measured the ability of WT NK1.1^−^ Tbet^+^ ILC proliferation in the presence of tumor cell supernatant and then compared the relative proliferation of PD1^−/−^Tbet^+^ILCs. We found that B16 tumor supernatant significantly inhibited proliferation of WT NK1.1^−^ Tbet^+^ ILC as compared to PD1 deficient cells (**Fig. 4d-e**). Furthermore, within the B16 TME *in-vivo*, a significant increase in the proliferative potential of Tbet^+^ ILCs (as measured by Ki67 staining) was noted in PD1^−/−^ cohorts (**Fig. 4f-g**). Subsequently, we tested the molecular mechanism by which PD-1 enhanced proliferation within ILC subsets. Recent studies have explored the role of PD-1 in altering the metabolic phenotype of T cells ^34^ and myeloid cells ^35^. Specifically, metabolic regulation of myeloid cells within the TME by PD-1 has been shown to enhance anti-tumor responses. PD-1 has also been implicated in regulating ILC2 metabolism in allergic inflammation ^36^. In line with previous observations ^36^, PD-1 significantly regulated fatty acid metabolism and glycolysis in WT mice (**Fig. 4h-k**). We next investigated if a change in metabolic signatures could be identified at a single cell resolution within the WT and PD-1 KO ILC populations within the TME. We found that the PD-1 KO type I ILCs from cluster 10 had significantly decreased *apoe* gene transcript as compared to the WT cohorts (**Fig. S5f**). In keeping with increased glycolysis, PD-1 deficiency also increased mTOR signaling in Tbet^+^ILC TILs within B16 TME as identified by phosphorylated p70S6K (**Fig. 4l, S5g**). The molecular mechanism was confirmed in WT Tbet^+^ILC TILs whereby PD-1 blockade enhanced mTOR signaling and p70S6K phosphorylation (**Fig. 4m, S5h)**. Taken together, we propose the following molecular mechanism by which PD-1 regulates ILCs within the TME. Depending on the active alarmins within the TME, PD-1 expression is increased in the corresponding ILC subset (IL-33 in the case of ILC2s, and melanoma tumor cell derived lactate in case of Tbet^+^ ILCs). Blocking PD-1 enhances the proliferative capacity of the ILC subsets within the TME by enhancing glycolysis and upregulating the mTOR signaling pathway.

**Figure 4:**
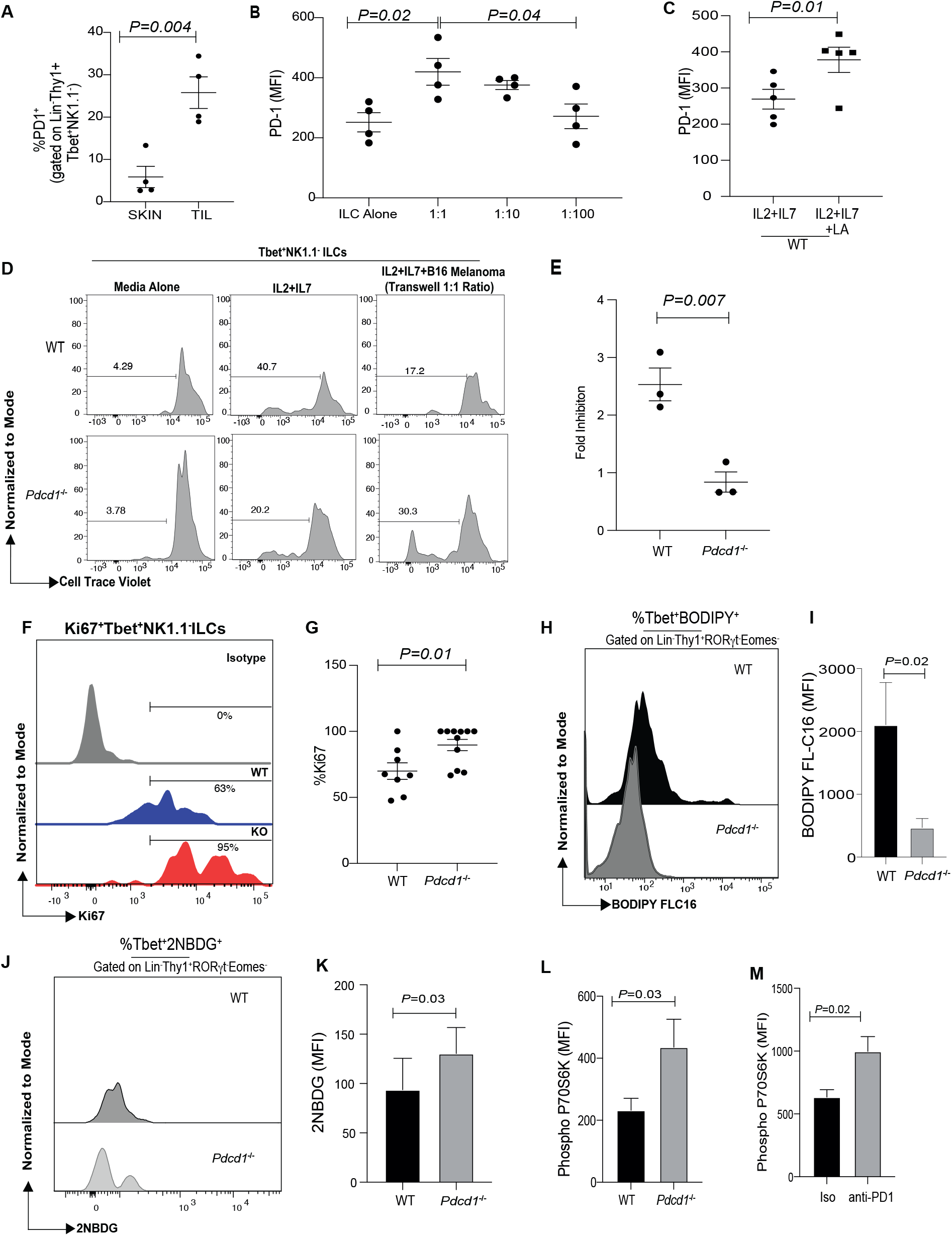
PD-1 inhibits the proliferation of Tbet^+^NK1.1^−^ ILCs within the B16 TME. C57BL6 WT *TbetZsGreen* mice (WT TBG) were reconstituted with B16F10 melanoma cells via subcutaneous injection. At day 12, tumors were resected and tumor infiltrating lymphocytes were isolated. In addition, ILCs were also isolated from surrounding normal skin. ILCs were characterized as Lineage^−^Thy1^+^, lineage gate included antibodies to CD3, CD4, NK1.1, CD49b, CD5, CD8, CD11b, CD11c, CD19, Ter119, F4/80, B220, and Gr1. PD-1 expression was measured in skin derived and tumor derived Tbet^+^RORγt^−^ILCs **A**. Transwell experiments were repeated with B16 melanoma cells at various ratios and then PD-1 expression was monitored in cultures stimulated with either cytokines alone (IL2, IL7) or in the presence of B16 melanoma cell line at 1:1, 1:10 (1 melanoma cell:10 lymphocytes) and 1:100 ratio after 6hrs. PD-1 protein expression on Tbet^+^NK1.1^−^ ILCs at various ratios was measured by flow cytometry **B**. Splenocytes were incubated for 24hrs with IL2 and IL7 alone or with lactic acid and then PD-1 expression on Tbet^+^ILCs were measured **C**. Transwell experiments were repeated with a 1:1 ratio of B16 melanoma cells along with IL2 and IL7. Splenocytes were stained with cell trace violet and then proliferation was measured at day 5 post stimulation. Representative flow plots showing proliferation of Tbet^+^RORγt^−^NK1.1^−^ ILCs from WT and PD1^−/−^ cultures are shown **D**, Fold inhibition of proliferation in the presence of melanoma supernatant in WT and PD1^−/−^ NK1.1^−^Tbet^+^ ILCs is shown **E**. Fold Inhibition was measured as follows. WT ILC proliferation in response to IL2 and IL7 was measured and then the rate of proliferation with B16 melanoma supernatant was measured. Fold inhibition between the two culture conditions was calculated and then plotted for WT and KO. C57BL6 WT *TbetZsGreen* mice (WT TBG) or *Pdcd1*^*−/−*^*TbetZsGreen* mice (PD1^−/−^ TBG) were reconstituted with B16F10 melanoma cells via subcutaneous injection. At day 14, tumors were resected and tumor infiltrating lymphocytes were isolated, stimulated for 4hrs in the presence of cytokine stimulation cocktail and then characterized by flow cytometry. ILCs were characterized as Lineage^−^Thy1^+^, lineage gate included antibodies to CD3, CD4, NK1.1, CD49b, CD5, CD8, CD11b, CD11c, CD19, Ter119, F4/80, B220, and Gr1. Representative flow cytometry showing Ki67 expression in Tbet^+^RORγt^−^NK1.1^−^ tumor derived ILCs from WT and PD1^−/−^ cohorts **F** and cumulative data shown in **G**. BODIPY (4,4-Difluoro-5,7-Dimethyl-4-Bora-3a,4a-Diaza-s-Indacene-3-hexadecanoic Acid) MFI is shown in **H** and (**I**, n=5, paired one-tailed student t test); 2NBDG uptake is shown in **J**,) and (**K**, n=4, paired one-tailed student t test). TILs were harvested and stimulated with IL2 and IL7 for 15mins and then phosphoP70S6Kinase was measured (**L**, n=4, paired one-tailed student t test). WT TILs were stimulated for 3 days with either isotype control or anti-PD1 antibody. At day 3, TILs were stimulated for 15 mins with IL2 and IL7 and then phosphoP70S6Kinase was measured (**M**, n=3, paired one-tailed student t test). Data shown are Mean+SEM, n=4-5 mice per cohort used per experiment, each data point represents the number of mice used, statistical significance was performed using a student t test.

### PD1 regulates Tbet^+^NK1.1^−^ILCs within the TME in AOM-DSS induced colorectal cancer

In order to test the reproducibility of our observation within the orthotopic melanoma model, we next investigated if PD-1 controlled ILCs within the TME in an inducible model of cancer within a different tissue. A model of colorectal cancer (CRC) was chosen since it has been shown that ILCs support CRC growth via IL-22 and we wondered if within this model, PD-1 can increase Tbet^+^NK1.1^−^ ILCs with a type 1 phenotype. In AOM-DSS, PD-1 mice were significantly more susceptible to dextran sodium sulphate (DSS) mediated weight loss during the first cycle but susceptibility was reduced in the third cycle as compared to WT cohorts (**Fig. 5a**). In addition, there was significantly lower number of tumors noted within the intestine of *pdcd1*^*−/−*^ mice (**Fig.5b**) along with an increase in inflammation within the intestine (as measured by the intestinal length, with reduced length associated with greater inflammation; **Fig. 5c**). We next harvested the tumors to determine the frequency of ILCs. Similar to our melanoma data, we found a significant increase in Tbet^+^NK1.1^−^ ILCs within the tumors (**Fig. 5d-g**). Of note, no difference was noted within the normal tissue or in MLNs (data not shown). These data suggest that PD-1 deficiency can enhance Tbet^+^NK1.1^−^ ILCs within CRC.

**Figure 5:**
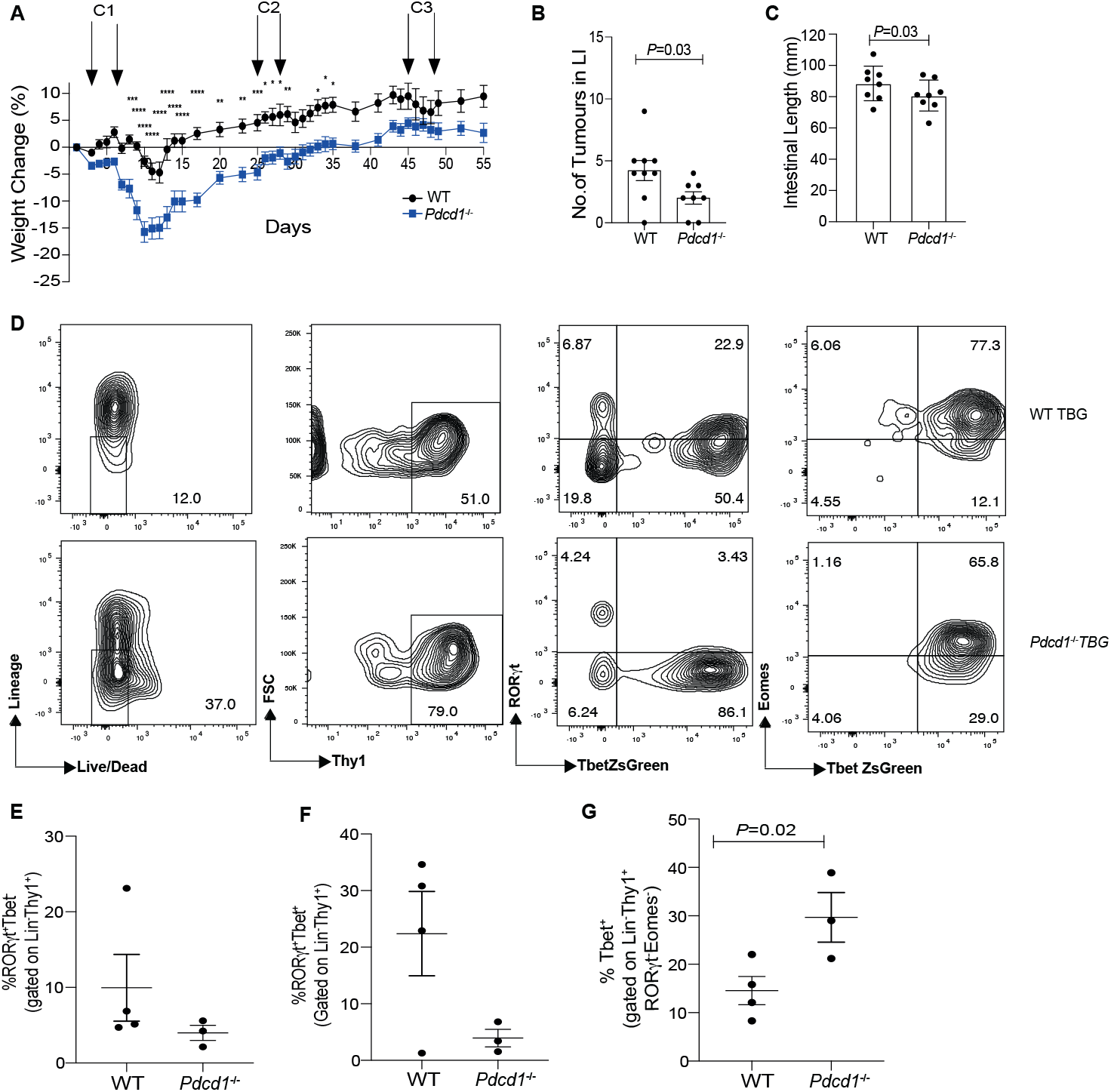
PD-1 regulates Tbet^+^RORγt^−^ ILCs within the colorectal TME. C57BL6 WT *TbetZsGreen* mice (WT TBG) or *Pdcd1*^*−/−*^*TbetZsGreen* mice (PD1^−/−^ TBG) were treated with one dose of AOM followed by three cycles of DSS. Weight loss was monitored in the cohorts during the experimental protocol as a measure of clinical signs of inflammation **A**. Animals were harvest at day 55 and then no of tumors were counted within the intestine **B** and the intestinal length was measured **C**. Tumors were resected and ILC frequency was measured by flow cytometry. ILCs were characterized as Lineage^−^Thy1^+^, lineage gate included antibodies to CD3, CD4, NK1.1, CD49b, CD5, CD8, CD11b, CD11c, CD19, Ter119, F4/80, B220, and Gr1. Representative flow cytometry is shown in **D** and cumulative data is shown in **E-G**. Data shown are Mean+SEM, statistical significance was performed using a student t test and number of mice in each cohort is represented as a data point in the graph.

### Blocking antibodies to PD-1 significantly increases NK1.1^−^Tbet^+^ILCs in subcutaneous and metastatic B16 melanoma

We next tested if the phenomenon in the PD-1 KO mice can be reproduced in a therapeutic context. Since an increase in NK1.1^−^Tbet^+^ ILCs were noted both in the TME and lungs (**see Fig. S4f**) we tested whether these ILC subsets were increased in metastatic melanoma model. WT mice were subcutaneously reconstituted with B16 cells and then tumors were allowed to engraft. Mice were treated with either isotype or anti-PD1 at days 7, 9, 11, 12 and the tumor growth curve was monitored (**Fig. 6a**). Mice were euthanized on day 14 and then ILC subset frequency was analyzed. A significant increase in NK1.1^−^Tbet^+^ ILCs was noted in the anti-PD1 treated cohorts as compared to the control isotype treatment cohorts (**Fig. 6b-c**). We next tested the frequency of these cells in a metastatic model of melanoma and found a similar increase in NK1.1^−^Tbet^+^ ILCs (**Fig. 6d-e**). These data suggest that anti-PD1 can significantly increase NK1.1^−^Tbet^+^ ILCs both within the subcutaneous and metastatic tumor microenvironment.

**Figure 6:**
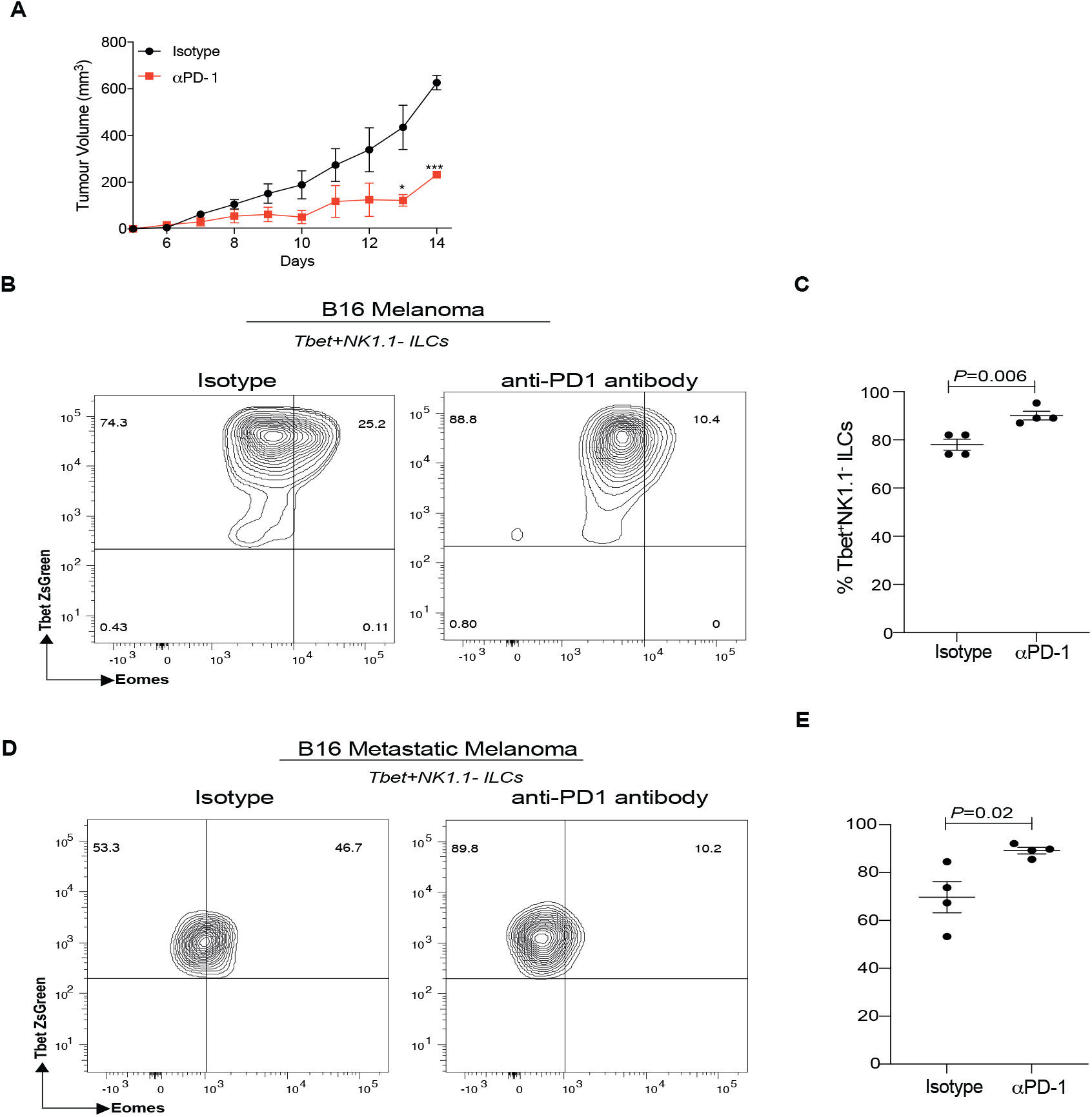
PD-1 blocking antibodies significantly increases tumor derived Tbet^+^RORγt^−^ ILCs ILCs in subcutaneous melanoma, colon carcinoma and metastatic melanoma murine models. C57BL6 TbetZsGreen mice were reconstituted with B16F10 melanoma cells via subcutaneous injection. Mice were treated with either isotype control or anti-PD1 therapy on days 7, 9, 11 and 13. Tumor growth was monitored **A**. At day 14, tumors were resected and tumor infiltrating lymphocytes were isolated. Tbet^+^ILCs were characterized as Lineage^−^Thy1^+^ RORgt^−^NKp46^−^ cells, lineage gate included antibodies to CD3, CD4, NK1.1, CD49b, CD5, CD8, CD11b, CD11c, CD19, Ter119, F4/80, B220, and Gr1. Representative flow plot depicting Tbet^+^ ILCs **B** and cumulative data from B16 melanoma TILs **C**. Data from metastatic melanoma TILs is shown in **D-E**. Data shown are Mean+SEM, each data point refers to the number of mice per cohort used per experiment, statistical significance was performed using a student t test.

### PD-1 blockade partially protects murine recipients from tumor burden by enhancing helper ILCs in the absence of NK cells and adaptive immune cells

Recent reports have shown that PD-1 blockade can enhance anti-tumor immunity by specifically targeting the innate immune cells such as macrophages in the TME ^35^. Therefore, we investigated the contribution of ILCs versus macrophages and NK cells in eliciting anti-tumor responses in the presence of PD-1 inhibition. NK1.1^+^ cell depleted *Rag*^*−/−*^ host mice were injected with B16 melanoma cells. On tumor establishment, animals were repeatedly injected with NK1.1^+^ cell depleting antibodies with or without anti-PD-1 (**Fig. 7a**). Within this experimental set up, we found that blocking PD-1 partly enhanced the survival of mice independent of NK cells (**Fig. 7b**). In addition to increased survival, a significant decrease in tumor burden was also noted in the murine recipients that received PD-1 immunotherapy on depleting NK1.1^+^ cells (**Fig. 7c, S6a**). However, using a similar experimental set up, depleting NK cells and ILCs (using Thy1 antibody) abolished any protective tumor effects rendered by PD-1 blocking in these murine recipients (**Fig. 7d-e, S6b**). It is worth noting that depletion of NK cells and ILCs continued after the establishment of tumors as these cells may play a significant role at this phase of tumor progression. We next determined the immune cell subsets that are enhanced within the melanoma TME in the absence of NK cells and adaptive immune cells. We found depleting NK cells significantly enhanced IL-7R^+^ (CD127^+^) ILCs within the TME (**Fig. 7f & S6c**). On PD-1 blockade, the frequency of IL-7R^+^ ILCs within the TME was further enhanced (**Fig. 7f**). Within this experimental set up, no difference in the frequency of myeloid and DC compartment was observed (**Fig. S6c-g**). We next investigated the cytokine profile of ILCs within the melanoma TME and found that blocking PD-1 significantly increased the frequency of ILCs that were capable of producing dual effector cytokines such as IL-17 and IFNγ (**Fig. 7g & S6h**). In line with our previous observation ^31^, we found that the IL13^+^ ILCs were enriched in IFNγ production within the TME (**Fig. S6h-i**). Taken together, our data suggests that PD-1 regulates IFNγ production and type 1 ILC phenotype within the melanoma TME.

**Figure 7:**
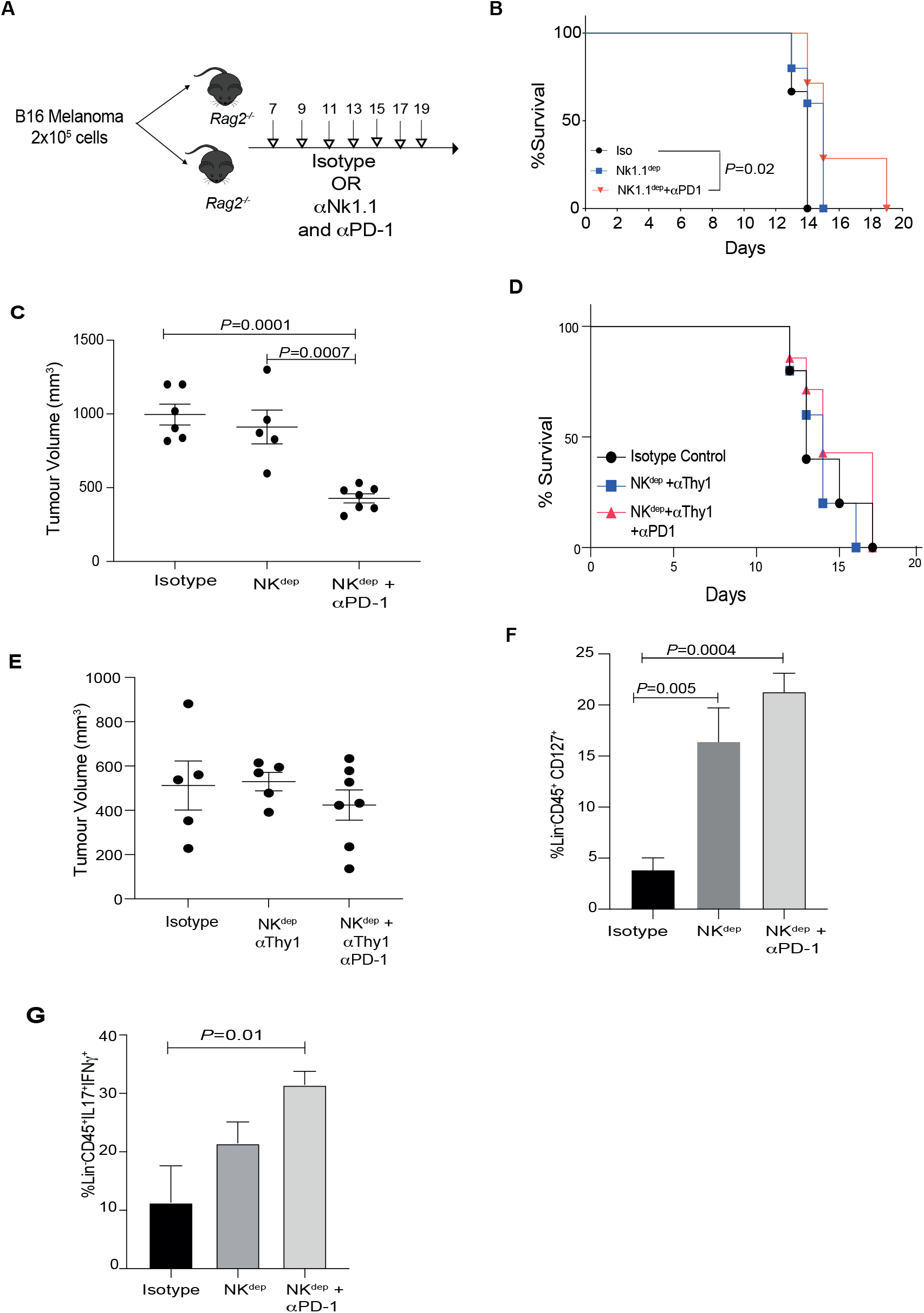
PD-1 blockade enhances the survival of tumor bearing *Rag* mice. B6 *Rag*^−/−^ mice were reconstituted with B16F10 cells via subcutaneous injection. Mice were treated with either isotype control, NK depleting antibody (NK1.1) alone or in combination with anti-PD1 antibody **A**. Survival of mice in the different cohorts were monitored over a period of 20 days **B**. The experiment was repeated in mice with either isotype control or NK depletion plus anti-PD1 therapy followed by measurement of tumor volume shown at day 14 **C**. B6 *Rag*^−/−^ mice were reconstituted with B16F10 cells via subcutaneous injection. Mice were then treated with either isotype control, NK depleting antibody (NK1.1) and ILC depleting antibody (Thy1) alone or in combination with anti-PD1 antibody. Survival of mice in the different cohorts were monitored **D** and tumor volume shown at day 12 **E**. B6 *Rag*^−/−^ mice were reconstituted with B16F10 cells via subcutaneous injection. Mice were treated with either isotype control, NK depleting antibody (NK1.1) alone or in combination with anti-PD1 antibody. At day 12, tumors were resected and tumor infiltrating lymphocytes were isolated and characterized by flow cytometry. Helper ILCs were characterized as Lineage^−^CD45^+^CD127^+^, lineage gate included antibodies to CD3, CD5, CD8, CD11b, CD11c, CD19, Ter119, F4/80, B220, and Gr1^+^. Frequency of helper ILCs in shown in **F**. TILs were stimulated with cytokine stimulation cocktail for 4hrs. Cells were then subjected to intracellular flow cytometry in order to measure effector cytokines. Frequency of IFNγ and IL-17 positive ILC TILs **G**. Data are represented as Mean+SEM, with each cohort consisting of n=5-10 mice per cohort. A Kaplan-meier survival curve analysis was performed for tumor survival experiments, one-way ANOVA was performed for identifying significance in tumor volumes and ILC frequency between cohorts.

### PD-1 regulates human Tbet^+^ILC subsets in cutaneous melanoma and SCC

We next tested if PD-1 played a similar role in regulating Tbet^+^ RORγt^−^ ILC population in humans. In humans, we classified the various ILC subsets as previously described ^37 38 39^. ILC1s were classified as Lin^−^CD127^+^CD117^−^CRTh2^−^, ILC2 as Lin^−^CD127^+^CRTH2^+^ and ILC3 as Lin^−^ CD127^+^CD117^+^CRTH2^−^ RORgt^+^Nkp44^+^cells (**Fig.S7a**). ILC-1 populations were further discriminated based on Tbet expression (**Fig.S7b**) ^40^. In all three ILC populations, PD1 expression was noted with a significant expression seen in Tbet^+^ subset of ILC1 (**Fig. 8a, S7c**). We next investigated if tumor supernatant can induce PD-1 within the Tbet^+^ ILC1s. For this experiment, we utilized human melanoma and cutaneous squamous cell carcinoma cell lines. Similar to murine experiments, PD-1 expression was significantly increased in the presence of melanoma and cSCC tumor supernatants (**Fig. 8b, S7d**). Next, we investigated if this increase had a functional impact on ILC proliferation similar to the murine observations. We found that similar to the murine studies, an increase in Tbet^+^ ILC proliferation was noted on blocking PD-1 in normal human donors (**Fig. 8c-d, S7e**). We next tested whether our observation on human ILCs with cSCC cell lines were relevant in primary tumors of human cSCC. Recently, a comprehensive atlas of human cSCC has been published, along with the report that ILCs can populate cSCC tumours ^41 42^. However, whether Tbet^+^ILCs exists and elicit functional anti-tumour responses within cSCC is unknown. Moreover, whether PD-1 restricts ILC anti-tumor responses in human cSCC is unclear. Our data confirms that helper ILCs are present within human cSCC tumour microenvironment. (**Fig.8e**). Out of these various populations, high PD-1 expression was associated with cluster 12 (ILC2) and Tbet (clusters 6 & 13) (**Fig. 8f**). In addition to Tbet^+^ ILCs, Nkp44^+^ epithelial ILCs were also noted within the TILs which expressed PD-1 (cluster 1, Fig. 8f). However, given that CD56 was included in the lineage gate, caution needs to be applied in interpreting the role of PD-1 in NKp44^+^ epithelial ILCs which will be part of future work.In line with Luci *et al*, we also found that ILC2 cluster was significantly reduced within cSCC microenvironment. Next, in order to demonstrate functional significance of PD-1 on ILCs, we tested whether PD-1 blockade can enhance Tbet^+^ ILC subset in primary cSCCs. Our data demonstrates that culturing ILC TILs in the presence of PD-1 blocking antibody can significantly increase Tbet^+^ ILCs (**Fig. 8g-h**). Our data is the first to identify a functional role for PD-1 on Tbet^+^ILCs in cSCC. Taken together, these data suggest that ILCs are likely to be involved in tumor immunity and that effects on the function of PD-1 expressing ILCs during anti-PD-1 immunotherapy regimens may play a role in enhancing patient anti-tumor responses.

**Figure 8:**
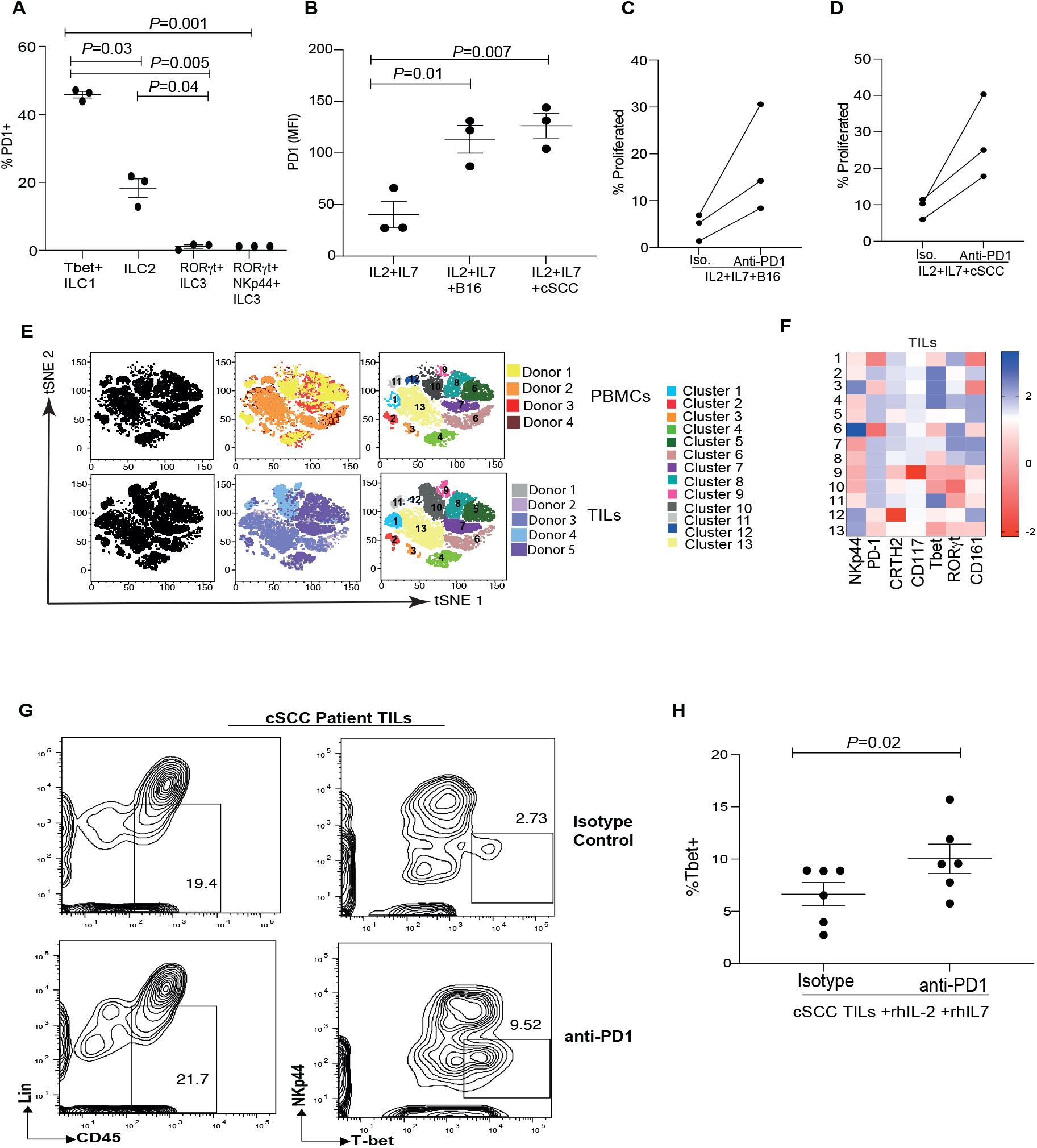
PD-1 regulates human Tbet^+^ ILC proliferation in human cSCC. Expression of PD-1 in various ILCs subsets from normal human donor peripheral blood mononuclear cells is shown **A**. Human melanoma and cutaneous squamous carcinoma cell lines were incubated at a 1:1 ratio in a 24 well transwell plate in the presence of normal donor PBMC. PD-1 protein expression within Tbet^+^ ILCs from the various culture conditions is shown in **B**. The rate of proliferation of Tbet^+^ RORγt^−^ ILCs in the presence of melanoma either with isotype or anti-PD1 antibody is shown in three donors **C**, similarly the proliferation of Tbet^+^ RORγt^−^ ILCs in the presence of cSCC either with isotype or anti-PD1 antibody is shown in three donors **D**. Blood and tumor tissue were obtained from cSCC patients and then helper ILC subsets were characterized using flow cytometry. Tbet^+^ ILCs were characterized as Lin^−^CD45^+^CD127^+^CRTH2^−^CD117^−^ cells. Lineage gate included CD3, CD5, CD11b, CD11c, CD14, CD16, CD19, CD20, CD56 and TCRα/β. tSNE analysis of ILC population in 5 cSCC donors and 4 matched blood donors are shown in **E-F**. TILs were expanded for 3 days in the presence of hIL-2, hIL-7 and either in the presence isotype antibody or anti-PD1 antibody. Frequency of Tbet^+^ ILCs were measured by flow cytometry **G-H**. Data shown are Mean +SEM from n=3-5 cSCC and blood samples. Statistical significance was performed using a student t test.

## Discussion

The efficacy of anti-PD-1 based therapies for multiple cancers, such as melanoma, bladder, lung cancers and Hodgkin lymphoma, has been overwhelmingly demonstrated by clinical trials and experimental murine models. The central premise of anti-PD-1 based therapies suggest that these interventions rescue dampened CD8^+^ T cell function within the TME resulting in more robust anti-tumor responses. Emerging literature demonstrates that PD-1 can also enhance innate immunity within TME but the initial innate immune population that is activated by PD-1 in exerting anti-tumor effect is unclear with some studies suggesting NK cells while others indicate myeloid cells 33, 43. The ability of PD-1 to regulate innate immune cell function, including ILCs implicate a potential use for checkpoint inhibitors in harnessing tissue mediated immunity within the TME that are not enriched in inflammatory infiltrates. Indeed, supporting this hypothesis are recent reports demonstrating that blocking antibodies to PD-1 can drive IL-33 primed ILC-2 function within the pancreatic and metastatic TME ^25 32^.

The role of ILCs in promoting antitumor responses has been controversial since all helper like ILCs have been implicated in both pro and anti-tumorigenic functions within the literature. However, whether these populations can be harnessed by using checkpoint inhibitors is not known. We have found novel roles for PD-1 on ILCs within TME, which include (1) identifying a subset of NK1.1^−^Tbet^+^ ILC populations within TME that expressed PD-1, (2) determining that PD-1 selectively increases the frequency and function of NK1.1^−^Tbet^+^ ILCs within a number of solid cancers, and (3) demonstrating PD-1 regulates the proliferation of both murine and human NK1.1^−^ Tbet^+^ ILCs by harnessing metabolic regulation. To our knowledge this is the first study to demonstrate that Tbet^+^ NK1.1^−^ ILCs occupy the TME and are regulated by PD-1. Our study demonstrates a novel type 1 subset and identifies a previously unknown regulatory role for PD-1 within tumor resident Tbet^+^ NK1.1^−^ ILCs.

In B16, the role of ILC2s in driving anti-tumor responses has been extensively reported ^22 44 44 45 24^, within the context of IL-33 with no data on whether PD-1 can regulate ILCs in the absence of cytokines. In our experimental systems, using a single cell approach, experimental murine models and ex vivo human immune cell cultures, we found that PD-1 knock out mice had significantly increased number of NK1.1^−^Tbet^+^ ILCs in various TMEs. In addition, this population was found in both WT and PD1 KO mice in similar frequency within steady state. These data suggest that although healthy tissues are enriched in different ILC subsets, when perturbed with tumor, anti-PD-1 treatment can expand NK1.1^−^Tbet^+^ ILC in different tumor types and enhance production of IFNγ by this subset. These data drive the possibility of uniform treatment strategies for multiple solid cancers that are not enriched for inflammatory infiltrates whereby PD-1 blockade could be used to boost tissue resident NK1.1^−^Tbet^+^ ILC which possess a type 1 phenotype.

Our data also suggest that there is a functional consequence to increased NK1.1^−^Tbet^+^ ILC numbers within the TME. In line with previous reports, depleting NK cells in *Rag* mice did not inhibit tumor progression ^24^ but combining NK depletion with PD-1 blocking antibody significantly increased survival of *Rag* mice with B16 tumors and showed significant reduction in tumor volume. This anti-tumor response was primarily driven by ILCs with a type 1 phenotype. We did not find any differences within other innate immune cell compartment including macrophages as previously reported ^43^, and on depleting ILCs in *Rag* mice, any protection against tumor was lost. The difference in our study versus Gordon *et al*, could be due to the timing of treatment of the tumor. In our experimental set up, we first allowed the tumors to be established and then deleted ILCs using antibody treatment and therefore were able to isolate the treatment effect of PD-1. We conclude that in the absence of adaptive immunity, myeloid and NK cells, NK1.1^−^Tbet^+^ ILCs play a predominant role in anti-tumor responses in the presence of anti-PD1 blocking antibodies. We also propose that NK1.1^−^Tbet^+^ ILCs may drive robust adaptive immune responses in the presence of anti-PD1 blocking antibodies.

Our data contradicts a study on the role of PD-1 in modifying NK cell function in a metastatic lung melanoma model whereby a significant decrease in tumor size was noted with PD-1 blockade, which the authors largely attributed to NK cells ^33^. This can be due to the differences in the murine model utilized since our experiments were focused on the orthotopic model of melanoma and was not tested in the metastatic model. Also, in our model, NK depletion studies were performed in *Rag* recipients rather than WT mice as performed by Hsu et al ^33^, suggesting that there may be off-target effects from the adaptive immune system that may modulate NK cell function within the WT mice metastatic environment. Of note, recent reports have suggested that the expression of PD-1 on NK cells may be co-opted from other immune cells within the TME further suggesting a bystander role for adaptive immune cells in contributing to PD-1 effect on NK cells ^46, 47^.

In contrast to our previous observation in host pathogen responses ^31^, we found that the frequency of ILC2s were negligible within the tumor in PD1 ko mice (data not shown). Indeed, this phenomenon is in line with other studies on melanoma TME, where significantly reduced ILC2 numbers within the tumor were noted due to increased lactate production ^24^. This immune suppression mediated by melanoma can be overcome by administrating IL-33 ^24^ and furthermore PD-1 blockade in combination with IL-33 can enhance ILC2 mediated chemokine production. Increase in chemokine production recruited DCs which in turn enhanced CD8^+^ tumor immunity in pancreatic cancer ^25^. Herein lies the importance of our current study which specifically focusses on how PD-1 regulates anti-NK1.1^−^Tbet^+^ ILCs in the absence of IL-33, within the TME. Our data uncouples the effect of IL-33 and PD-1 within the TME and demonstrate that PD-1 selectively regulates Tbet^+^ILCs.

The molecular mechanism by which PD-1 can regulate Tbet^+^ILC populations within the TME is not known. We found a unique expression pattern for PD-1 on ILCs whereby it was restricted to the TME ILCs and not surrounding normal skin. Furthermore, within the TME, PD-1 expression is selectively increased on Tbet^+^ ILCs, suggesting a role for tumor derived products activating the immune compartment in TME resulting in PD-1 expression. In line with Wagner et al, we found that lactate which is produced by melanoma cells was capable of inducing PD-1 on Tbet^+^ ILCs. Blocking PD-1 specifically enhanced proliferation of Tbet^+^ ILC in both *in vivo* murine and *in vitro* human tumor models. These data demonstrate a new tumor derived immune evasion program driven by lactate and PD-1 by which Tbet^+^ ILC mediated anti-tumor responses are inhibited.

We next sought to investigate if Tbet^+^ ILC proliferation was controlled by the metabolic regulation of ILCs by PD-1. Indeed, in line with ILC2 biology ^48^, we found that within the TME, NK1.1^−^ Tbet^+^ ILCs predominantly utilized fatty acid oxidation for their metabolic demands. However, when PD-1 was absent, fatty acid metabolism was significantly downregulated with a concomitant increase in glycolysis similar to ILC2s ^36^. Taken together, our data suggests that PD-1 is induced on Tbet^+^ ILCs by tumor derived lactate, which then selectively inhibits the proliferation of the Tbet^+^ ILC within the TME by modulating metabolic pathways.

The mechanistic data presented within this study was performed in PD-1 deficient mice and can be reproduced in therapeutic situations in the presence of PD-1 blocking antibody. Irrespective of the tumor cell, we found an increase in type 1 helper ILCs on PD-1 blockade in both melanoma and metastatic melanoma. These data suggest that within the metastatic melanoma TME, NK1.1^−^ Tbet^+^ ILCs may be regulated by PD-1 based immunotherapeutics.

Several immunotherapeutic strategies exist for the treatment of melanoma but there is a single approved systemic immunotherapeutic regimen for advanced cSCC ^49^. cSCC is characterized by high mutational burden, is frequently seen in patients on immune suppressive drugs and has been reported to have a dysfunctional peritumoral immune response ^50 51, 52^ which suggests that it may respond well to immunotherapy. Recently, the use of PD-1 blocking antibodies has been tested in cSCC patients ^53 49 54^ and has shown significant efficacy in half of the patients treated but the underlying immunobiology is not yet understood. Our data demonstrating that PD-1 has a functional role in ILCs within human cSCC tumor’s suggests that ILCs may play a role in anti-tumor immunity in cSCC patients. Of note is the enrichment of ILC populations within the TME compared to peripheral blood, highlighting the importance of characterizing and investigating immune infiltrates within the tumor tissue in order to understand the role of ILCs when treating patients with immunotherapies in these cancers. Furthermore, since PD-1 controls the frequency of Tbet^+^ ILCs in cSCC, our data suggests a potential new stratification strategy for cSCC patients whereby the frequency of Tbet^+^PD1^+^ ILCs within the TILs might determine the efficacy of anti-PD-1 therapy in this patient population.

In summary, we are the first to demonstrate that PD-1 regulates the frequency and function of Tbet^+^ NK1.1^−^ helper ILCs in a tumor dependent manner. In human cancers, PD1 consistently regulates the proliferation of Tbet^+^RORγt^−^ helper ILCs. It remains to be deciphered to what extent NK1.1^−^Tbet^+^ ILCs can be harnessed for boosting anti-tumor responses by utilizing PD-1 based immunotherapies. However, the data presented in this study clearly highlights the critical need to understand the regulatory function of PD1 on ILC subsets in order to fully harness the potential of PD-1 in modulating immune responses through ILCs.

## Acknowledgements

We would like to thank Prof. Andrew Mellor for critically reading the manuscript and providing insightful comments. The work was funded by an Academy of Medical Sciences Springboard Award supported by the Wellcome Trust. The work was also funded by an MRC project grant, a Leo Foundation Future Leader Award and a Lister Prize. J.X.L is funded by an NU Overseas Research Scholarship award and a NC3R PhD studentship, G.M is funded by an MRC DiMEN PhD studentship. We would like to thank Edyta Kowalczyk (BD Biosciences) for her Bioinformatics expertise, data QC analysis and support for data analysis.

## Author Contributions

J.X.L, C.Y.L, G.E.M, D.M, G.H, S.L, M.P, W.P, M.A, J.C, A.F, performed experiments and analyzed data. A.S performed Bioinformatic analysis, R.P, G.S, E.H and P.E.L critically read the manuscript, provided intellectual input and edited the manuscript. S.A conceptualized the work, analyzed data, provided intellectual input and wrote the paper.

## Declaration of Interests

The authors disclose no conflict of interest.

## Materials and Methods

### Animals

Wild-type C57BL/6 (WT), B6.*Pdcd1*^*−/−*^ (PD-1^−/−^), *B6*.*Pdcd1*^*−/−*^*TbetZsGreen, B6*.*TBetZsGreen* and *B6*.*Rag2*^*−/−*^ littermates were bred and maintained in a pathogen-free facility at Newcastle University under a home office approved project license. All experimental animals were 8-12 weeks of age. All murine experimental procedures were performed at Newcastle University incorporated the NC3R guidelines for animal research and the data were presented according to the ARRIVE guidelines.

### Cell lines

B16F10 melanoma and MC38 adenocarcinoma colon cancer were kindly provided by Dr Arunakumar Gangaplara, NCI, NIH. Tumor cells were cultured in Dulbecco’s Modified Eagle Medium (DMEM) supplemented with 10% FBS (Labtech), sodium pyruvate (1mM; Sigma) and penicillin-streptomycin (100 units/ml penicillin; 100μg/ml streptomycin; Gibco™) and maintained at 37°C with 5% CO2. C8161 human melanoma cell line and MET1 human squamous cell carcinoma (SCC) cell line (Ximbio) were kindly donated by Professor Penny Lovat, Newcastle University. C8161 cells were cultured with DMEM media supplemented with 10% FBS (Labtech), sodium pyruvate (1mM; Sigma) and penicillin-streptomycin (100 units/ml penicillin; 100μg/ml streptomycin; Gibco™). MET1 cells were cultured with DMEM/F12 media (4:1) supplemented with 10% FBS (Labtech), penicillin streptomycin (100units/ml penicillin; 100μg/ml streptomycin), hydrocortisone (0.4μg/ml; Sigma), cholera toxin (8.5ng/ml; Sigma), tri-iodo-L-threonine (20pM; Sigma), Adenine (180μM; Sigma), Insulin (5μg/ml; Sigma), epithelial growth factor (EGF; 2pg/ml; Sigma) and transferrin (5μg/ml; Sigma).

### Animal Models

#### Subcutaneous tumor models of B16F10 and MC38

WT or B6.*Pdcd1*^*−/−*^, B6.*Pdcd1*^*−/−*^TbetZsGreen, B6.TbetZsGreen, B6.*Rag2*^−/−^ mice were inoculated with 2 × 10^5^ B16F10 melanoma cells or MC38 cells via subcutaneous injection into the flank as previously reported ^55^. Tumor volume was recorded daily from day 5 post inoculation. Tumor volume was calculated by the following equation: Tumor volume = π/6 × 0.5 x length x width^2^ as described previously ^56^. In certain experiments, mice were treated with anti-PD-1 (200μg/mouse; clone: RPMI-40); anti-NK1.1 (200μg/mouse; clone:PK136) or isotype control (IgG2a, 200μg/mouse, clone: 2A3; IgG1a, 200μg/mouse; C1.18.24 respectively) via intraperitoneal injection at day 7, 9 and 11 unless otherwise indicated. Mice were euthanized at day 12, unless otherwise stated for tumor infiltrating lymphocyte (TIL) immunobiology assays. In some experiments, animals were injected with BODIPY (200μg/mouse) or vehicle before harvest.

#### Metastatic melanoma

WT or *B6*.*TbetZsGreen or B6*.*Pdcd1*^*−/−*^ mice were inoculated with either 0.5 or 2 × 10^5^ B16F10 melanoma cells in 200μl of PBS via intravenous tail-vein injection. Mice were monitored for clinical signs of weight loss and euthanized at >25% weight loss. In certain experiments mice were treated with anti-PD-1 (200μg/mouse; clone: RPMI-40) or isotype control (IgG2a, 200μg/mouse; clone: 2A3) at day 7, 9, 11. Tissue was harvested at day 12 and analyzed for the presence of ILCs unless stated otherwise.

#### AOM-DSS induced colorectal cancer

The AOM-DSS model was set up as previously reported ^57^. Briefly, mice were treated with one dose of Azoxymethane (AOM; 12mg/kg) at day 1 and then 3% dextran sodium sulphate (DSS) was added as drinking water from day 3 for a week. Mice were allowed to recover for 3 weeks and then a 2^nd^ cycle of DSS was started. After the 3^rd^ cycle, animals were euthanized and immunobiology studied.

### ILC isolation from tumor and normal tissue

#### TIL isolation

ILCs from the tumor tissue were isolated as previously described ^58^. Briefly, tumor tissue was incubated at 37°C for 30minutes in FBS free DMEM media containing liberase™ TL (0.25mg/ml; Roche) and DNAse I (0.5mg/ml; Roche). Single-cell suspensions were prepared by mechanically disrupting tissue through a 100μm nylon cell strainer into FBS. Lymphocytes were isolated using lymphocyte separation media (LSM; Promocell) and washed twice with complete media (DMEM supplemented with 10% FBS, glutamine (2mM), non-essential amino acids (0.1mM), sodium pyruvate (1mM), 2-mercaptoethanol (50μM) and penicillin and streptomycin (100 units/ml penicillin; 100μg/ml streptomycin; Gibco™) in order to remove traces of LSM. Cells were then analyzed by flow cytometry or stimulated to induce cytokine production. Ethical approval for experiments conducted on human tissue was provided by the South Central Hampshire B NRES Committee (reference number 07/H0504/187). Fresh tissue samples of cSCC and non-lesional skin were obtained from patients during surgery at the Dermatology Department, University Hospital Southampton NHS Foundation Trust. For lymphocyte isolation, samples of tumor and separately, non-lesional skin, were finely disaggregated with scalpels, incubated at 37°C for 1.5 hours in RPMI media containing collagenase I-A (1mg/ml; Sigma-Aldrich) and DNAse I (10µg/ml; Sigma-Aldrich). The resulting suspension was then passed through a 70µm cell strainer and centrifuged over an Optiprep (Sigma-Aldrich) density gradient. Lymphocytes were then extracted and washed with PBS before use in experiments. In certain experiments, TILs were incubated for 60 mins with 2NBDG (200μg/ml) or PBS and then ILCs analyzed by flow cytometry.

#### ILC isolation from spleen

Single cell-suspension of splenocytes was generated by mechanically disrupting tissue through a 40μm filter into complete media. Cells were incubated with red blood cell (RBC) lysis buffer (Biolegend) for 3 minutes at room temperature and were then washed in complete media once before analysis by flow cytometry.

#### ILC isolation from tumor draining lymph nodes

Tumor draining lymph nodes (TDLN) were isolated from the inguinal draining lymph nodes. Single cell-suspension was generated by mechanically disrupting TDLNs through a 40μm filter into complete media. Cells were then washed and analyzed by flow cytometry.

#### ILC isolation from lungs

Lungs were perfused with PBS *in situ* via the pulmonary artery prior to isolation. Tissue was incubated at 37°C for 15 minutes in FBS free DMEM media containing Liberase™ TL (0.25mg/ml; Roche) and DNAse I (0.5mg/ml; Roche). Single cell suspension was generated by mechanically disrupting tissues through a 100µm filter into FBS followed by a Percoll gradient centrifugation (40% Percoll, GE Healthcare; FBS free media containing 0.5mg/ml DNase, Roche). Cells were then washed with complete media and analyzed by flow cytometry.

#### ILC isolation from small intestine

Small intestine tissue was harvested into complete media. Fecal matter was removed, and tissue was washed in buffer (PBS containing 5% FBS, Hepes (1mM; Sigma), 50μM 2-mercaptoethanol (50μM; Sigma) and penicillin and streptomycin (100 units/ml penicillin; 100μg/ml streptomycin; Gibco™)).This was followed by a wash with PBS to remove traces of FBS. Tissue was incubated at 37°C for 30 minutes in FBS free DMEM media containing Liberase™ TL (0.25mg/ml; Roche) and DNAse I (0.5mg/ml; Roche). Single cell suspension was generated by mechanically disrupting tissues through a 100µm filter into FBS. This was followed by percoll gradient centrifugation (40% Percoll (GE Healthcare), containing 0.5mg/ml DNase (Roche)). Cells were washed with complete media and analyzed by flow cytometry.

#### ILC isolation from skin

Mouse normal dorsal skin was harvested, finely chopped and incubated for 2 hours in FBS free DMEM media containing Liberase™ TL (0.25mg/ml; Roche) and DNAse I (0.5mg/ml; Roche) at 37°C. Single-cell suspensions were prepared by mechanically crushing tissue slurry through a 100μm nylon mesh into FBS. Cells were then subsequently filtered through 70- and 40-μm nylon filters. Cells were then analyzed via flow cytometry.

### Antibodies

All the antibodies used to characterize murine and human ILCs were purchased from BioLegend or eBioscience unless otherwise stated. For analysis of murine cell surface markers, the following antibodies were used: Lineage consisted of CD3 (clone: 1452C11), CD5 (clone: 53-7.3), CD8 (clone: 53-6.7), CD11b (clone: M1170), CD11c (clone: N418), CD19 (clone: 1D3/CD19), CD49b (clone: HMα2), Ter119 (clone: TER-119), Gr1 (clone: RB8-9C5), F4/80 (clone:BM8), Nk1.1 (clone:PK136), B220 (clone:RA3-8B2). ILCs were stained for CD45 (clone:30-F11), CD90.2 (clone: 30-H12), CD127 (clone: A7R34), CD25 (clone: PC61), KLRG1 (clone:2F1/KLRG1), Nkp46 (clone:29A1.4), CD49a (clone: HMα1), PD-1 (clone: RPMI-30), PDL-1 (clone:10F.9G2), PDL-2 (clone: Ty25). ST2 (clone:DJ8) was purchased from MD Bioproducts. For ILC analysis in human peripheral blood mononuclear cells (PBMC), the following fluorochrome-conjugated antibodies were used: Lineage consisting of CD3 (clone: OKT3), CD5 (clone: L17F12), CD14 (clone: M5E2), CD16 (HIB19), CD19 (HIB19), CD20 (2H7), CD56 (clone HCD56), CD11b (clone:ICRF44), CD11c (clone: 3.9) and TCRα/β (clone: IP26), CD45 (clone:H130), CD127 (clone:A019D5), CD161 (clone:HP-3G10), c-kit (CD117;clone:104D2), CRTH2 (clone:BM16), PD-1 (EH12.2H7) and Nkp44 (clone:P44-8). For analysis of murine cytokine production, the following fluorochrome-conjugated antibodies were used: IL-5 (clone: TRFK5), IL-10 (clone: JES5-16ES), TNFα (clone: MP6-XT22), IFNγ (clone: XMG1.1), RORγt (clone: B2D), EOMES (clone: Dan11mag), IL-17 (clone: eBio17B7), IL-22 (clone: IL22JOP), IL-13 (clone: eBio13A) and Ki67 (clone: So1A15). Human PBMCs were stained with the following intracellular fluorochrome-conjugated antibodies; Tbet (clone: 4B10), IFNγ (clone: 4S.B3), IL-17 (clone: BL168), IL-5 (clone: TRFK5), TNFα (clone: Mab11), RORγt (clone: B2D) and IL-13 (clone: JES10-5A2).

### Flow cytometry

Single cell suspensions were generated from indicated organs and stained with Live/Dead fixable dead cell stain kit as per manufacturer’s instructions (Invitrogen). For murine ILC analysis, cells were incubated with biotin labeled lineage cocktail (CD3^+^, CD5^+^, CD8^+^, CD11b^+^, CD11c^+^, CD19^+^, CD49b^+^, Ter119^+^, F4/80^+^, B220^+^, NK1.1^+^ and Gr1^+^) followed by streptavidin. Cells were then stained with a combination of markers including CD45, CD90.2, CD127, CD25, KLRG1, Nkp46, PD-1, PDL-1, PDL-2 and ST2. For analysis of NK and myeloid immune subsets, TILs were stained with NK1.1, CD49a, CD49b, F4/80, Gr1, CD11b and CD11c. Cells were then fixed and permeabilized for intracellular markers as follows: Tbet, RORγ, Ki67 and EOMES. In order to measure murine intracellular cytokine (IC) production, TILs were stimulated with cytokine stimulation cocktail (Invitrogen; Thermo Fisher Scientific) for 4 hours at 37°C. Cells were then fixed and permeabilized (Fixation/Permeabilization kit; BD Bioscience). ILCs were then stained with: IL-5, IL-13, IL-17, IL-22, IFN-γ and TNF-α.

Human PBMCs and TILs were washed with PBS prior to staining. 1 × 10^6^ cells were stained with Live/Dead fixable dead stain kit as per manufactures instructions (Invitrogen). Cells were then incubated with cell surface antibodies: Lineage cocktail BV510 or FITC (CD3^+^, CD5^+^, CD11b^+^, CD11c^+^, CD14^+^, CD16^+^, CD19^+^, CD20^+^, CD56^+^ and TCRα/β^+^), CD45, CD127, CD161, CRTH2, c-Kit, Nkp44 and PD-1. Cells were then fixed and permeabilized (Fixation/Permeabilization kit; BD Bioscience) and stained for intracellular transcription factors as follows: Tbet and RORγt.

ILCs were defined by the following gating strategies: Murine ILCs were defined as Lin^−^ Thy1^+^; ILC2s were defined as CD127^+^CD25^+^KLRG1^−/−^ST2^−/−^; NCR^+^ ILC3s were defined as RORγt^+^NKp46^+^ and NCR^−^ ILC3s were defined as RORγt^+^Nkp46^−^, murine ILC subset regulated by PD1 was defined as Lin^−^ Thy1^+^ NK1.1^−^Tbet^+^NKp46^-.^ Human ILCs were defined as follows: Lin^−^ CD45^+^CD161^+^CD127^+^. ILC1s were further defined as CRTH2^−^CD117^−^; ILC2s were defined as CRTH2^+^CD117^−^; ILC3s were defined as CRTH2^−^CD117^+^. Cells were analyzed using BD LSR Fortessa X20 with FACs DIVA software (BD Bioscience) and analysis was performed with FCS Express (De Novo) or FlowJo 10.1 software (Tree Star).

### Rhapsody Single Cell Sequencing

TILs were isolated as previously described from tumours and then cells were stained with lineage markers (lineage gate included CD3^+^, CD4^+^, CD5^+^, CD8^+^, CD11b^+^, CD11c^+^, CD19^+^, CD49b^+^, Ter119^+^, F4/80^+^, B220^+^ and Gr1^+^), Thy1, and Abseq antibodies and sample Tags. Abseq antibody-oligos used were as follows: CD25, CD103, CD119, CD37, CD223, CD272, CD273, CD274, CD278, CD279, IL17Rb, IL23R, IL33R, CD335, CD3 and NK1.1. Cells were incubated for 20 minutes at 4C and then washed three times with Miltenyi buffer. TILs were then stained with DAPI and flow sorted for Lineage^−^Thy1^+^ population. Samples were then pooled and loaded on to rhapsody cartridges and then experiment were performed as per manufacturer’s instructions. Data analysis was performed using the SeqGeq software.

### Transwell assays

Splenocytes were RBC lysed and then plated at a concentration of 0.5 × 10^6^ cells per ml. Transwell inserts (0.4 micron; ThermoFisher Scientific) were seeded with B16F10 melanoma cells at a 1:1 ratio with splenocytes (unless otherwise stated) and were incubated for indicated time points at 37°C prior to flow cytometry analysis. For proliferation assays, transwell inserts were removed after 6 hours. For human experiments, PBMCs were acquired from healthy donors and were cultured at a concentration of 0.5 × 10^6^ per ml. Transwell inserts were seeded with either C8161 human melanoma cell line or human cSCC cell line at a 1:1 ratio with PBMCs. For human experiments, transwell inserts were removed after 16 hours. Plates were incubated at 37°C for indicated time points and were then analyzed by flow cytometry.

### *In-vitro* Proliferation Assays

For cell trace violet experiments, murine splenocytes isolated from B6.TbetZsGreen mice were stained in PBS with Cell Trace Violet (Invitrogen) as per manufactures instructions. Cells were cultured with IL-2 (40ng/ml), IL-7 (40ng/ml), αPD-1 (20μg/ml; clone: RMP1-14) or Isotype IgG2a (20μg/ml; clone: 2A3) as indicated for 5 days in cell culture media (DMEM supplemented with 10% FBS, glutamine (2mM), non-essential amino acids (0.1mM), sodium pyruvate (1mM), 2-mercaptoethanol (50μM), penicillin and streptomycin (100 U/M)). Cytokines were replenished on day 2 and day 4. Proliferation was measured on day 5 by flow cytometry. For human proliferation assays, human PBMCs acquired from healthy donors were stained in PBS with Cell Trace Violet (Invitrogen) as per manufacturer’s instructions. Cells were cultured with IL-2 (40ng/ml), IL-7 (40ng/ml), αPD-1 (20μg/ml; clone:EH12.2H7) or Isotype IgG1 (20μg/ml; clone: MG1-45) as indicated for 7 days in cell culture media (RPMI supplemented with 10% FBS, glutamine (2mM), non-essential amino acids (0.1mM), sodium pyruvate (1mM), 2-mercaptoethanol (50μM), penicillin and streptomycin ((100 units/ml penicillin; 100μg/ml streptomycin; Gibco™)). Cytokines were replenished on day 2 and day 4. Proliferation was measured on day 7 by flow cytometry.

### Lactate Assays

WT or PD1^−/-^ splenocytes were incubated with IL2 (100 ng/ml) plus IL7 (100 ng/ml) alone or in combination with lactic acid (20 mM) for 24 hrs and then PD1 expression was measured by flow cytometry. B16F10 tumor cells were expanded and then supernatant was tested for lactic acid production as per the manufacturer’s instructions (Abcam). Briefly, 2×10^6^ cells were seeded in 24 well plates and then supernatant harvested after 4hrs or 24 hrs. The amount of lactic acid was determined using a lactic acid fluorometry kit

### Phospho-P70S6Kinase Measurement

Tumors were resected when they reached >600mm^3^ and then TILs were isolated. TILs were stimulated with IL2 (80ng/ml) and IL-7 (40ng/ml) for 15 minutes. TILs were washed once with PBS and then stained for phosphoP70S6Kinase antibody and then analyzed by flow cytometry. In some experiments, TILs were enriched using CD90.2 microbeads as per manufacturer’s instructions and then cultured for 3 days with IL2 plus IL7 alone or in combination with isotype (20μg/ml) or anti-PD1 antibody (20μg/ml). At day 3 post cultures, TILs were washed with complete media once and then restimulated with IL2 and IL7 for 15mins. Following stimulation, phosphorylation of P70S6Kinase was measured.

### Statistical Analysis

Statistical analysis was performed with GraphPad Prism using an unpaired student two-tailed T test for groups of two and a ONE-WAY ANOVA for multiple groups. Results are expressed as mean± standard error of the mean (SEM) and P-values ≤0.05 were considered significant. Survival curve analysis was performed using a Kaplan-Meier survival curve and a log rank test.

## Notes

### Competing Interest Statement

The authors have declared no competing interest.

## References

1. Spits, H. et al. Innate lymphoid cells--a proposal for uniform nomenclature. Nat Rev Immunol 13, 145–149 (2013).

2. Bernink, J.H. et al. Human type 1 innate lymphoid cells accumulate in inflamed mucosal tissues. Nat Immunol 14, 221–229 (2013).

3. Fuchs, A. et al. Intraepithelial type 1 innate lymphoid cells are a unique subset of IL-12- and IL-15-responsive IFN-gamma-producing cells. Immunity 38, 769–781 (2013).

4. Klose, C.S.N. et al. Differentiation of type 1 ILCs from a common progenitor to all helper-like innate lymphoid cell lineages. Cell 157, 340–356 (2014).

5. Daussy, C. et al. T-bet and Eomes instruct the development of two distinct natural killer cell lineages in the liver and in the bone marrow. J Exp Med 211, 563–577 (2014).

6. Peng, H. et al. Liver-resident NK cells confer adaptive immunity in skin-contact inflammation. J Clin Invest 123, 1444–1456 (2013).

7. Mjosberg, J.M. et al. Human IL-25- and IL-33-responsive type 2 innate lymphoid cells are defined by expression of CRTH2 and CD161. Nat Immunol 12, 1055–1062 (2011).

8. Monticelli, L.A. et al. Innate lymphoid cells promote lung-tissue homeostasis after infection with influenza virus. Nat Immunol 12, 1045–1054 (2011).

9. Moro, K. et al. Innate production of T(H)2 cytokines by adipose tissue-associated c-Kit(+)Sca-1(+) lymphoid cells. Nature 463, 540–544 (2010).

10. Neill, D.R. et al. Nuocytes represent a new innate effector leukocyte that mediates type-2 immunity. Nature 464, 1367–1370 (2010).

11. Cella, M. et al. A human natural killer cell subset provides an innate source of IL-22 for mucosal immunity. Nature 457, 722–725 (2009).

12. Sonnenberg, G.F., Monticelli, L.A., Elloso, M.M., Fouser, L.A. & Artis, D. CD4(+) lymphoid tissue-inducer cells promote innate immunity in the gut. Immunity 34, 122–134 (2011).

13. Sawa, S. et al. Lineage relationship analysis of RORgammat+ innate lymphoid cells. Science 330, 665–669 (2010).

14. Satoh-Takayama, N. et al. Microbial flora drives interleukin 22 production in intestinal NKp46+ cells that provide innate mucosal immune defense. Immunity 29, 958–970 (2008).

15. Buonocore, S. et al. Innate lymphoid cells drive interleukin-23-dependent innate intestinal pathology. Nature 464, 1371–1375 (2010).

16. Sanos, S.L. et al. RORgammat and commensal microflora are required for the differentiation of mucosal interleukin 22-producing NKp46+ cells. Nat Immunol 10, 83–91 (2009).

17. Cupedo, T. et al. Human fetal lymphoid tissue-inducer cells are interleukin 17-producing precursors to RORC+ CD127+ natural killer-like cells. Nat Immunol 10, 66–74 (2009).

18. Crellin, N.K. et al. Regulation of cytokine secretion in human CD127(+) LTi-like innate lymphoid cells by Toll-like receptor 2. Immunity 33, 752–764 (2010).

19. Gao, Y. et al. Tumor immunoevasion by the conversion of effector NK cells into type 1 innate lymphoid cells. Nat Immunol 18, 1004–1015 (2017).

20. Jovanovic, I.P. et al. Interleukin-33/ST2 axis promotes breast cancer growth and metastases by facilitating intratumoral accumulation of immunosuppressive and innate lymphoid cells. Int J Cancer 134, 1669–1682 (2014).

21. Li, J. et al. Biliary repair and carcinogenesis are mediated by IL-33-dependent cholangiocyte proliferation. J Clin Invest 124, 3241–3251 (2014).

22. Ikutani, M. et al. Identification of innate IL-5-producing cells and their role in lung eosinophil regulation and antitumor immunity. J Immunol 188, 703–713 (2012).

23. Saranchova, I. et al. Type 2 Innate Lymphocytes Actuate Immunity Against Tumours and Limit Cancer Metastasis. Sci Rep 8, 2924 (2018).

24. Wagner, M. et al. Tumor-Derived Lactic Acid Contributes to the Paucity of Intratumoral ILC2s. Cell Rep 30, 2743–2757 e2745 (2020).

25. Moral, J.A. et al. ILC2s amplify PD-1 blockade by activating tissue-specific cancer immunity. Nature 579, 130–135 (2020).

26. Kirchberger, S. et al. Innate lymphoid cells sustain colon cancer through production of interleukin-22 in a mouse model. J Exp Med 210, 917–931 (2013).

27. Chan, I.H. et al. Interleukin-23 is sufficient to induce rapid de novo gut tumorigenesis, independent of carcinogens, through activation of innate lymphoid cells. Mucosal Immunol 7, 842–856 (2014).

28. Eisenring, M., vom Berg, J., Kristiansen, G., Saller, E. & Becher, B. IL-12 initiates tumor rejection via lymphoid tissue-inducer cells bearing the natural cytotoxicity receptor NKp46. Nat Immunol 11, 1030–1038 (2010).

29. Nussbaum, K. et al. Tissue microenvironment dictates the fate and tumor-suppressive function of type 3 ILCs. J Exp Med 214, 2331–2347 (2017).

30. Salimi, M. et al. Activated innate lymphoid cell populations accumulate in human tumour tissues. BMC Cancer 18, 341 (2018).

31. Taylor, S. et al. PD-1 regulates KLRG1(+) group 2 innate lymphoid cells. J Exp Med 214, 1663–1678 (2017).

32. Jacquelot, N. et al. Blockade of the co-inhibitory molecule PD-1 unleashes ILC2-dependent antitumor immunity in melanoma. Nat Immunol 22, 851–864 (2021).

33. Hsu, J. et al. Contribution of NK cells to immunotherapy mediated by PD-1/PD-L1 blockade. J Clin Invest 128, 4654–4668 (2018).

34. Patsoukis, N. et al. PD-1 alters T-cell metabolic reprogramming by inhibiting glycolysis and promoting lipolysis and fatty acid oxidation. Nat Commun 6, 6692 (2015).

35. Strauss, L. et al. Targeted deletion of PD-1 in myeloid cells induces antitumor immunity. Sci Immunol 5 (2020).

36. Helou, D.G. et al. PD-1 pathway regulates ILC2 metabolism and PD-1 agonist treatment ameliorates airway hyperreactivity. Nat Commun 11, 3998 (2020).

37. Vely, F. et al. Evidence of innate lymphoid cell redundancy in humans. Nat Immunol 17, 1291–1299 (2016).

38. Lim, A.I. et al. Systemic Human ILC Precursors Provide a Substrate for Tissue ILC Differentiation. Cell 168, 1086–1100 e1010 (2017).

39. Nagasawa, M. et al. KLRG1 and NKp46 discriminate subpopulations of human CD117(+)CRTH2(-) ILCs biased toward ILC2 or ILC3. J Exp Med 216, 1762–1776 (2019).

40. Mazzurana, L. et al. Tissue-specific transcriptional imprinting and heterogeneity in human innate lymphoid cells revealed by full-length single-cell RNA-sequencing. Cell Res 31, 554–568 (2021).

41. Ji, A.L. et al. Multimodal Analysis of Composition and Spatial Architecture in Human Squamous Cell Carcinoma. Cell (2020).

42. Luci, C. et al. Cutaneous Squamous Cell Carcinoma Development Is Associated with a Temporal Infiltration of ILC1 and NK Cells with Immune Dysfunctions. J Invest Dermatol 141, 2369–2379 (2021).

43. Gordon, S.R. et al. PD-1 expression by tumour-associated macrophages inhibits phagocytosis and tumour immunity. Nature 545, 495–499 (2017).

44. Kim, J. et al. Intratumorally Establishing Type 2 Innate Lymphoid Cells Blocks Tumor Growth. J Immunol 196, 2410–2423 (2016).

45. Long, A. et al. Type 2 Innate Lymphoid Cells Impede IL-33-Mediated Tumor Suppression. J Immunol 201, 3456–3464 (2018).

46. Cho, M.M., Quamine, A.E., Olsen, M.R. & Capitini, C.M. Programmed cell death protein 1 on natural killer cells: fact or fiction? J Clin Invest 130, 2816–2819 (2020).

47. Judge, S.J. et al. Minimal PD-1 expression in mouse and human NK cells under diverse conditions. J Clin Invest 130, 3051–3068 (2020).

48. Wilhelm, C. et al. Critical role of fatty acid metabolism in ILC2-mediated barrier protection during malnutrition and helminth infection. J Exp Med 213, 1409–1418 (2016).

49. Migden, M.R. et al. PD-1 Blockade with Cemiplimab in Advanced Cutaneous Squamous-Cell Carcinoma. N Engl J Med 379, 341–351 (2018).

50. Lai, C. et al. OX40+ Regulatory T Cells in Cutaneous Squamous Cell Carcinoma Suppress Effector T-Cell Responses and Associate with Metastatic Potential. Clin Cancer Res 22, 4236–4248 (2016).

51. Shapanis, A. et al. Identification of proteins associated with development of metastasis from cutaneous squamous cell carcinomas (cSCCs) via proteomic analysis of primary cSCCs. Br J Dermatol 184, 709–721 (2021).

52. Lai, C. et al. CD8+CD103+ tissue-resident memory T cells convey reduced protective immunity in cutaneous squamous cell carcinoma. J Immunother Cancer 9 (2021).

53. van Baar, M.L.M., Guminski, A.D., Ferguson, P.M. & Martin, L.K. Pembrolizumab for cutaneous squamous cell carcinoma: Report of a case of inoperable squamous cell carcinoma with complete response to pembrolizumab complicated by granulomatous inflammation. JAAD Case Rep 5, 491–494 (2019).

54. Falchook, G.S. et al. Responses of metastatic basal cell and cutaneous squamous cell carcinomas to anti-PD1 monoclonal antibody REGN2810. J Immunother Cancer 4, 70 (2016).

55. Stathopoulou, C. et al. PD-1 Inhibitory Receptor Downregulates Asparaginyl Endopeptidase and Maintains Foxp3 Transcription Factor Stability in Induced Regulatory T Cells. Immunity 49, 247–263 e247 (2018).

56. Schatton, T. et al. Identification of cells initiating human melanomas. Nature 451, 345–349 (2008).

57. Parang, B., Barrett, C.W. & Williams, C.S. AOM/DSS Model of Colitis-Associated Cancer. Methods Mol Biol 1422, 297–307 (2016).

58. Mallett, G., Patterson, W., Payne, M. & Amarnath, S. Isolation and Characterization of Innate Lymphoid Cells within the Murine Tumor Microenvironment. Methods Mol Biol 2121, 153–164 (2020).

