## Supplementary figures and images for "Programmed cell death-1 receptor mediated regulation of Tbet^+^ NK1.1^−^ Innate Lymphoid Cells within the Tumor Microenvironment"

### Supplemental Figure 1

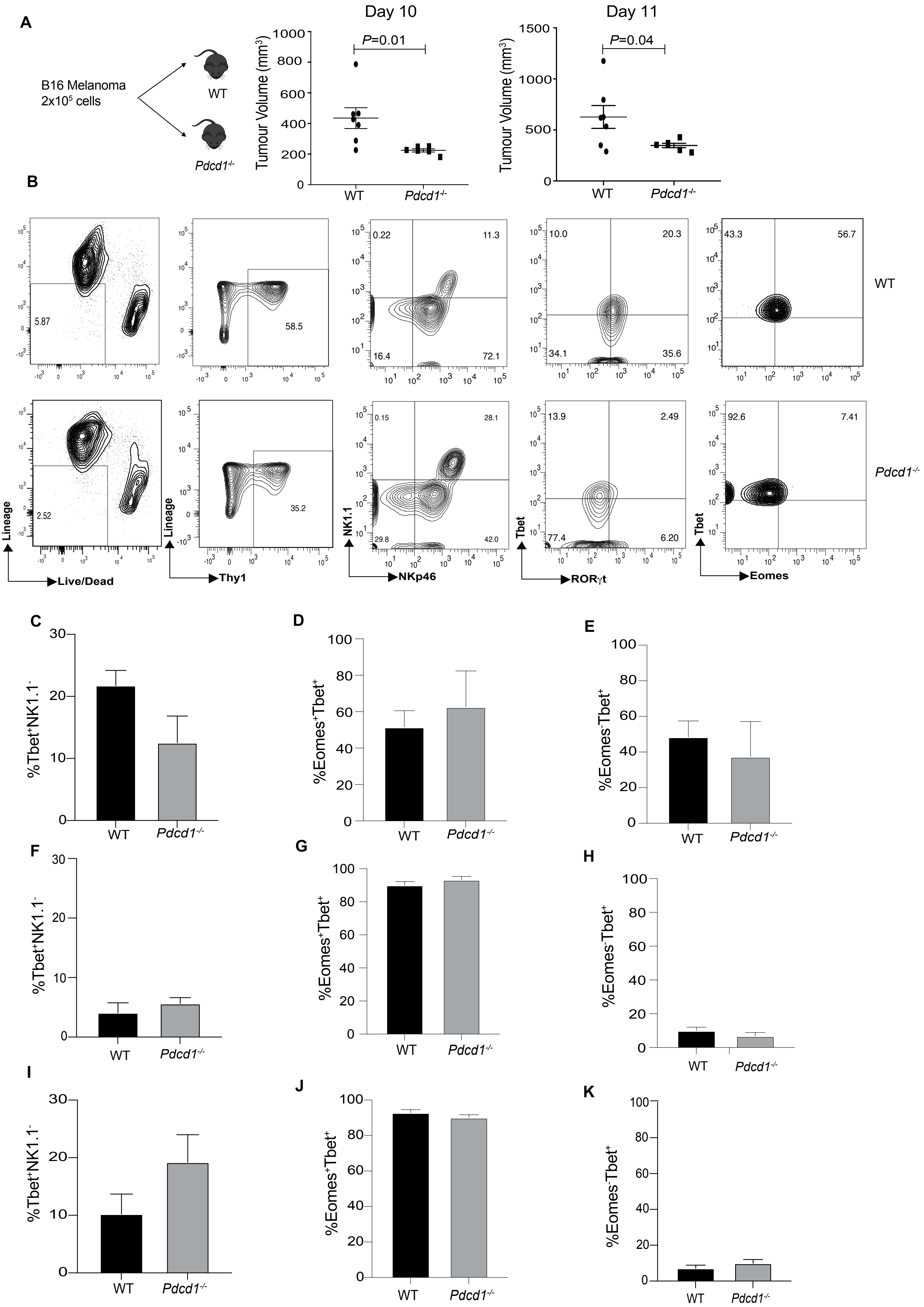

### Supplemental Figure 4

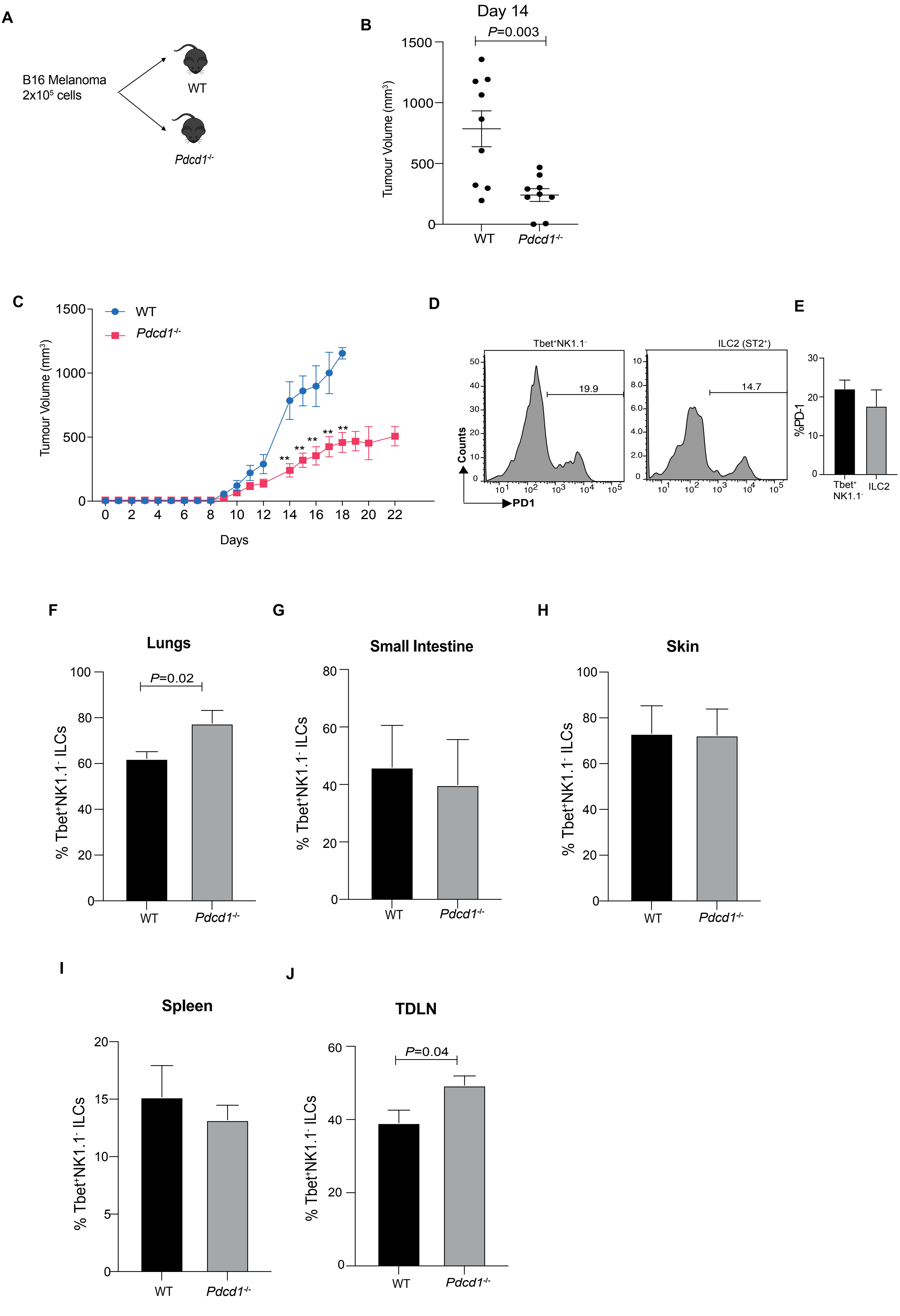

### Supplemental Figure 5

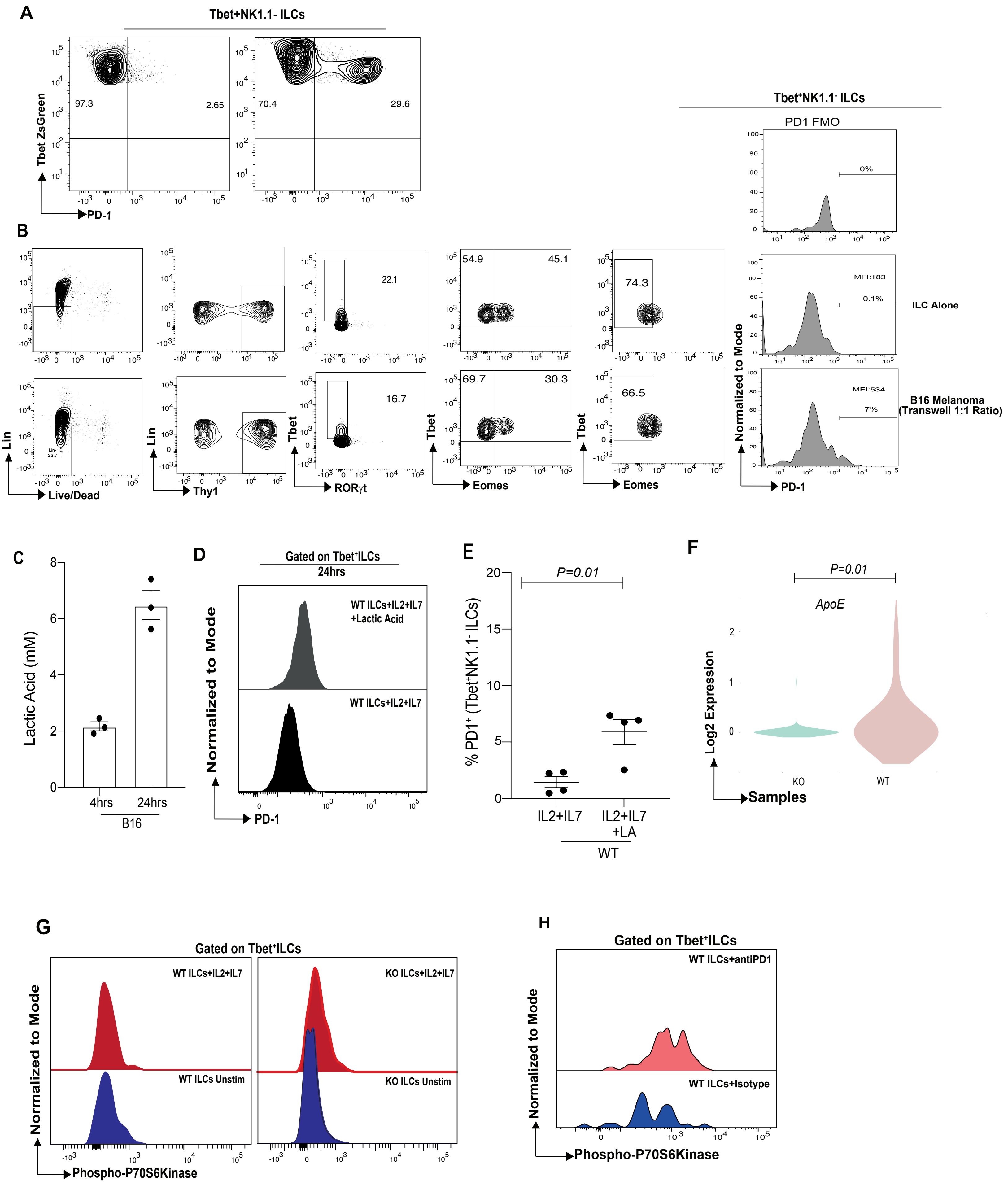

### Supplemental Figure 6

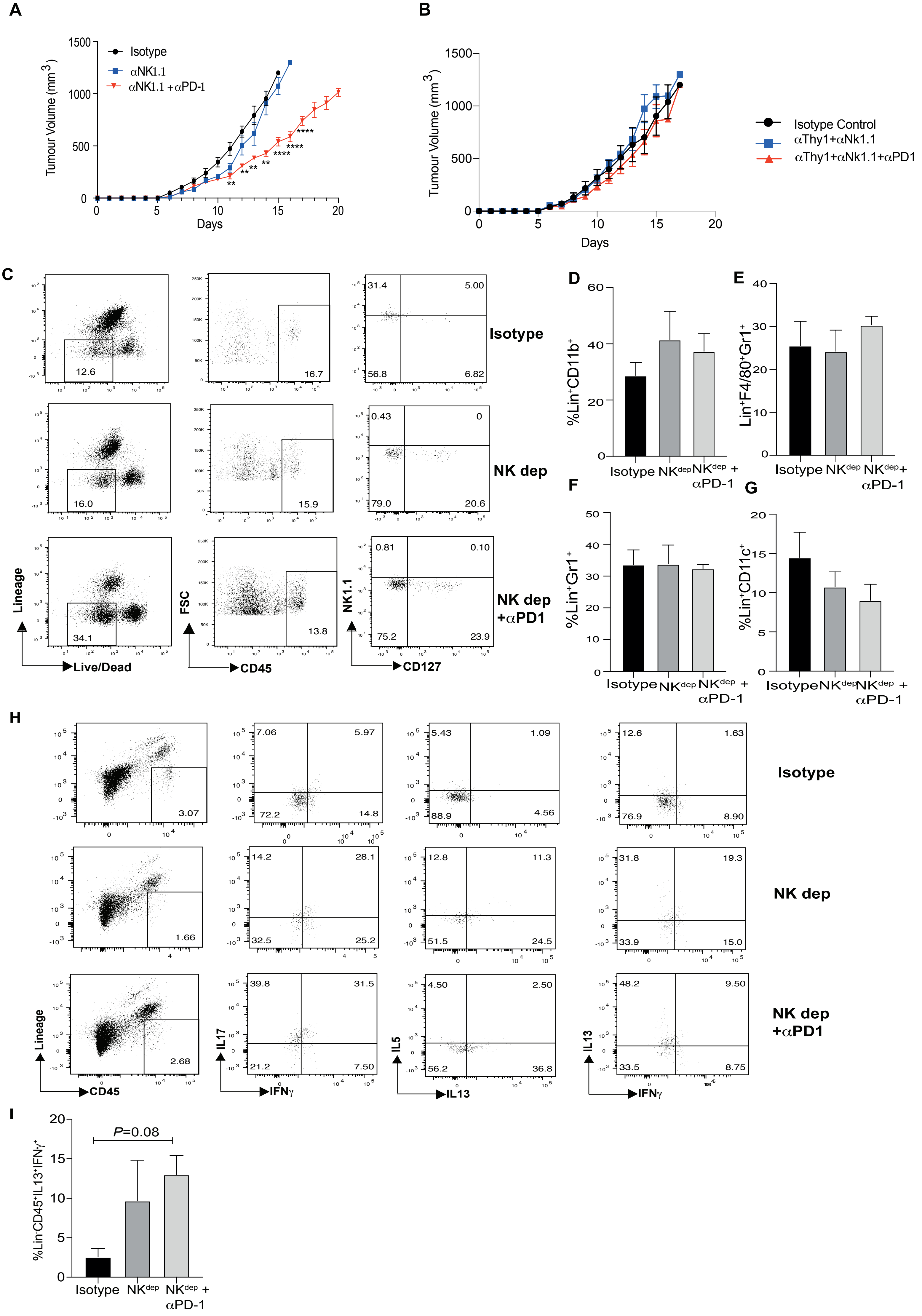

### Supplemental Figure 7

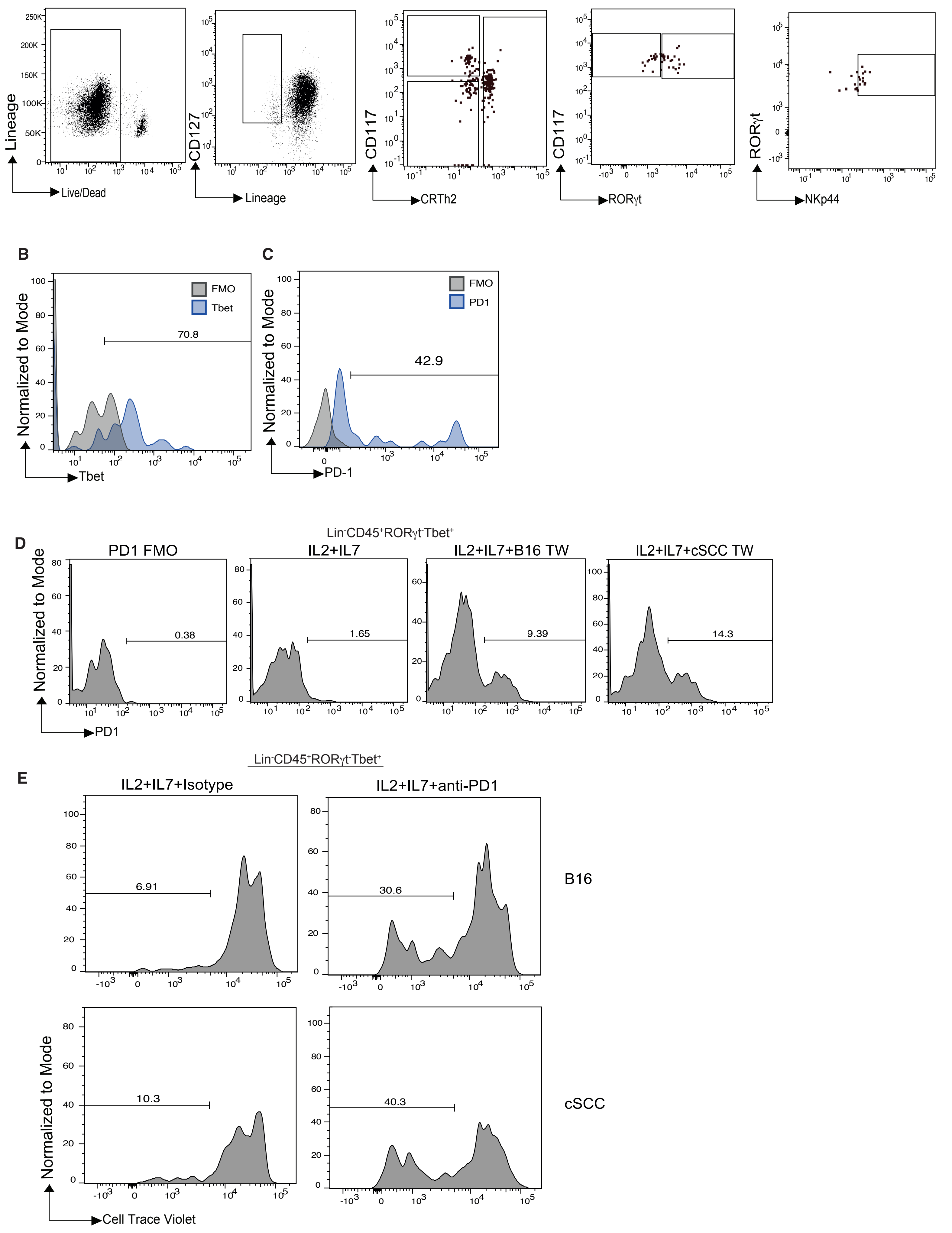
