## Supplemental Figure 2 for "Programmed cell death-1 receptor mediated regulation of Tbet^+^ NK1.1^−^ Innate Lymphoid Cells within the Tumor Microenvironment"

### Pathway Analysis of Clusters

#### Cluster_1

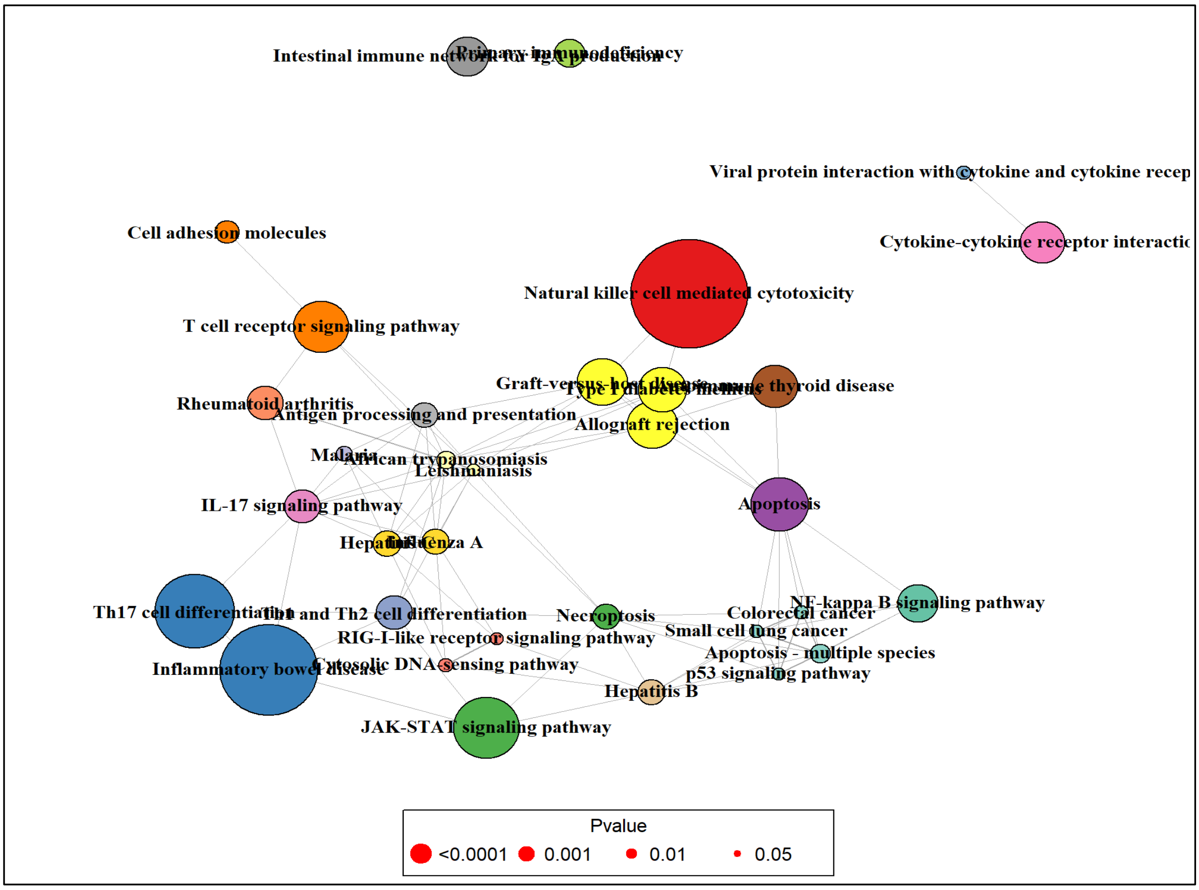


#### Cluster_2

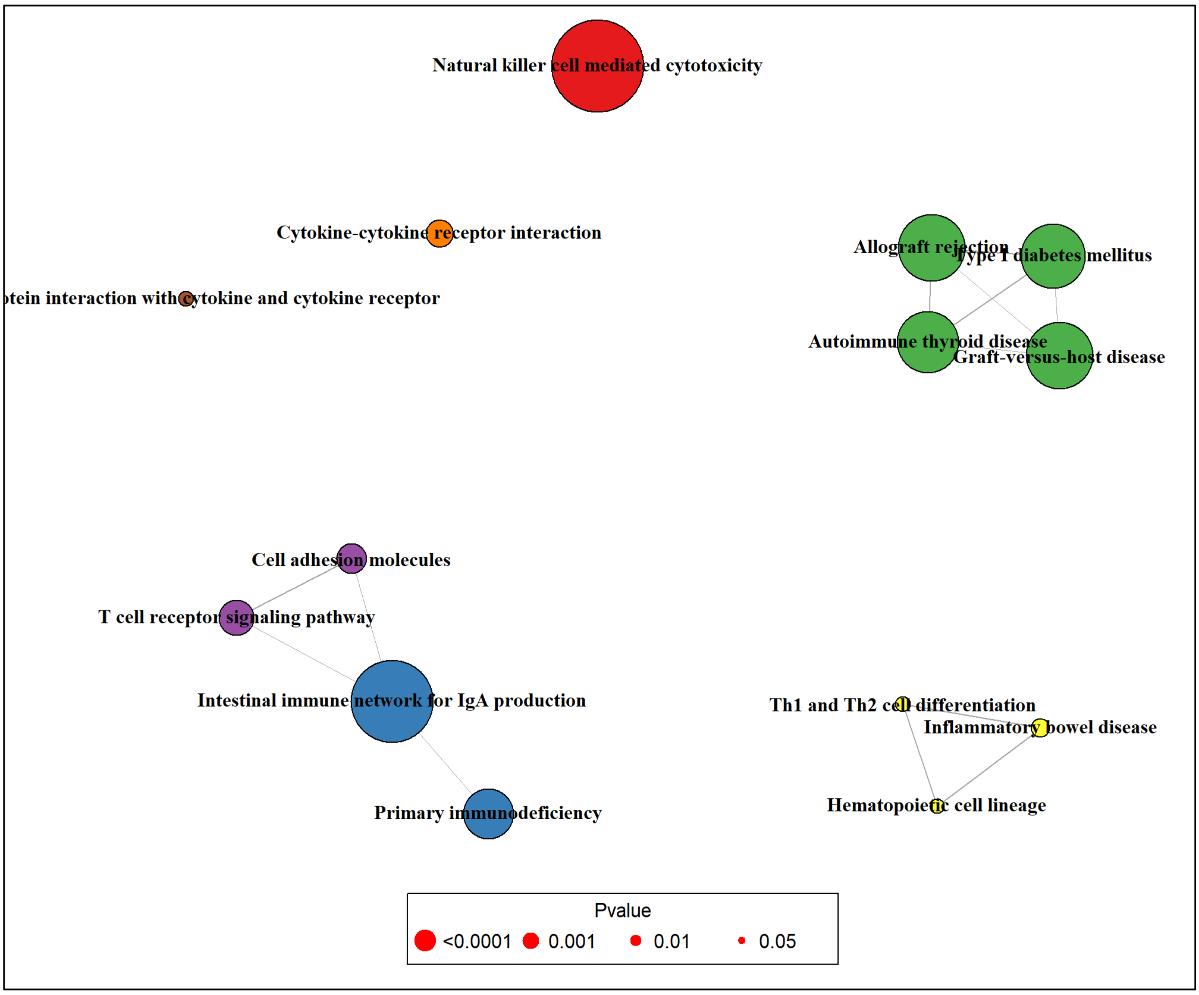

### Cluster_3

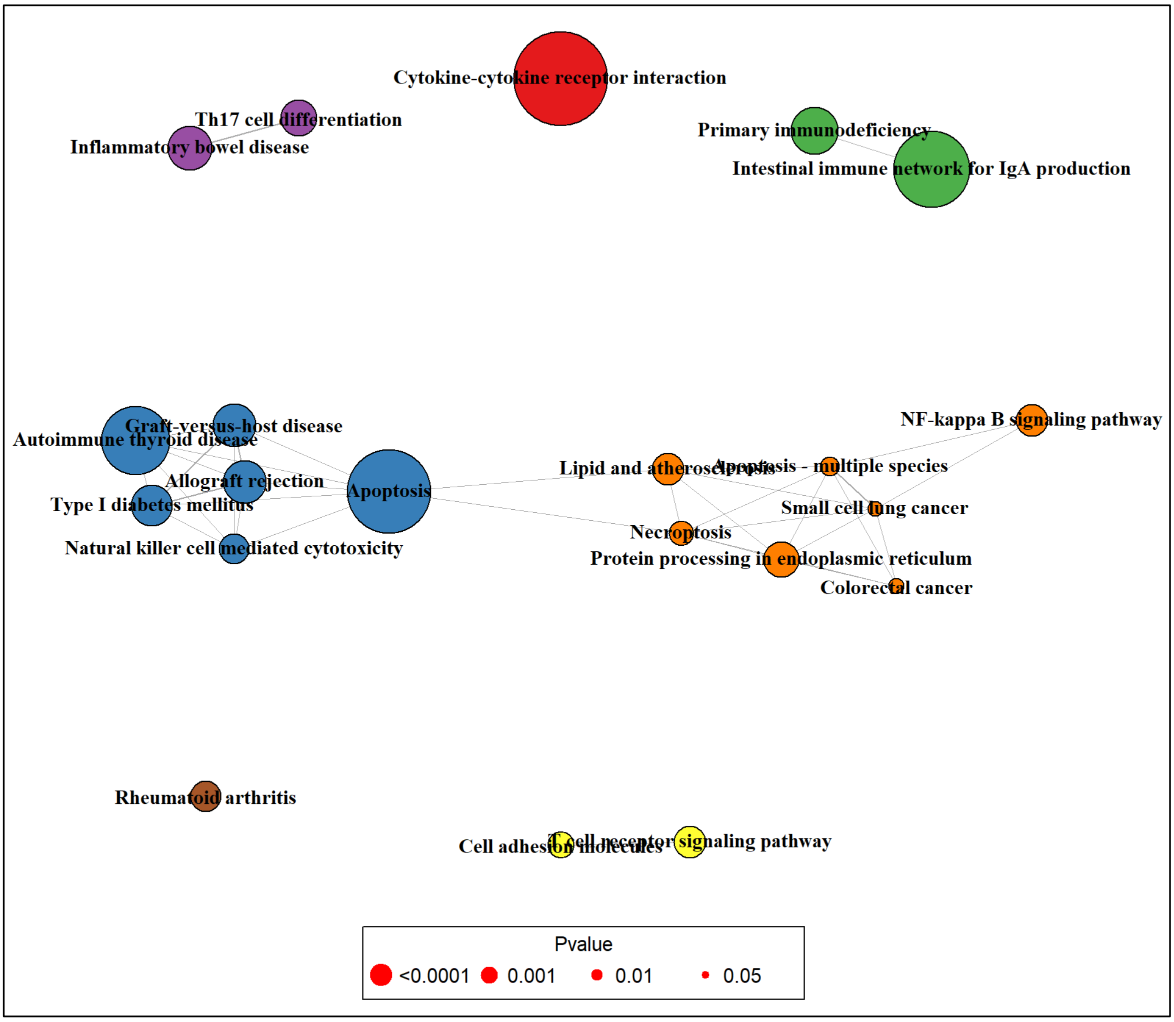


### Cluster_4

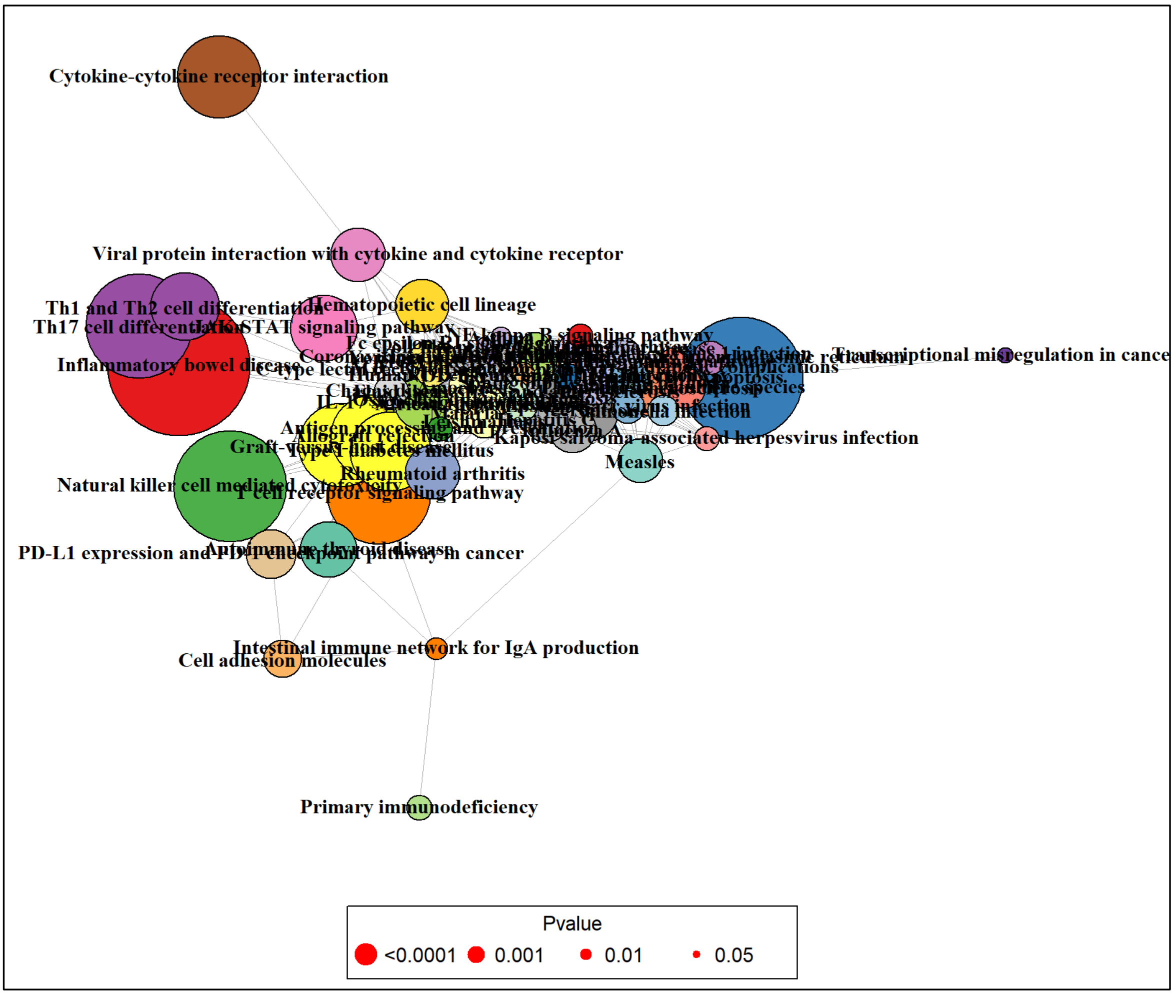

### Cluster_5

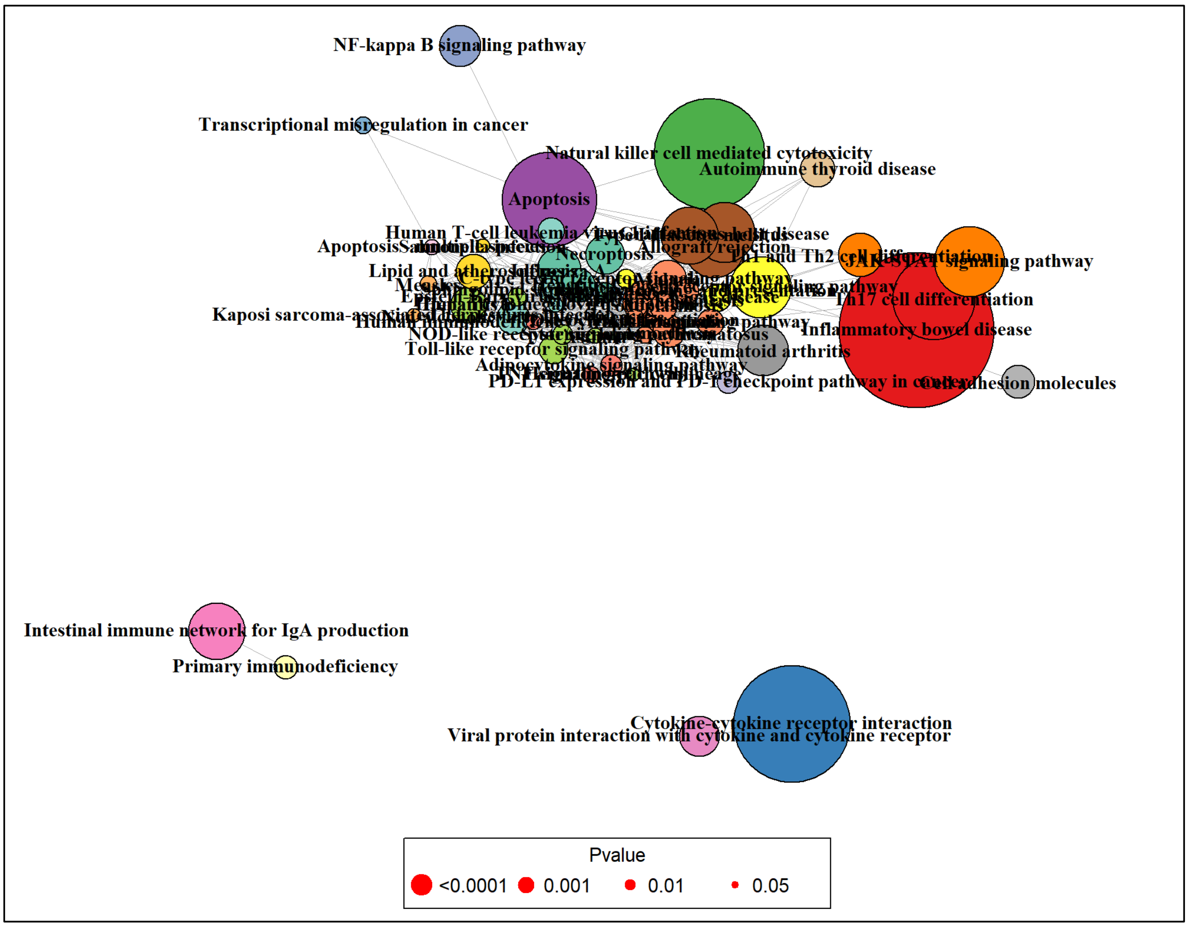


### Cluster_6

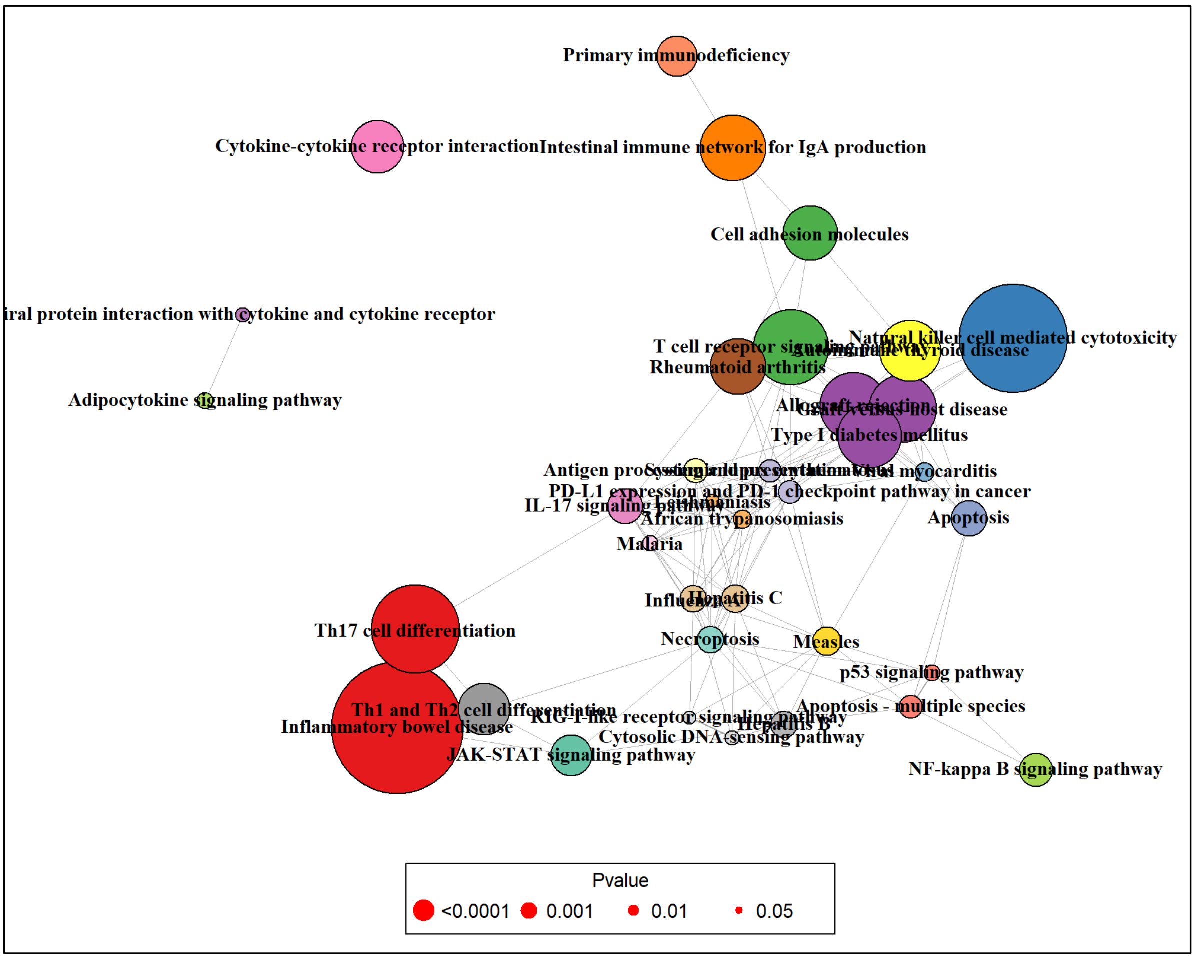

### Cluster_7

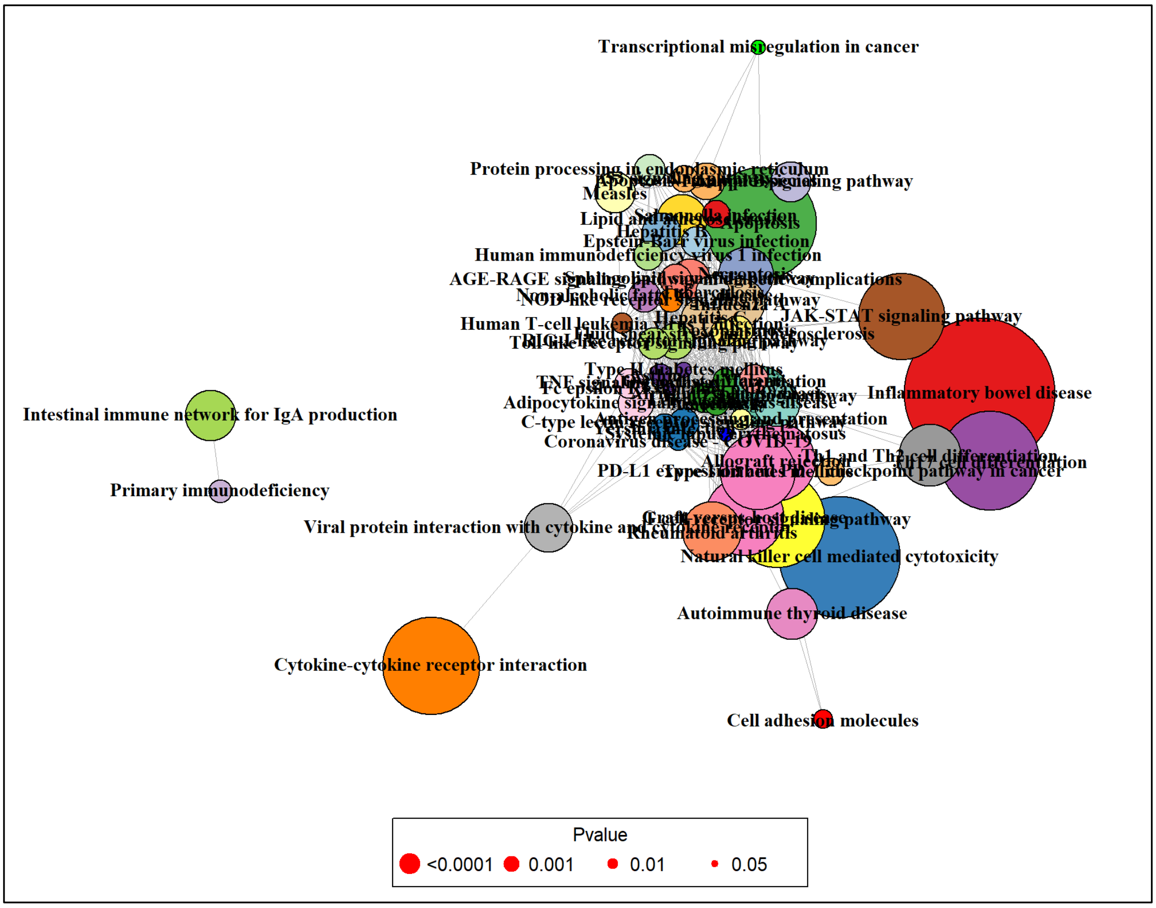


### Cluster_8

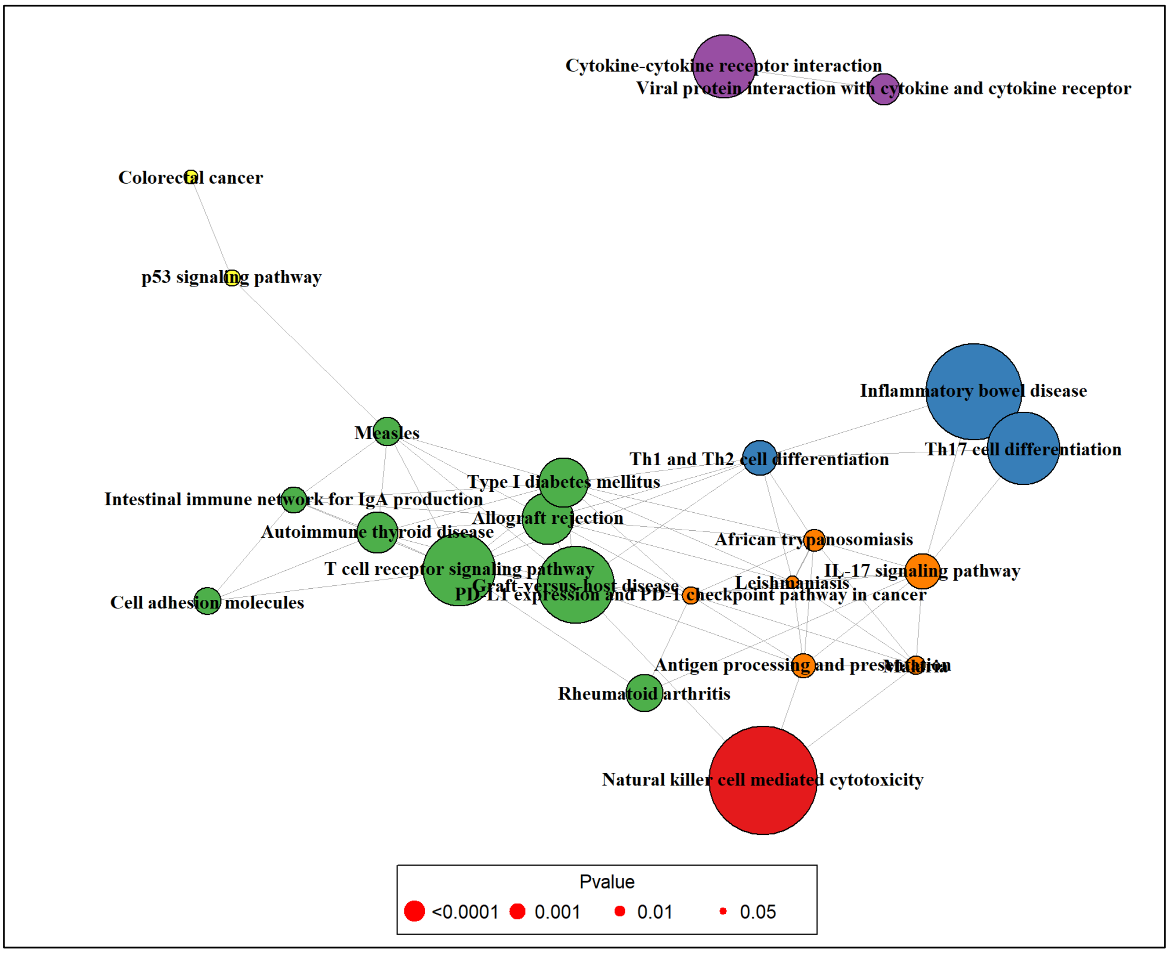

### Cluster_9

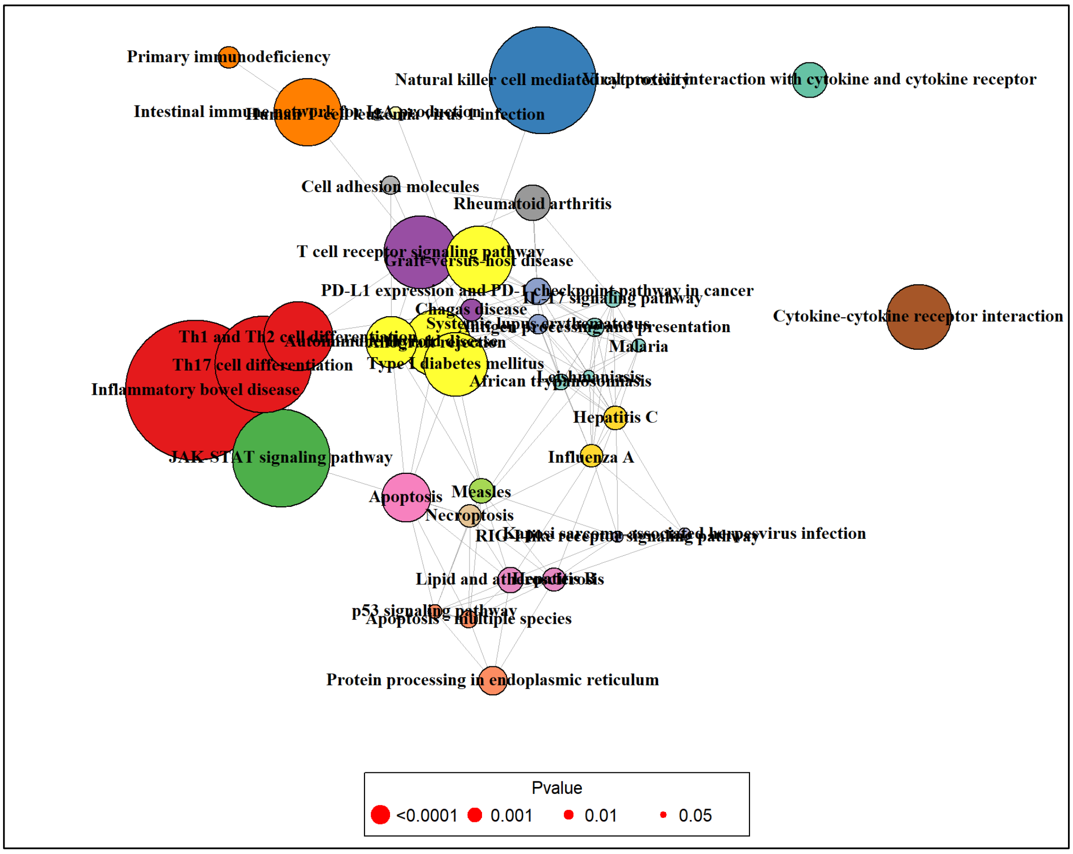


### Cluster_10

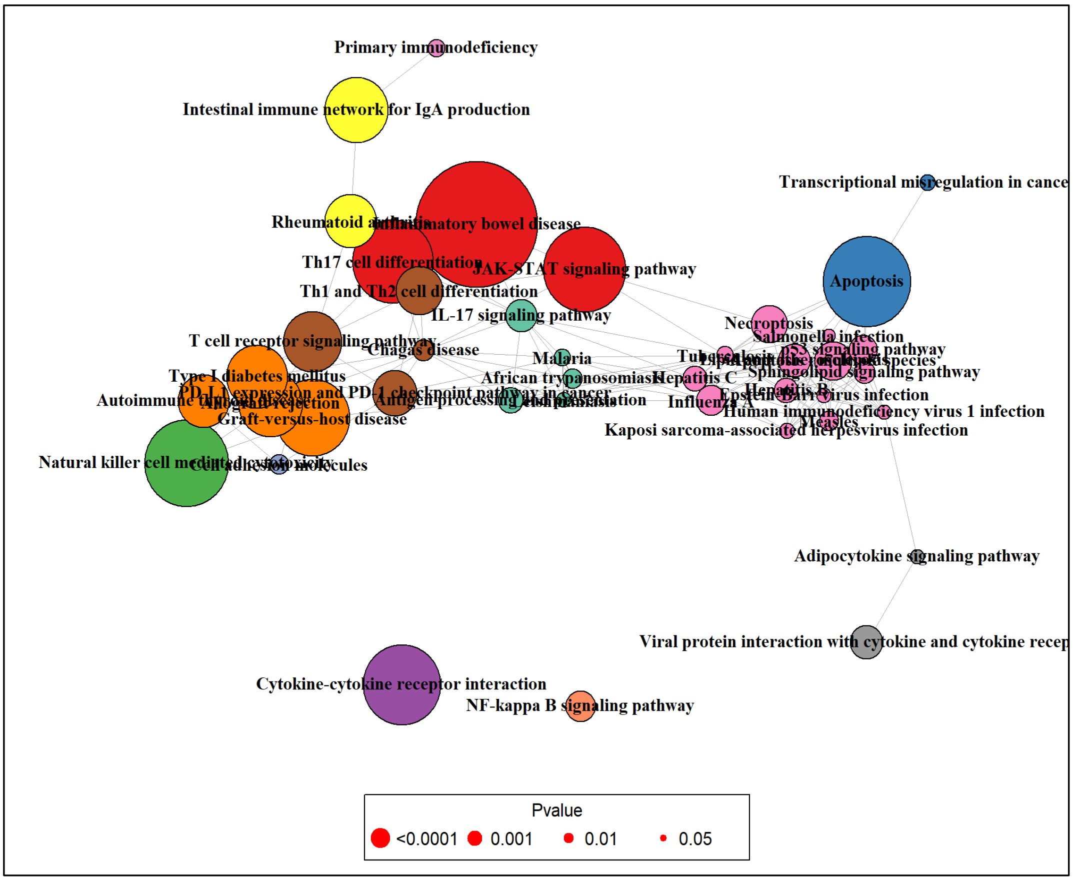

#### Cluster_11

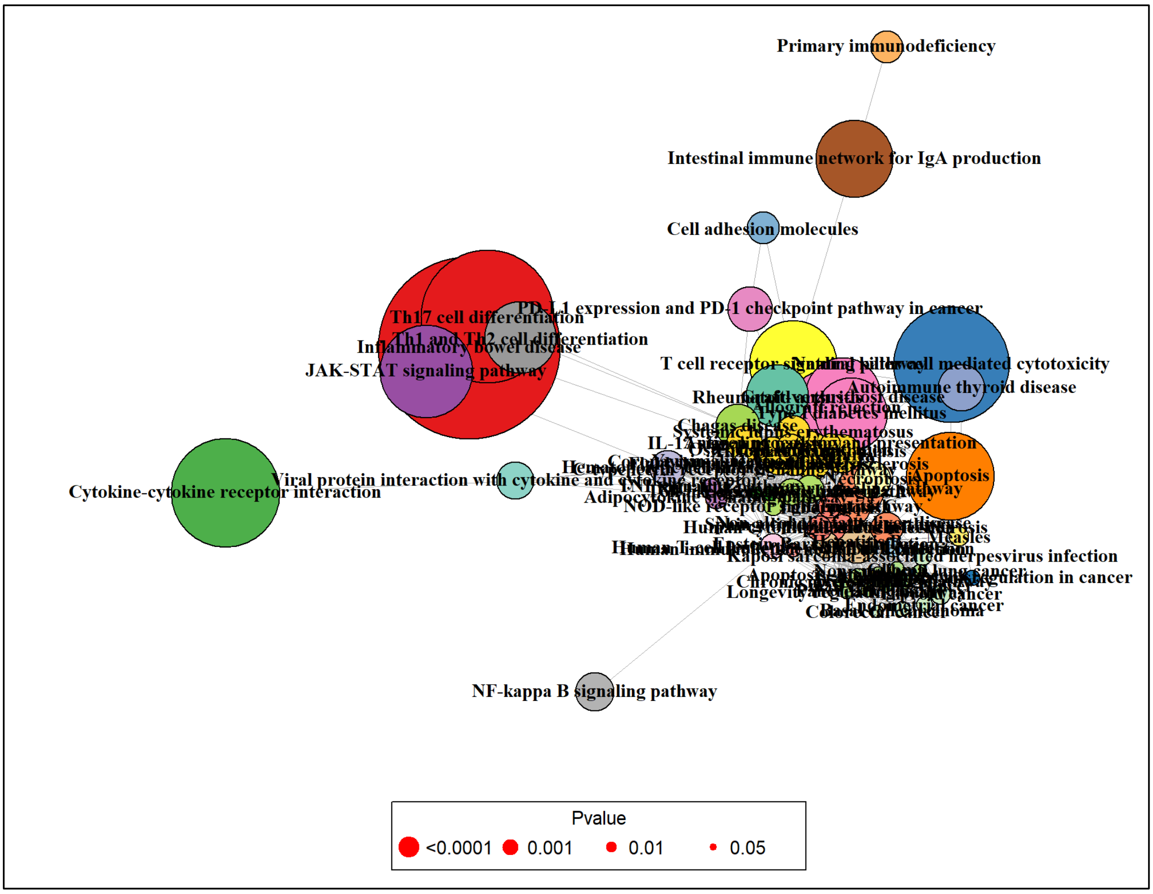


#### Cluster_12

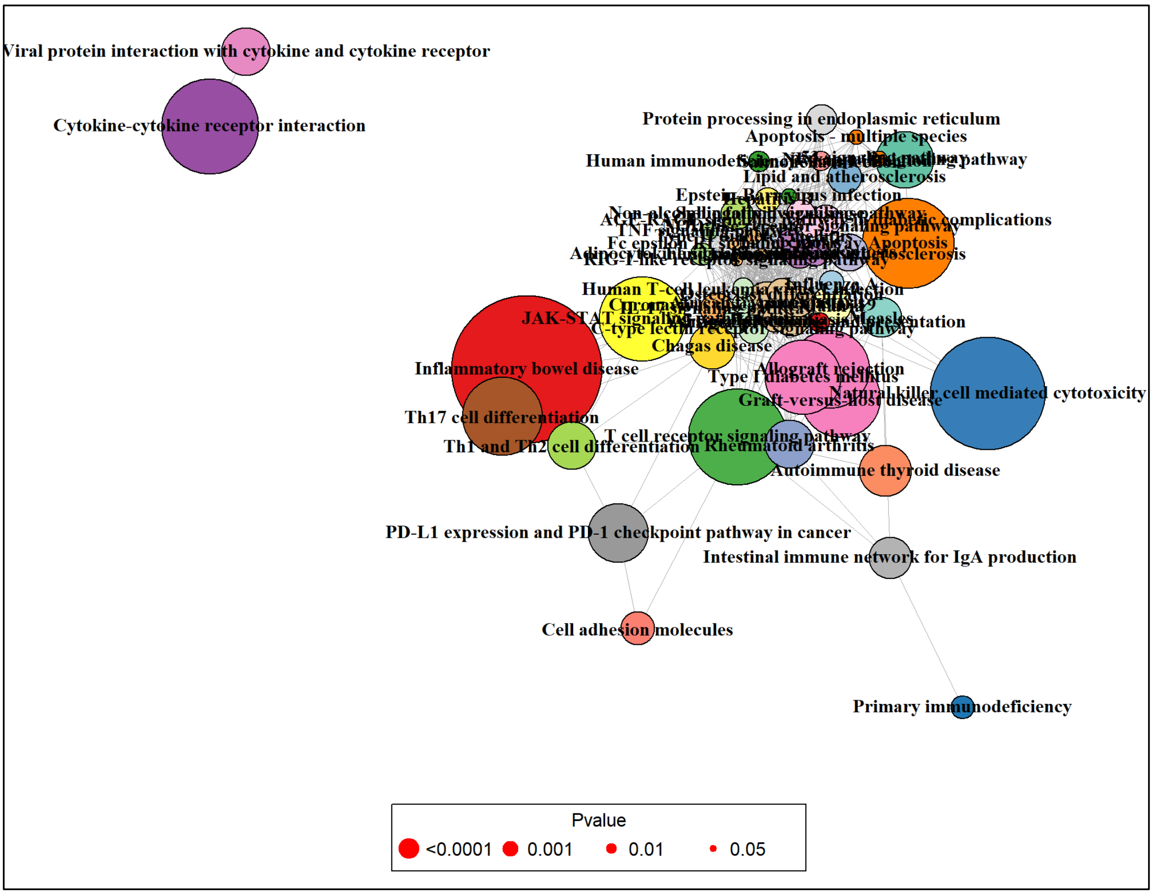

### Cluster_13

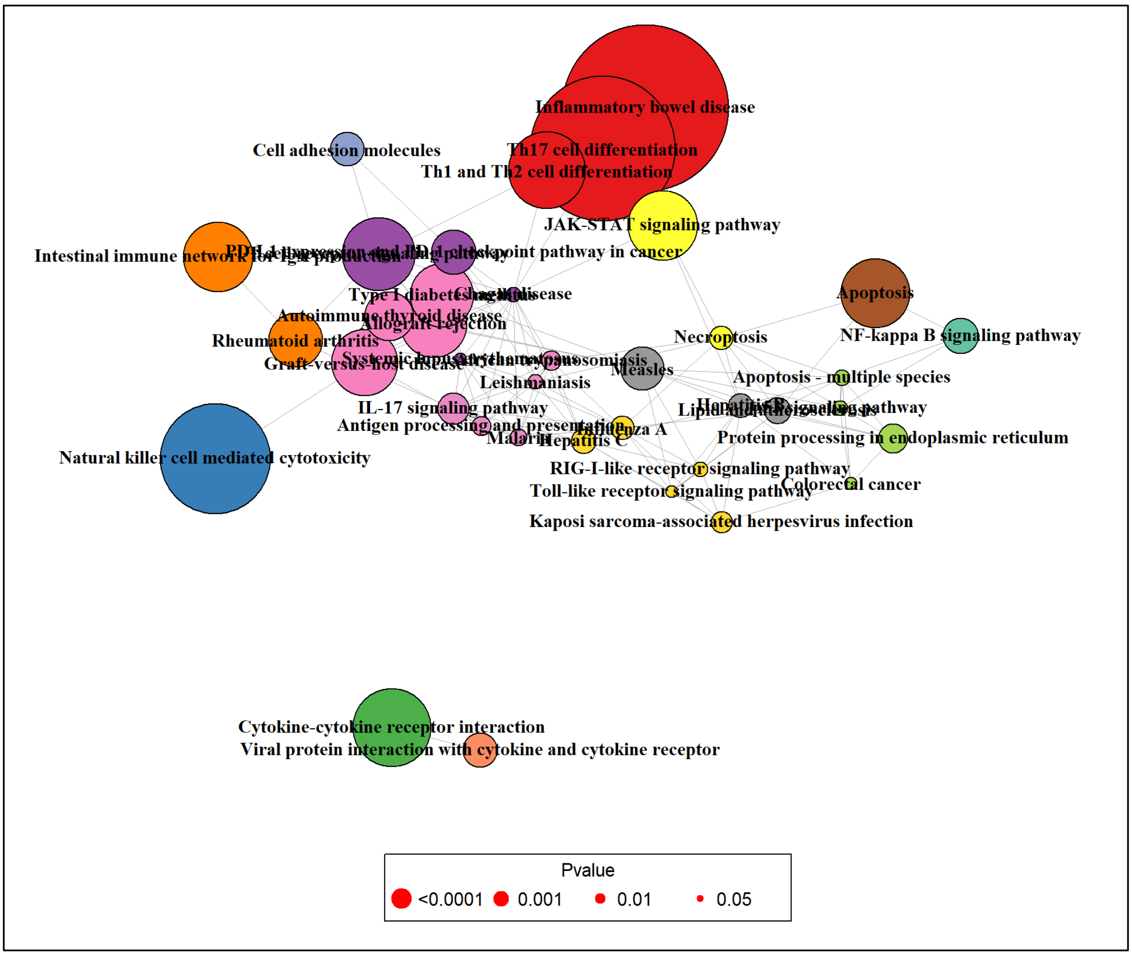
Inflammatory bowel disease
Cell adhesion molecules
Th17 cell differentiation
Th1 and Th2 cell differentiation
JAK-STAT signaling pathway
Apoptosis
NF-kappa B signaling pathway
Necroptosis
Autoimmune thyroid disease
Allograft rejection
Rheumatoid arthritis
Graft-versus-host disease
Measles
Leishmaniasis
Apoptosis - multiple species
IL-17 signaling pathway
Antigen processing and presentation
Malaria
Hepatitis C
Influenza A
Protein processing in endoplasmic reticulum
Natural killer cell mediated cytotoxicity
RIG-I-like receptor signaling pathway
Toll-like receptor signaling pathway
Colorectal cancer
Kaposi sarcoma-associated herpesvirus infection
Cytokine-cytokine receptor interaction
Viral protein interaction with cytokine and cytokine receptor
Pvalue
<0.0001
0.001
0.01
0.05
