## Supplemental Figure 3 for "Programmed cell death-1 receptor mediated regulation of Tbet^+^ NK1.1^−^ Innate Lymphoid Cells within the Tumor Microenvironment"

### Functional Gene Enrichment Analysis of Clusters

#### Cluster_1

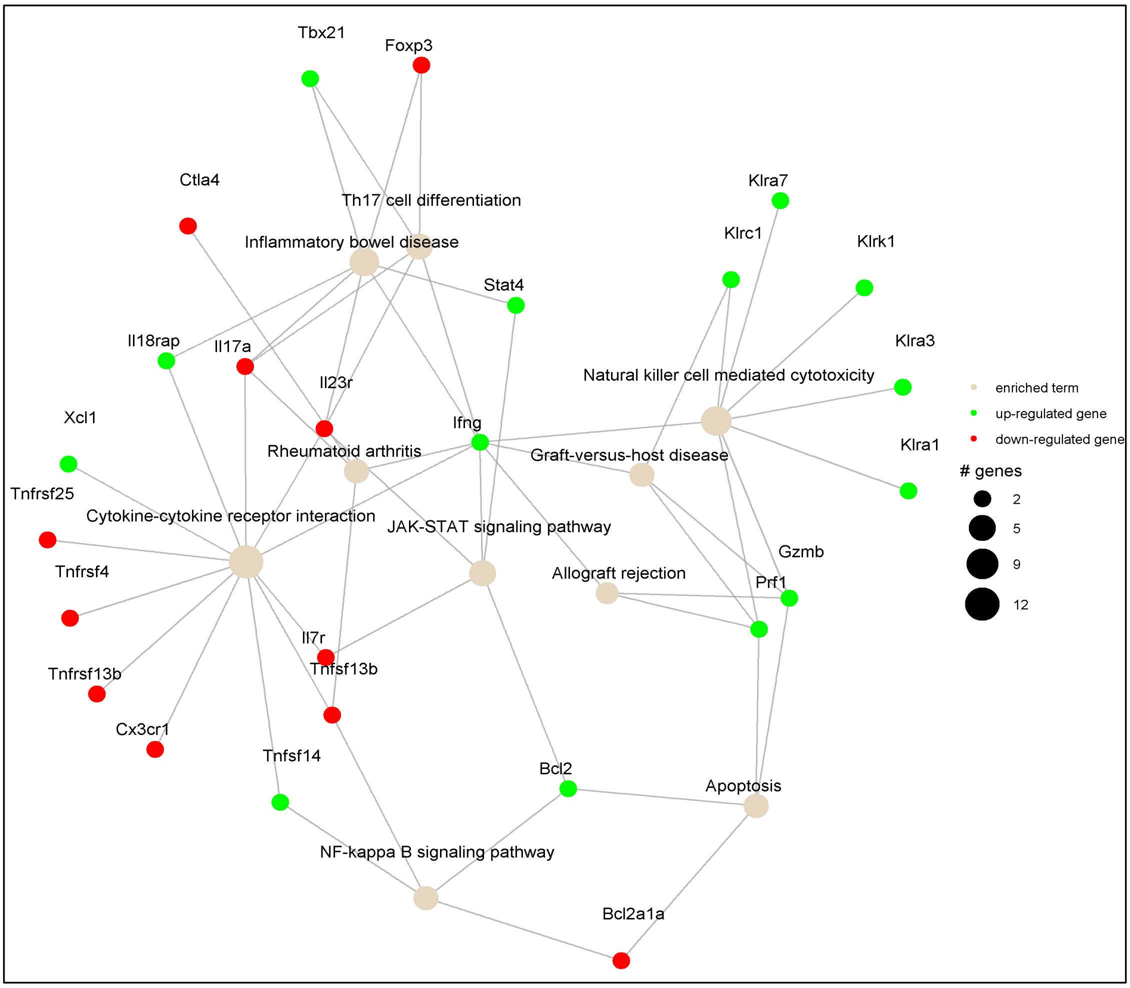


#### Cluster_2

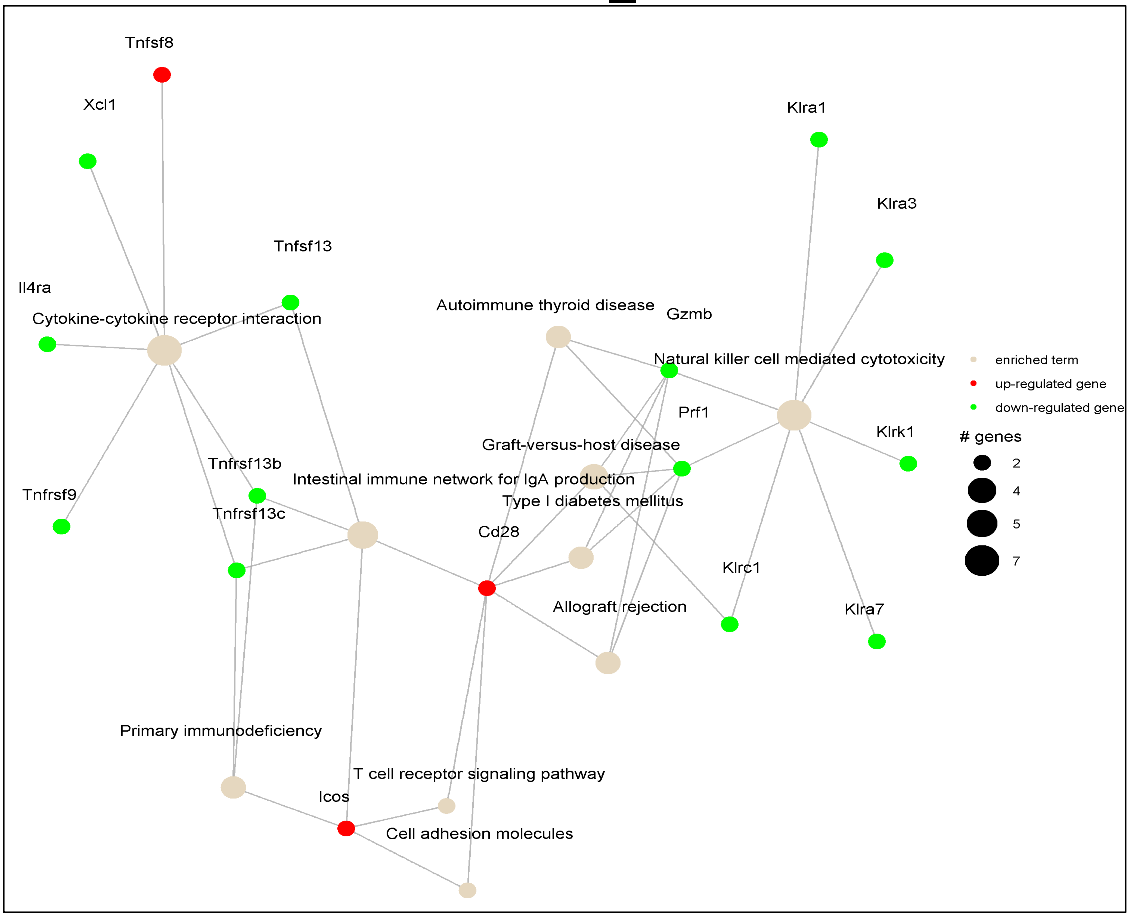

#### Cluster_3

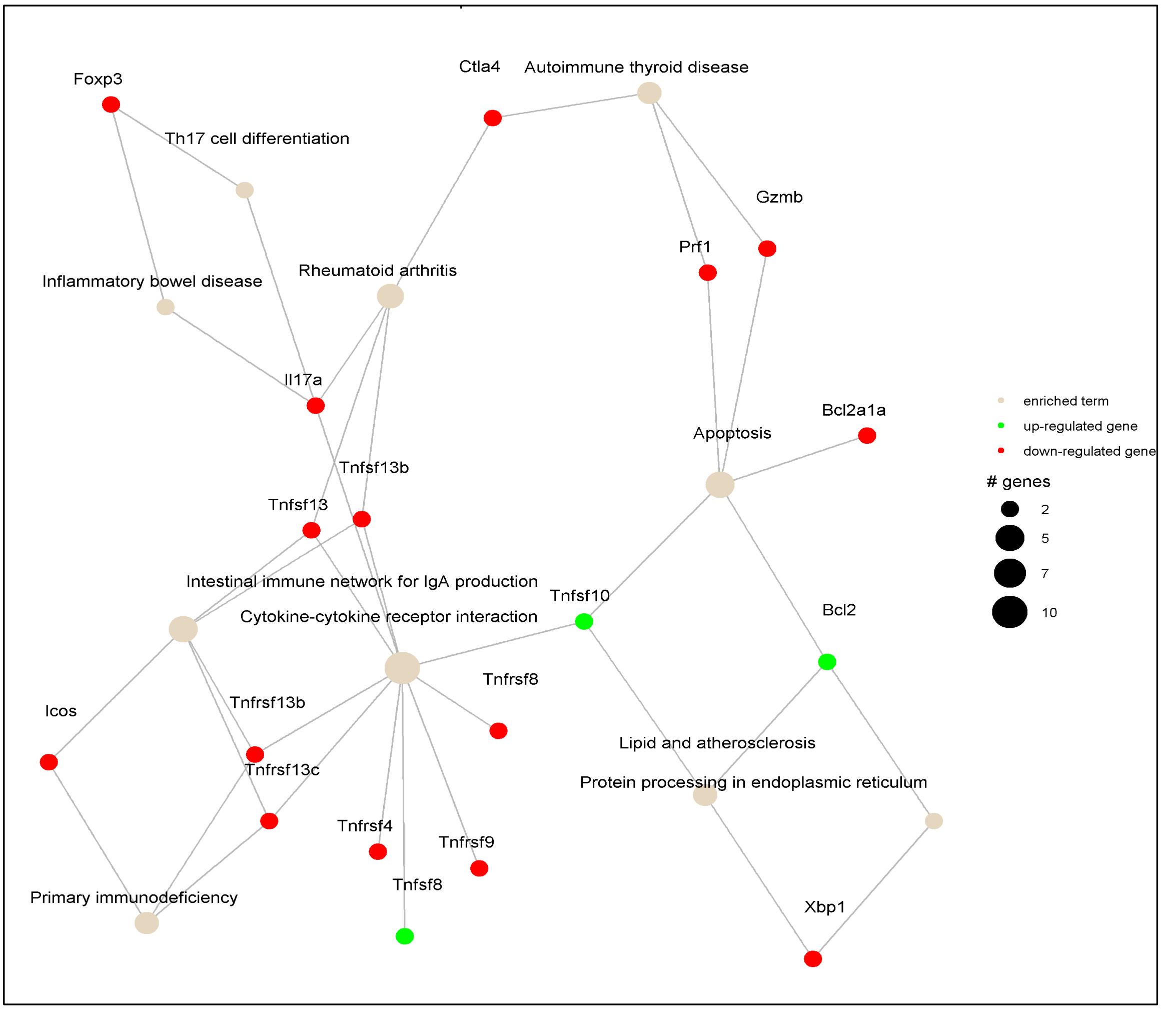


#### Cluster_4

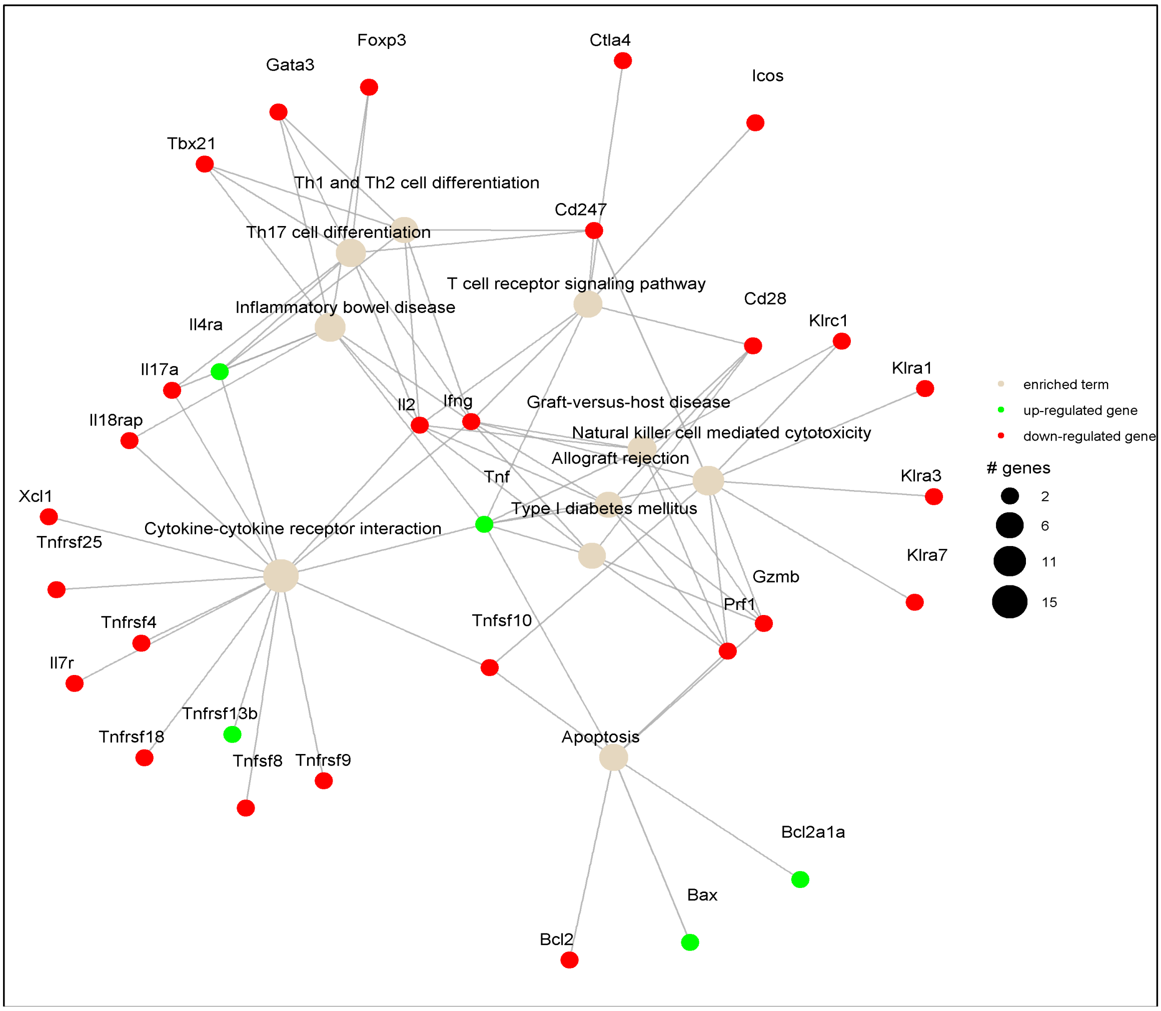

### Cluster_5

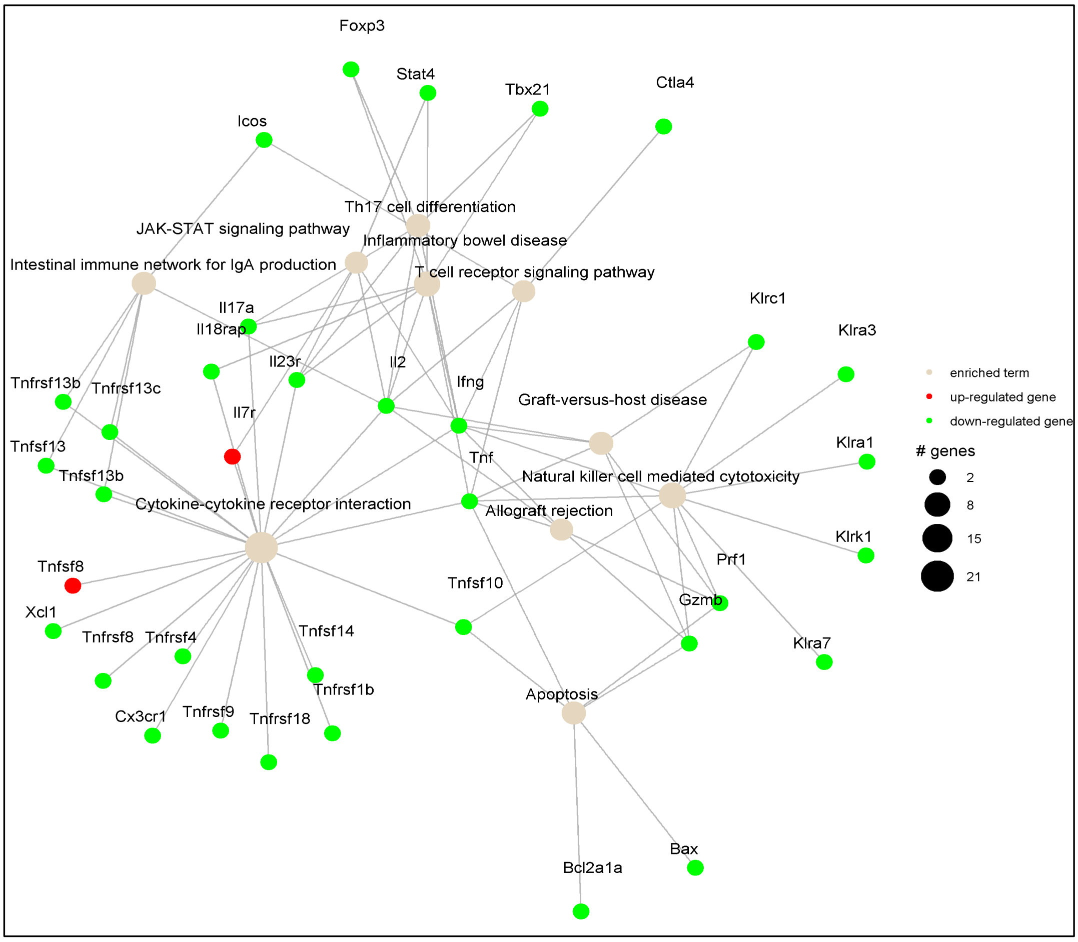


### Cluster_6

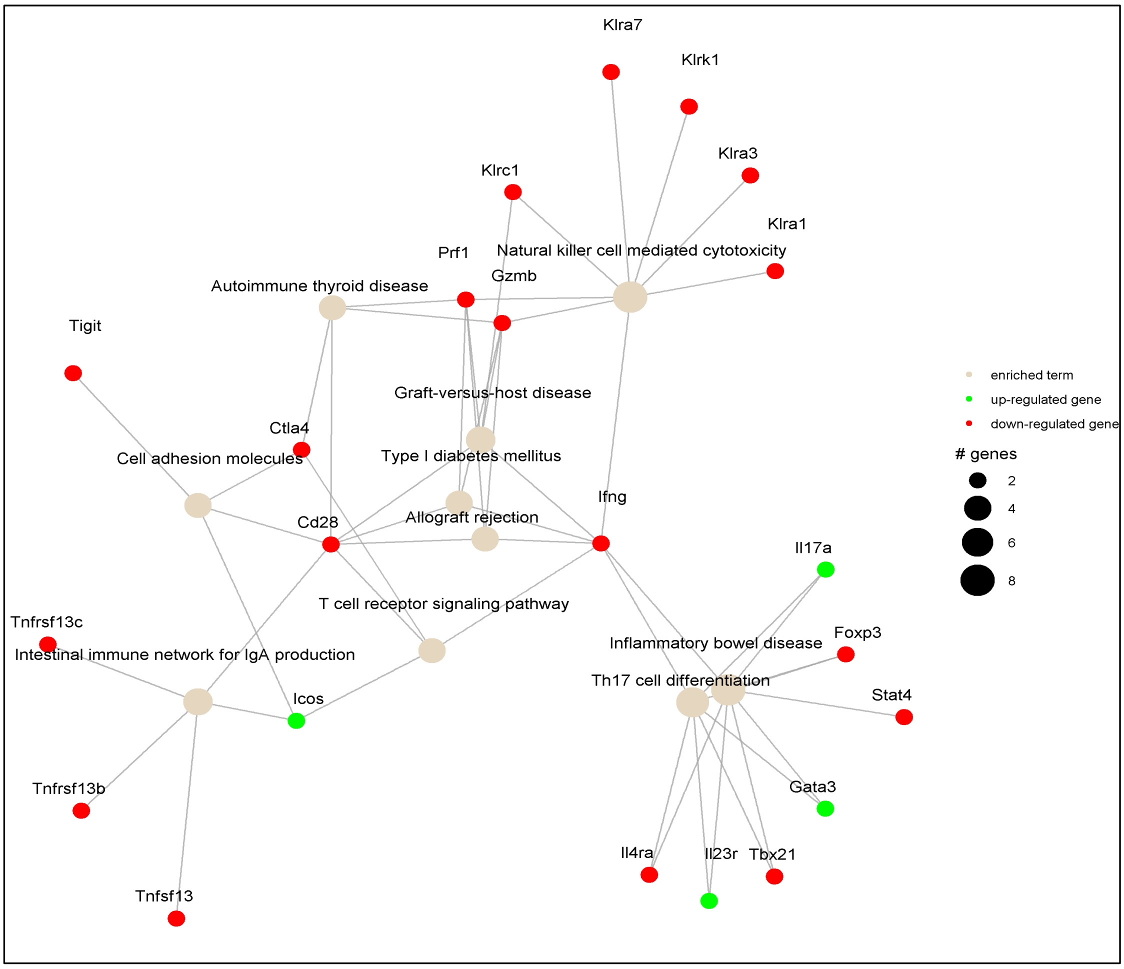

### Cluster_7

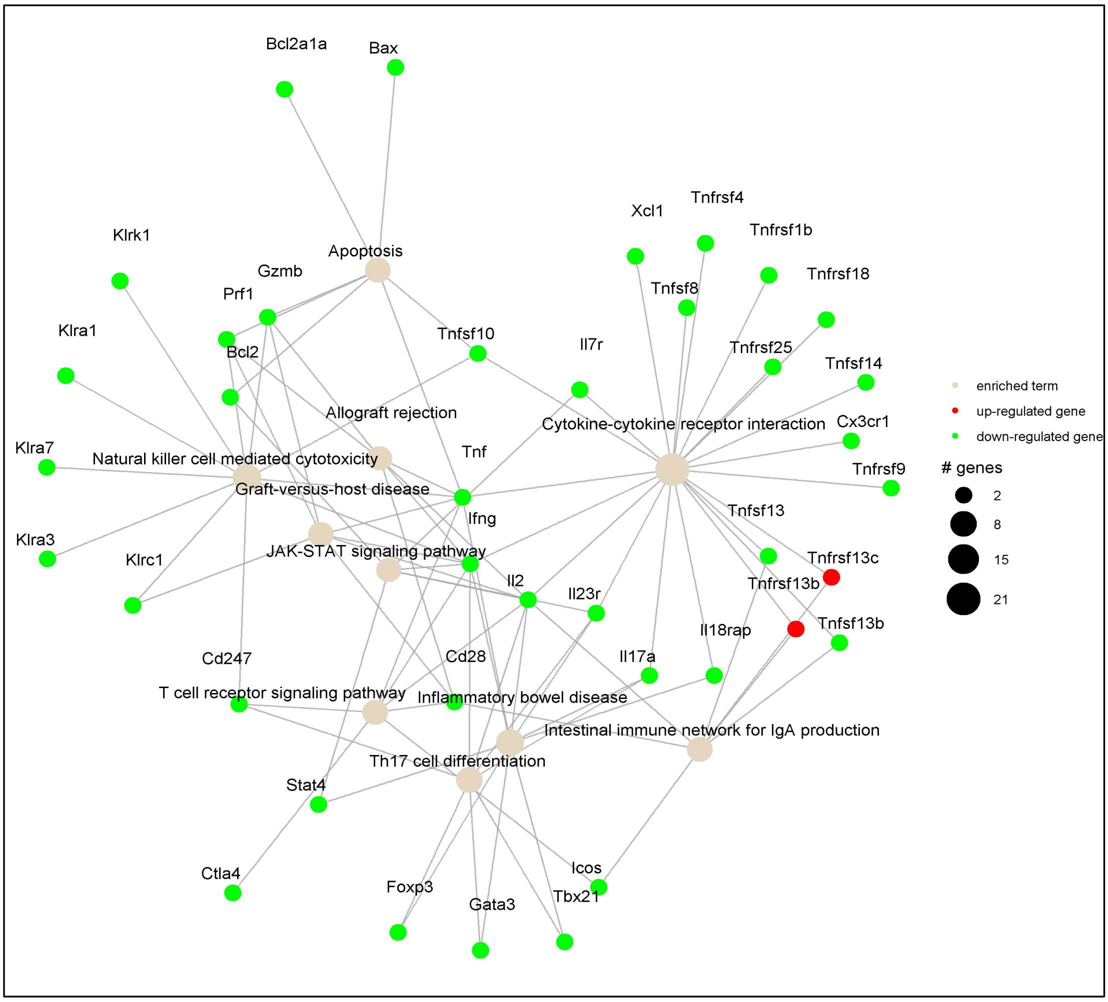


### Cluster_8

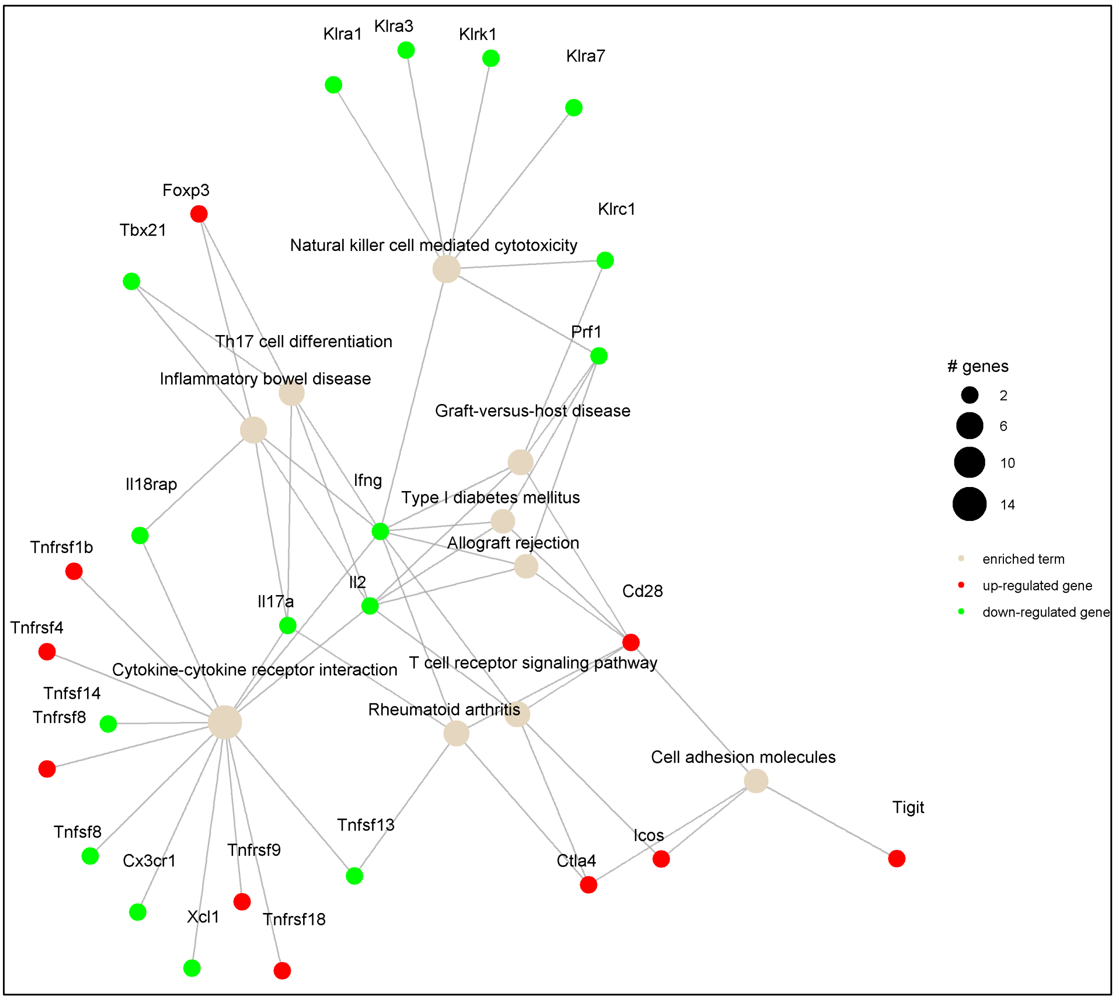

### Cluster_9

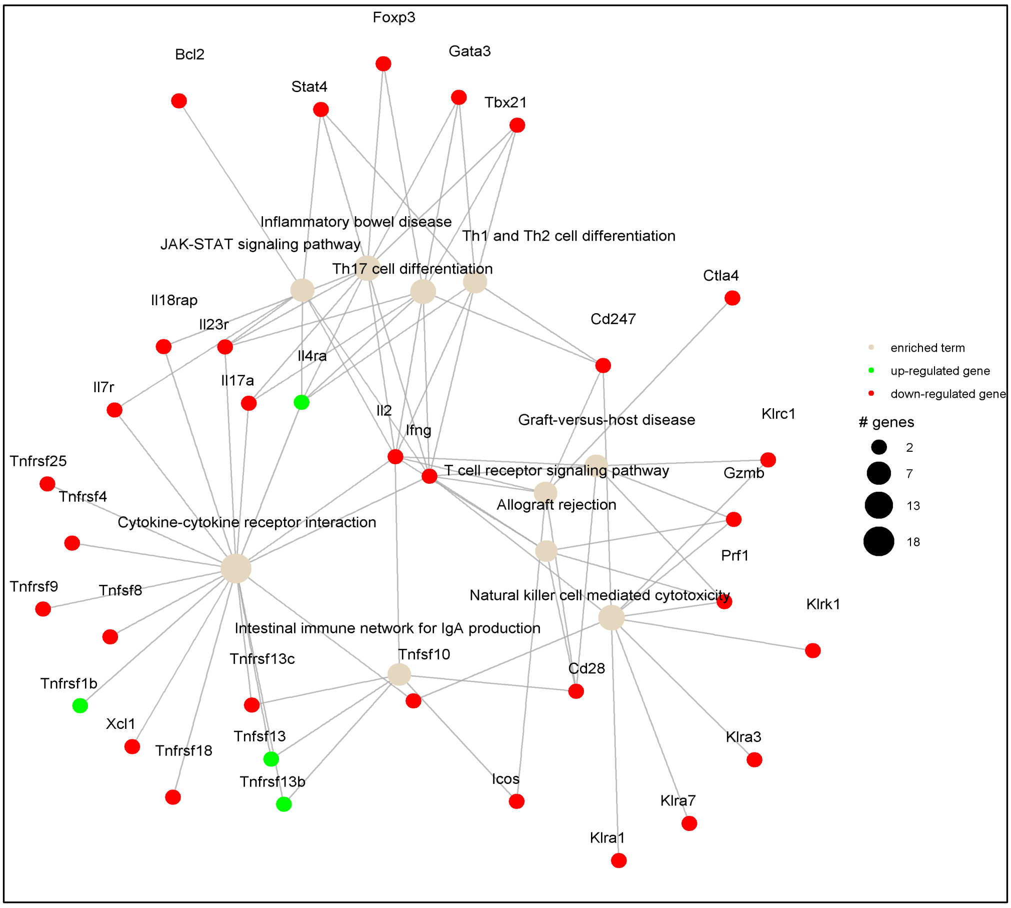


### Cluster_10

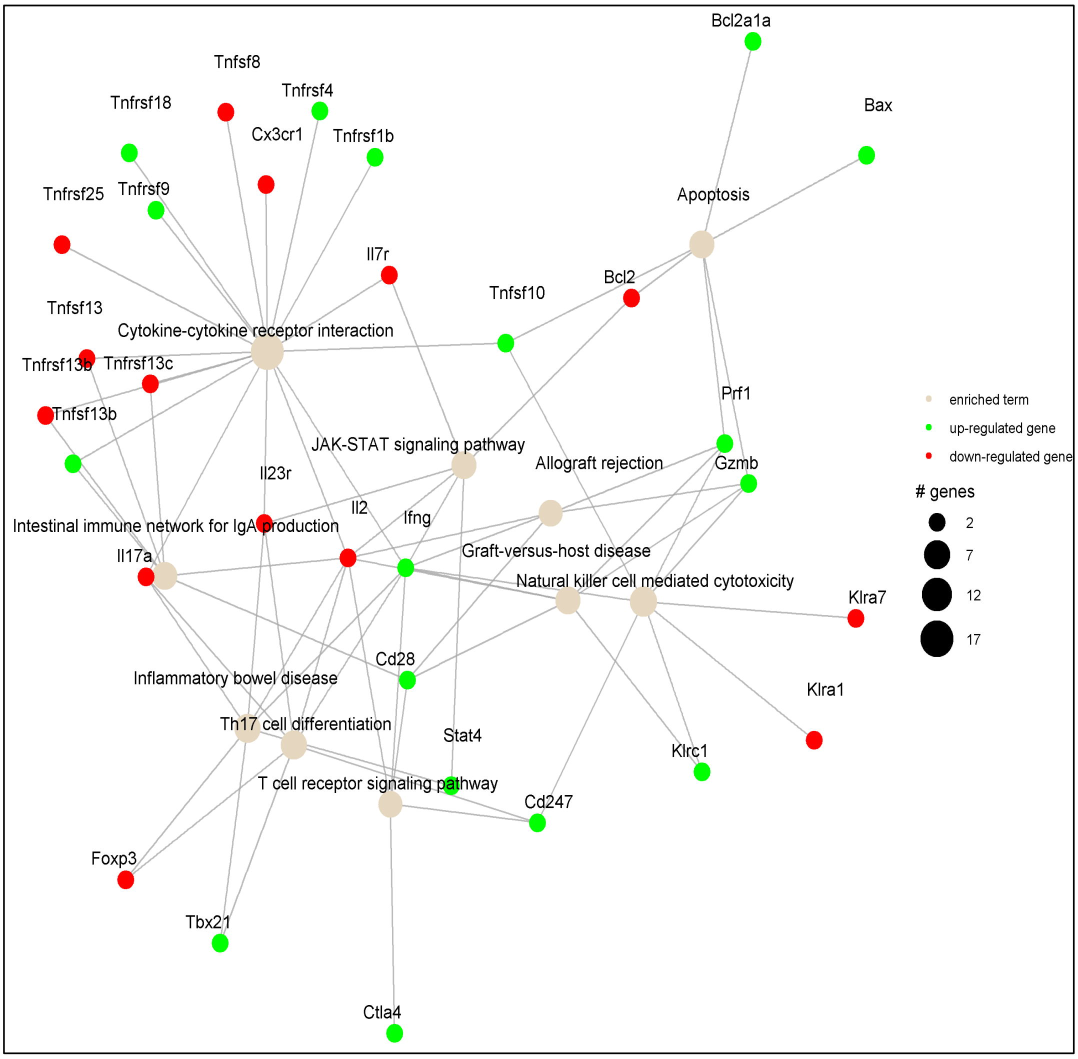

### Cluster_11

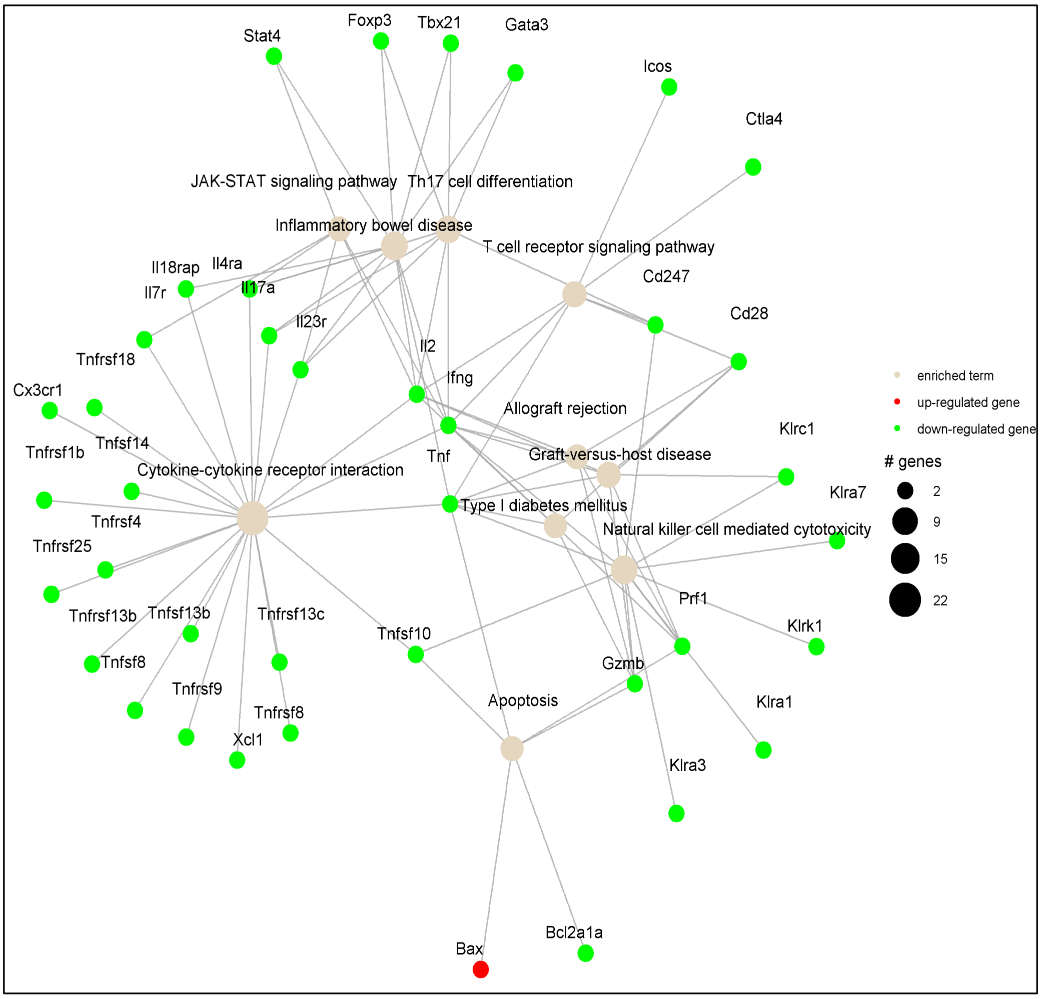


### Cluster_12

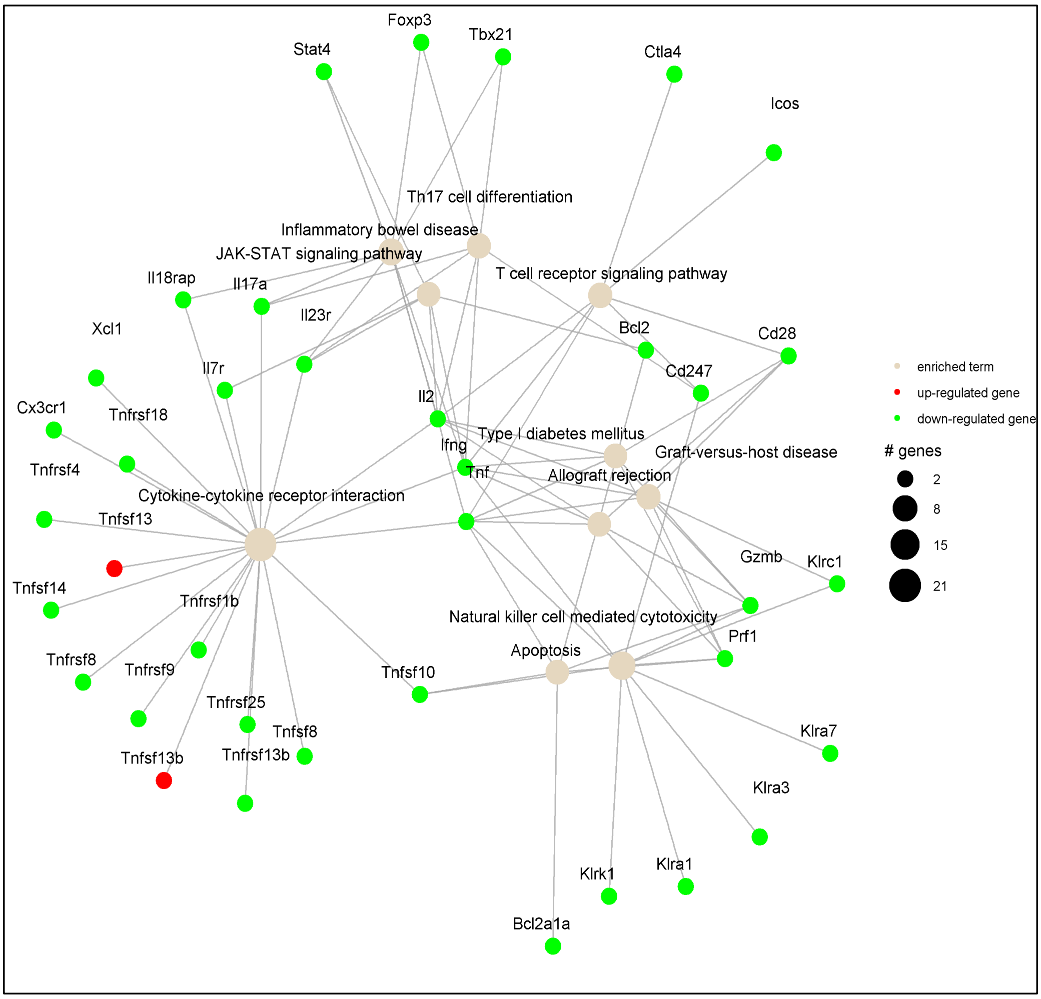

### Cluster_13

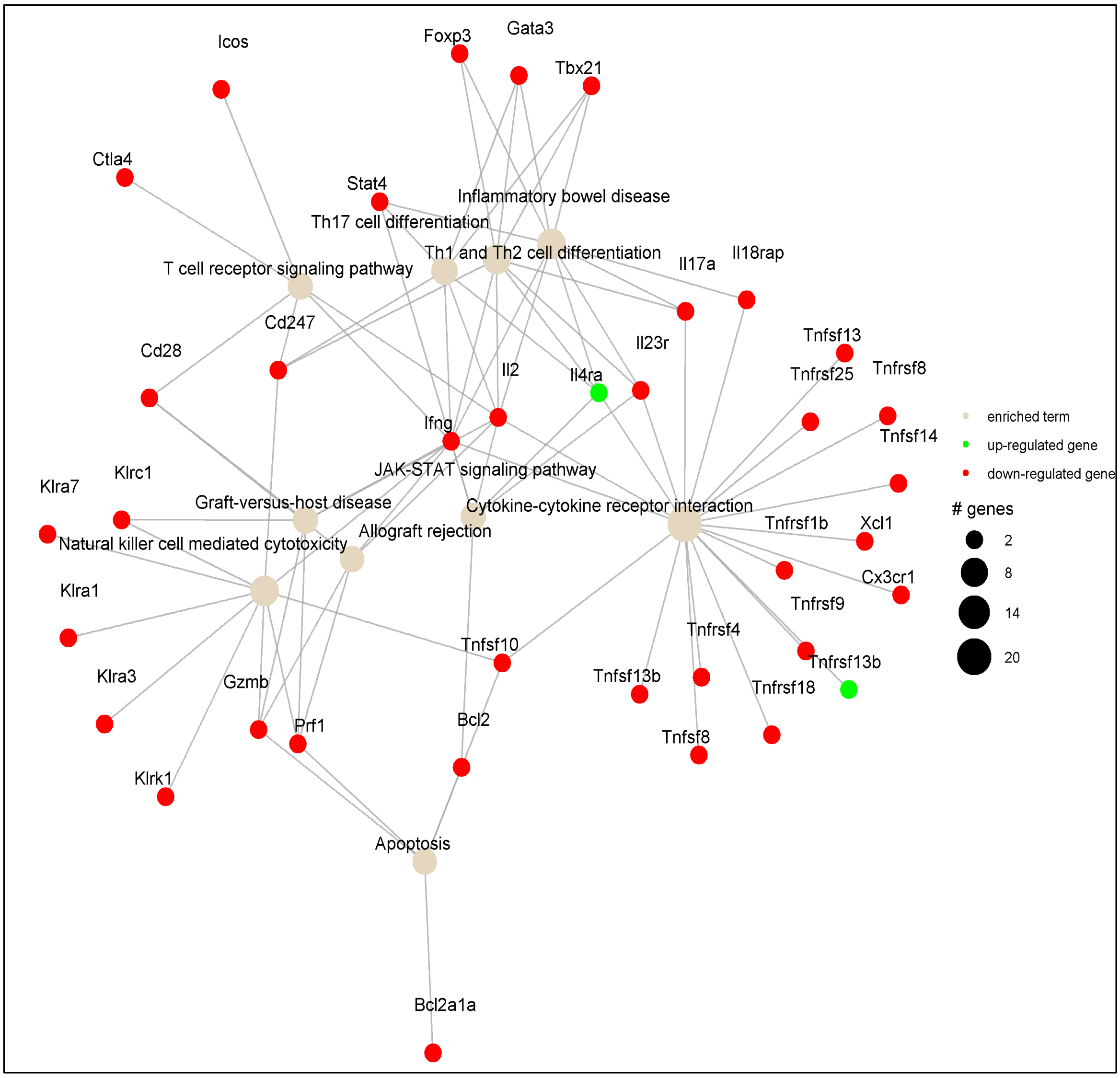

Icos
Foxp3
Gata3
Tbx21
Ctla4
Stat4
Inflammatory bowel disease
Th17 cell differentiation
Th1 and Th2 cell differentiation
Il17a
Il18rap
T cell receptor signaling pathway
Cd247
Cd28
Il23r
Tnfsf13
Tnfrsf25
Tnfrsf8
Il2
Il4ra
Ifng
Tnfsf14
JAK-STAT signaling pathway
Klrc1
Klra7
Graft-versus-host disease
Cytokine-cytokine receptor interaction
Tnfrsf1b
Xcl1
Natural killer cell mediated cytotoxicity
Allograft rejection
Cx3cr1
Klra1
Tnfrsf9
Tnfrsf4
Tnfsf10
Tnfrsf13b
Tnfsf13b
Klra3
Gzmb
Tnfrsf18
Prf1
Bcl2
Tnfsf8
Klrk1
Apoptosis
Bcl2a1a
enriched term
up-regulated gene
down-regulated gene
### genes
2
8
14
20
